# Single cell analyses of the effects of Amyloid-beta42 and Interleukin-4 on neural stem/progenitor cell plasticity in adult zebrafish brain

**DOI:** 10.1101/437467

**Authors:** Mehmet Ilyas Cosacak, Prabesh Bhattarai, Yixin Zhang, Caghan Kizil

## Abstract

Neural stem cells (NSCs) constitute the reservoir for new cells and might be harnessed for stem cell-based regenerative therapies. Zebrafish has remarkable ability to regenerate its brain by inducing NSC plasticity upon Alzheimer’s pathology. We recently identified that NSCs enhance their proliferation and neurogenic outcome in an Amyloid-beta42-based (Aβ42) experimental Alzheimer’s disease model in zebrafish brain and Interleukin-4 (IL4) is a critical molecule for inducing NSC proliferation in AD conditions. However, the mechanisms by which Aβ42 and IL4 affect NSCs remained unknown. Using single cell transcriptomics, we determined distinct subtypes of NSCs and neurons in adult zebrafish brain, identified differentially expressed genes after Aβ42 and IL4 treatments, analyzed the gene ontology and pathways that are affected by Aβ42 and IL4, and investigated how cell-cell communication is altered through secreted molecules and their receptors. Our results constitute the most extensive resource in the Alzheimer’s disease model of adult zebrafish brain, are likely to provide unique insights into how Aβ42/IL4 affects NSC plasticity and yield in novel drug targets for mobilizing neural stem cells for endogenous neuro-regeneration.

## Introduction

Zebrafish, with its extensive regenerative ability, has become a key model organism for studies on how tissues heal and regenerate (Alunni and Bally-Cuif, 2016; Kizil et al., 2012b; Zupanc, 2008). Although zebrafish provides unique opportunities to molecularly dissect the regenerative programs in many tissues, there are still hurdles that render the interpretation of complex cellular interactions and responses difficult, such as the heterogeneity of precursor cell populations and their differential response to stimuli (Marz et al., 2010).

Central nervous system regeneration is catching particular attention because of robust neural regeneration ability in zebrafish and clinical ramifications it might bring, which could not be possible to elucidate with mammalian models. For instance, we have recently generated an Amyloid toxicity model in adult zebrafish brain and identified that zebrafish can effectively enhance its neural stem cell proliferation and neurogenesis with key activity of Interleukin-4 (IL4) that mediates the crosstalk between disease pathology in neurons and initiation of regenerative output in stem cells (Bhattarai et al., 2017a; Bhattarai et al., 2016b; Bhattarai et al., 2017b; Kizil, 2018). We also found that IL4 can directly affect the human neural stem cells in a similar fashion to induce the proliferative and neurogenic ability (Papadimitriou et al., 2018). However, due to the heterogeneity of stem cell populations and neuronal subtypes in vertebrate brains, it was not possible to identify how Aβ42 and IL4 lead to enhanced stem cell plasticity and neurogenesis, which individual subtypes of stem cells and neurons responded to Aβ42 and IL4, and how individual responses were. Therefore, we performed a single cell sequencing in control, Aβ42-treated and IL4-treated adult zebrafish telencephalon and categorized the cell type identities, molecular programs of individual cell types, and how Aβ42 and IL4 alters those programs. Our results add further elaboration to our previous findings that Aβ42 and IL4 affect neural stem cells to enhance their neurogenic capacity by providing detailed analyses on heterogeneous cell populations. We are specifically providing cell type-specific information on the signaling pathways and cell-cell interactions through secreted molecules that are either lost or emerged after Aβ42 and IL4. We believe that our extensive datasets for cell type identification, gene expression, cell-cell interaction maps, GO-term analyses and determination of the transcription factor codes would provide a unique resource for detailed examination of adult zebrafish brain and its remarkable regenerative ability in homeostatic and neurodegenerative context.

## Results and Discussion

### Overall strategy for unbiased clustering of telencephalic cells of the adult zebrafish brain

To investigate the cell types of adult zebrafish telencephalon and their molecular properties, we designed an analysis pipeline (Figure 1A). We dissected telencephalon region of the adult brain of a transgenic zebrafish that expressed GFP under the *her4.1* promoter, which marks the neural stem/progenitor cells (NSPCs) with glial identity (Yeo et al., 2007). Using flow cytometry-assisted cell sorting, we removed the cell debris and the dead cells (Figure S1) and enriched the GFP+ and GFP− cells. To perform single cell sequencing, we mixed viable GFP+ and GFP− cells in a 1:1 ratio. After single cell sequencing, we mapped the resulting reads to zebrafish genome and performed unbiased clustering (Figure 1B-E). We determined the genes that are enriched in certain clusters using Seurat software (Figure 1F,G; Figure S2, Figure S3 Dataset S1A). By using literature-based cell type identifiers (Dataset S1B) and our “marker” genes, we ascertained cell type identities to the clusters (Figure 1H, I; Figure 2). To further investigate the cell types, we performed GO-term enrichment and pathways analyses (genes/pathways enriched in one cell cluster compared to other clusters) (Figure 3A-G; Dataset S1C-H), determined the transcription factor (TF) codes for individual cell clusters (which TFs are expressed in individual clusters and what is the similarity level of TF expression to other cell clusters) (Figure 3H-K; Dataset S1I), and generated an interaction map by analyzing the known secreted molecules and their cognate receptors (putative communication between different cell clusters using secreted molecules and their receptors based on expression patterns in cell clusters) (Figure 3L-M; Dataset S1J-L). We also used this pipeline for zebrafish brains treated with Amyloid-beta42 (Aβ42) and Interleukin-4 (IL4). We determined the differentially expressed genes in every cell cluster after Aβ42 (Figure 4; Dataset S2A-K) and IL4 (Figure 5; Dataset S3A-L). Our results that will be explained in subsequent sections provide detailed classification of NSPCs in zebrafish, give important insights into molecular programs of individual cell types, identify the molecular programs affected by Aβ42 and IL-4, and thereby explain the molecular basis of how neural stem cells enhance their plasticity and funnel into a regenerative state in Alzheimer’s disease conditions. Additionally, our data supports our previous findings that IL4 is a critical cue for NSPC plasticity (Bhattarai et al., 2016b; Papadimitriou et al., 2018) and furthers adds to our molecular understanding on how IL4 activates the NSPCs in zebrafish brain.

**Figure 1:**
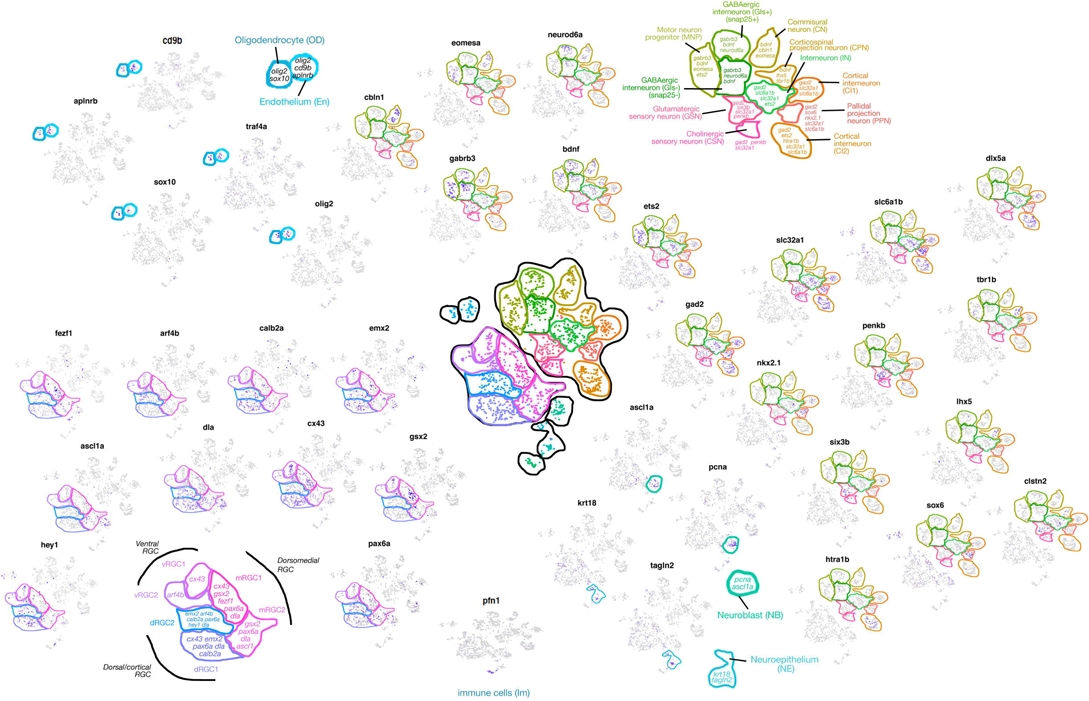
Cell sorting and categorization of cell types. (A) Overall strategy for cell sorting and single cell data analyses. (B) tSNE clustering. (C) Co-existence map of cells from control and Aβ42-treated brains on tSNE plots. (D) Identity distribution map. (E) Correlation strength plot. (F) Heat map of marker genes in identified clusters. (G) Dot plot for marker genes in identified clusters. (H) Feature plots for marker genes for glia, neurons, immune cells, neuroepithelium, and oligodendrocytes. (I) Schematic representation of cell type categorization on tSNE plot. See Figures S1-3, Dataset S1A.

**Figure 2:**
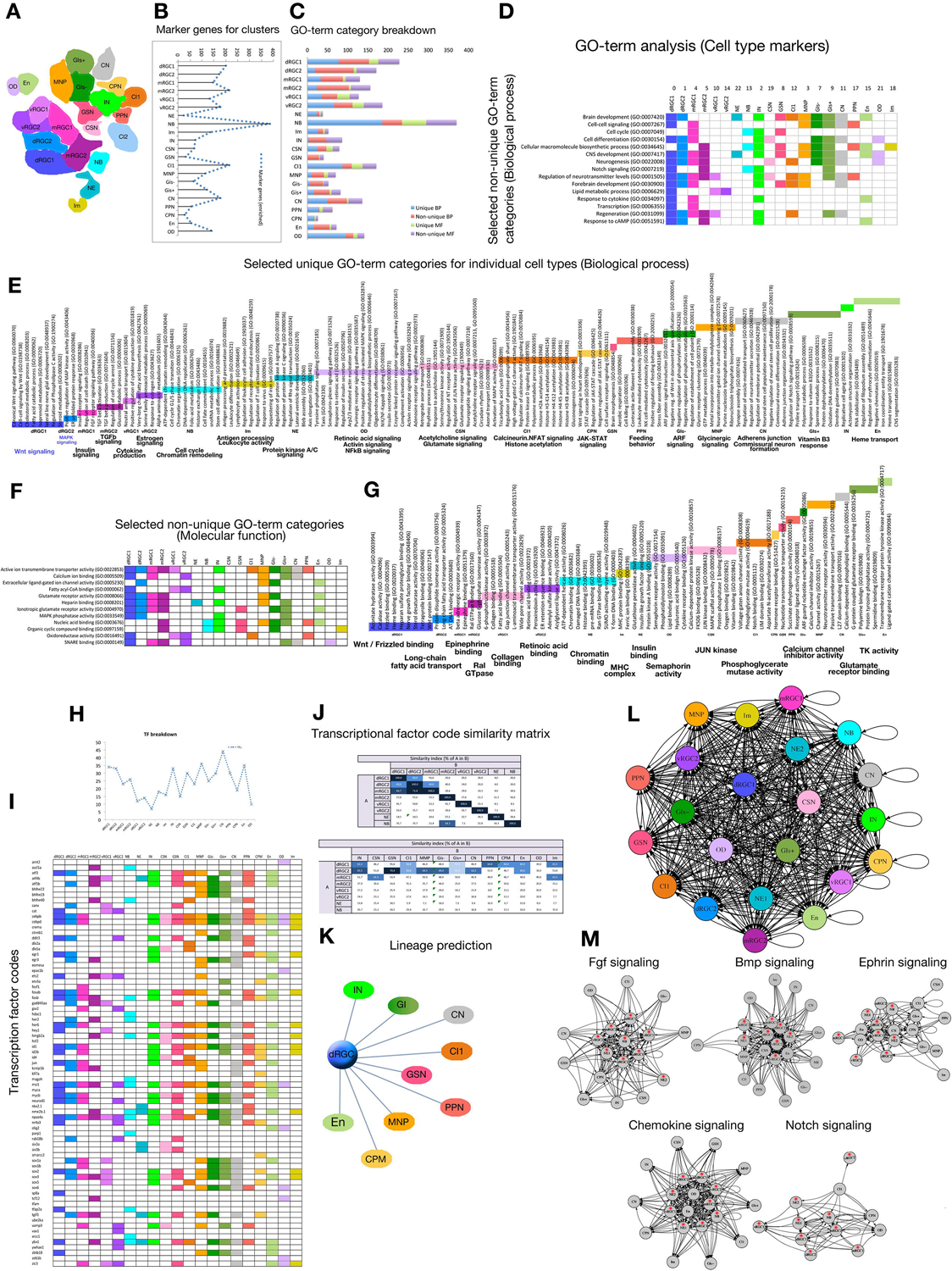
Identification of cell types. Schematic representation of feature plots for individual cell types. Genes that mark specific cell types are indicated. See Figure S3, Dataset S1B.

**Figure 3:**
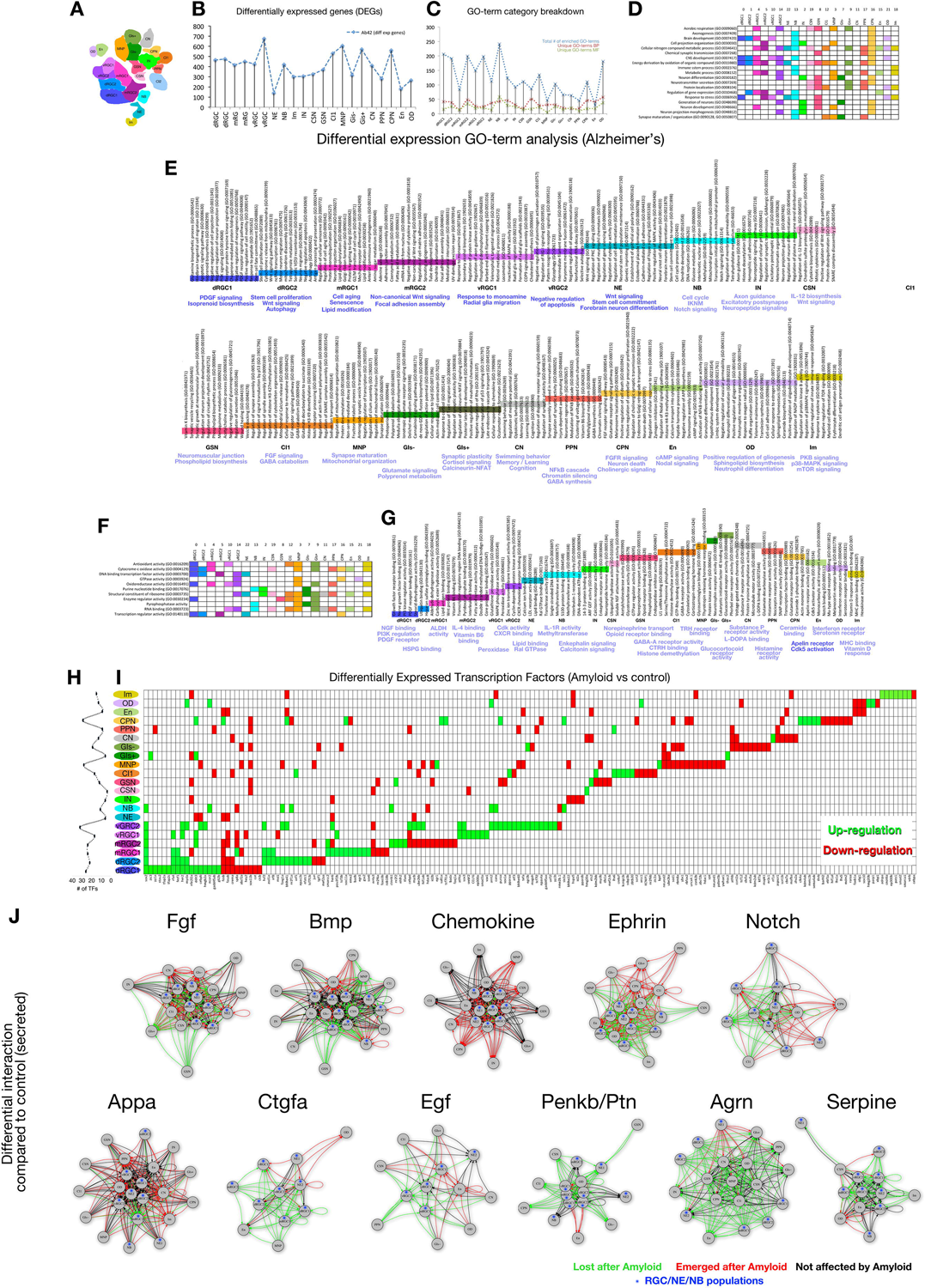
Analyses of cell types in control brains. (A) Colored scheme for individual cell types. (B) Chart showing the distribution of marker genes for clusters. (C) Breakdown of identified GO-term categories for individual cell types. (D) Selection of non-unique biological process (BP) GO-term categories. (E) Selected unique BP GO-term categories. (F) Selection of non-unique molecular function (MF) GO-term categories. (G) Selected unique MF GO-term categories. (H) Chart showing the number of transcription factors (TFs) expressed in individual cell types. (I) Chart showing individual TFs in all cell types. (J) TF similarity matrices. (K) Lineage prediction chart based on TF expression similarity. (L) Complete interaction map based on secreted molecules and their receptors. (M) Selection of interaction maps for Fgf, Bmp, Ephrin, Chemokine, and Notch signaling. See Dataset S1C-L.

**Figure 4:**
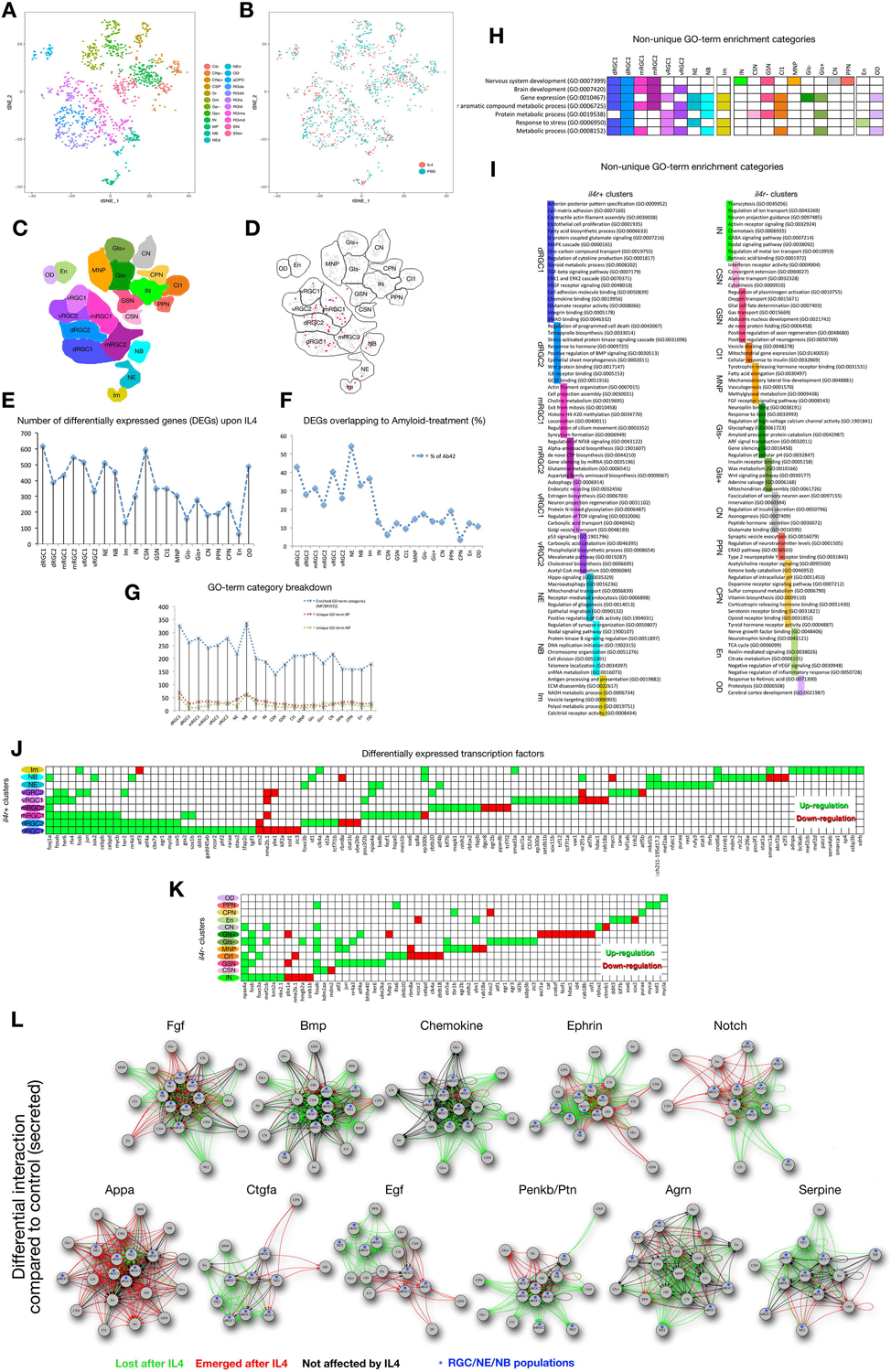
Analyses of cell types in Aβ42-treated brains. (A) Colored scheme for individual cell types. (B) Chart showing the distribution of differentially expressed genes in individual clusters. (C) Breakdown of identified GO-term categories for individual cell types. (D) Selection of non-unique biological process (BP) GO-term categories. (E) Selected unique BP GO-term categories. (F) Selection of non-unique molecular function (MF) GO-term categories. (G) Selected unique MF GO-term categories. (H) Line chart showing the number of differentially expressed transcription factors (TFs) in individual cell types after Aβ42. (I) Chart showing all differentially expressed transcription factors (TFs) in individual cell types. (J) Selected interaction maps based on differentially expressed secreted molecules and receptors. Arrowheads indicate the presence of the receptor in a certain cell type. Green lines: interactions that are lost upon Aβ42, red lines: newly formed interactions after Aβ42, black lines: interaction is both present in control and Aβ42-treated brains. Blue dots represent the neurogenic populations. See Dataset S2A-K.

**Figure 5:**
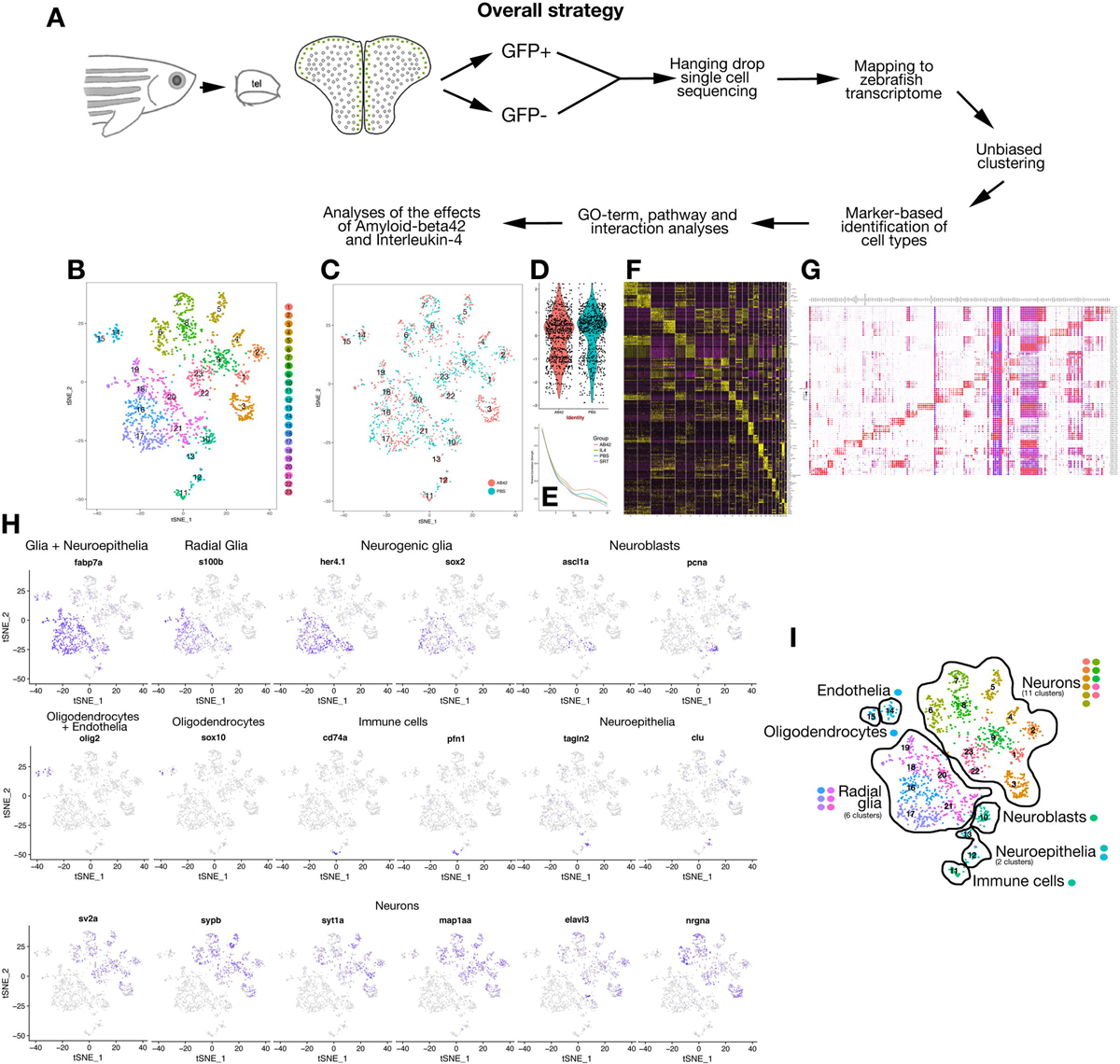
Analyses of cell types in Interleukin-4-treated brains. (A) tSNE plot for cell clusters. (B) Co-existence map of cells from control and IL4-treated brains on tSNE plots. (C) Colored scheme for individual cell types. (D) Feature plot for expression of interleukin-4 receptor gene *il4r*. Red cells are positive for *il4r*. (E) Chart showing the distribution of differentially expressed genes in individual clusters. (F) Chart showing the percentage of differentially expressed genes that overlap in IL4-treated and Aβ42-treated brains per individual cell types. (G) Breakdown of identified GO-term categories for individual cell types. (H) Selection of non-unique GO-term categories (MF+BP). (I) Selected unique GO-term categories (MF+BP). (J) Chart showing all differentially expressed transcription factors (TFs) in *il4r+* cell types. (K) Chart showing all differentially expressed transcription factors (TFs) in *il4r−* cell types. (L) Selected interaction maps based on differentially expressed secreted molecules and receptors in IL4-treated brains. Arrowheads indicate the presence of the receptor in a certain cell type. Green lines: interactions that are lost upon IL4, red lines: newly formed interactions after IL4, black lines: interaction is both present in control and IL-treated brains. Blue dots represent the neurogenic populations. See Dataset S3A-L.

### Cell clustering and identification of cell types

After unbiased clustering of cells and tSNE analyses, we identified 23 cell clusters (Figure 1B). In order to determine whether the clustering is reliable, we co-mapped cells from control brains and Aβ42-injected brains (Figure 1C). If the markers for cell clusters that are identified in an unbiased way were affected by treatment of Aβ42 and therefore would not be used as cell type markers, we would see non-overlapping control and Aβ42-treated cell clusters. However, we found that cells from control and Aβ42 conditions co-segregate into same cell clusters in all clusters with the exception of cluster 3, which was predominantly occupied by Aβ42-treated cells (Figure 1C) indicating that the unbiased use of the markers for defining cell clusters is reliable. Identity plot (Figure 1D) and shared component analysis (Figure 1E) showed that clustering is statistically reliable and acceptable. Using the Seurat software (Butler et al., 2018), we determined the marker genes for every cell cluster (Dataset S1A). When we performed unbiased heat-map (Figure 1F, Figure S2) and dot-plot (Figure 1G) for cluster-specific genes identified by the default settings of Seurat package, we found that every cluster contains cell-type marker genes that can be used to identify the clusters. In order to predict a gross categorization of cells in our clustering, we used the following widely-used marker combinations: radial glia (*fabp7a+*, *s100b+*, *her4.1+*) (Ito et al., 2010; Raponi et al., 2007; Yeo et al., 2007), neurogenic glia (*her4.1+*, *sox2+*) (Ellis et al., 2004), neuroblasts (*ascl1a+*, *pcna+*) (Kizil et al., 2012b; Kizil et al., 2012c; Marz et al., 2010), neuropeithelium (*fabp7a+*, *tagln2+*, *clu+*) (Lake et al., 2016), oligodendrocytes (*olig2+*, *sox10+*) (Morrens et al., 2012), endothelial cells (*olig2+*, *sox10−*) (Klein-Soyer et al., 2000), immune cells (*cd74a+*, *pfn1+*) (Fan et al., 2014), and neurons (*sv2a+*, *sypb+*, *syt1a+*, *map1aa+*, *elavl3+*, *nrgna+*) (Greif et al., 2013; Kwon and Chapman, 2011; Nowack et al., 2010) (Figure 1H, Figure S2). By drawing feature plots that show gene expression on individual cells (purple indicates the gene is expressed; Figure 1H, Figure S3), we concluded that our clustering can be subclassified into 7 cell types (Figure 1I): radial glia (6 clusters), neurons (11 clusters), neuroepithelia (2 clusters), oligodendrocytes (1 cluster), endothelia (1 cluster), neuroblasts (1 cluster), and immune cells (1 cluster).

### Further delineation of cell clusters

To determine the identity of cell clusters, we used a larger set of marker genes that are biologically verified to be characterizing certain cell types (Dataset S1B). Based on expression of these markers (feature plots in Figure 2 and Figure S3), we named all 23 clusters (Figure 2; Dataset S1B). For instance, the radial glial cells that comprised of 6 clusters could be identified as dorsal (dRGC) (high *emx2*, *hey1*, *calb2a*, *dla* and low *fezf1*, *gsx2* expression) (Gangemi et al., 2001; Ganz et al., 2011; Grandbarbe et al., 2003; Than-Trong et al., 2018), medial (mRGC) (high *gsx2*, *fezf1*, *dla* and low *emx2*, *calb2a* expression) (Pei et al., 2011; Shimizu and Hibi, 2009; Xu et al., 2006), and ventral (vRGC) (high *gsx2*, *arf4b* and low *fezf1*, *emx2*, *dla*, *pax6a* expression) (Curto et al., 2014; Ezratty et al., 2016; Gangemi et al., 2001; Ganz et al., 2011; Heins et al., 2002) (Figure 2; Figure S3; Dataset S1B). Additionally, these three RGC types can be subdivided into two clusters; however, a definitive unbiased distinction cannot be made with the current data. Yet, we have seen some marker genes differentially expressed in only one of the clusters. For instance, radial glial markers *s100a10* (Milosevic et al., 2017) and *cx43* (Duval et al., 2002) are expressed in only one of the dRGC, mRGC and vRGC clusters (dRGC1, mRGC1, vRGC1; Dataset S1B). Since these genes are associated with a more progenitor state of the radial glia, we can speculate that *s100a10+/cx43+* clusters could represent a radial glial state that is more progenitor-like. This is consistent with the absence of immature neuronal fate marker elavl3 (*Kizil et al., 2012b*; *Tallafuss et al., 2015*) in dRGC1 and mRGC1 but its presence in dRGC2 and mRGC2 (Dataset S1B, Figure S3). Given that all RGC clusters express widely-used glial markers *s100b (Raponi et al., 2007)*, *her4*, *fabp7a (Ito et al., 2010)*, *gfap (Lam et al., 2009)*, *msi1 (Kaneko et al., 2000)*, *slc1a3b (Untiet et al., 2017)*, *glula (Grupp et al., 2010)*, *sox2 (Ellis et al., 2004)*, *vim (Doetsch et al., 1997)*, *hopx (Li et al., 2015)* and *notch3 (Alunni et al., 2013)* (Figure 2; Figure S3; Dataset S1B), our unbiased separation of RGCs into six clusters with differential expression of regional markers in important in terms of delineating the heterogeneity of different glial progenitors.

Marker analyses also helped us to suggest cellular identities to 11 neuronal clusters (Figure 2; Figure S3; Dataset S1B). For instance, 6 clusters expressed glutamatergic neuron marker *gad2* (Sassa et al., 2007) and with additional markers, these clusters could be identified as glutamatergic sensory neurons (GSN, *six3b+* and *penkb+*) (Appolloni et al., 2008; Paul et al., 2017), cholinergic sensory neuron (CSN, *slc32a1+* and *penkb+*) (Fattorini et al., 2015; Paul et al., 2017), pallidal projection neurons (PPN, *sox6+* and *nkx2.1+*) (Ganz et al., 2011), cortical interneurons (CI1 and CI2, *slc32a1+* and *slc6a1b+*) (Fattorini et al., 2015; Hasel et al., 2017), and other interneurons (IN, *ets2+*) (Lake et al., 2016). CI2 specifically expresses the serine protease *htra1b*; however, the functional significance of this expression is unknown as of yet. Two out of the remaining five neuronal clusters can be classified as GABAergic interneurons (GIs, *neurod6a+* and *gabrb3+*) (Butt et al., 2007; Lujan et al., 2005; Ohtsuka et al., 2011) and based on Parvalbumin expression can be separated into *pvalb7a+* GIs (GIs+) and *pvalb7a−* GIs (GIs−) (Figure 2; Dataset S1B). The remaining three clusters are comprised of commissural neurons (CN, *cbln1+* and *eomesa+*) (Cameron et al., 2012; Otsuka et al., 2016), corticospinal projection/Cajal-Retzius neurons (CPN, *lhx5+* and *tbr1b+*) (Bedogni et al., 2010; Miquelajauregui et al., 2010) and motor neuron progenitors/excitatory neuron progenitors (MNP, *eomesa+* and *ets2+*) (Cameron et al., 2012; Lake et al., 2016). We could also verify the initial clustering of cell types with further markers such as endothelium (*cd9b*, *aplnrb*) (Klein-Soyer et al., 2000; Reaux et al., 2001), neuroepithelium (*krt18*, *tagln2*) (Lee et al., 2014; Merrick et al., 1995; Pollen et al., 2015) and mixed immune cells (*pfn1* and *cd74*) (Fan et al., 2014; Gil-Yarom et al., 2017). These results indicate that the unbiased clustering of cells and marker analyses can provide 23 different cell types in adult zebrafish telencephalon. This information is important because for instance in many studies including our own, zebrafish NSPC plasticity was assessed by relying on double labeling of S100b/PCNA (S100b+/PCNA+ cells are active progenitors and S100b+/PCNA-cells are quiescent progenitors) (Baumgart et al., 2012; Bhattarai et al., 2016b; Kizil et al., 2012c; Marz et al., 2010). However, our results show that there are at least 6 cell types that are *s100b+/pcna−* and they express in varying combinations various progenitor cell markers such as *ascl1a*, *neurod1*, *notch1b* and *hey1* (Figure 2; Figure S3; Dataset S1B). This indicates that those cells types constitute the heterogeneous NSPC populations that might be subjected to different regulatory molecular pathways. Elucidation of individual NSPC types is also important as it can allow us to identify the specific NSPC populations responsive to various injury or disease paradigms. Theoretically, only a subset of NSPCs could react to a certain disease and only these cells could underlie the extensive regenerative ability of zebrafish brain.

### Molecular characterization of individual cell types

Based on the cell clustering (Figure 3A) and identified marker genes (Figure 3B; Dataset S1A), we determined the gene ontology (GO) terms that are enriched for every cell cluster (Figure 3C; Dataset S1C,D). In total, 1,632 GO-terms for biological process (BP) and 842 GO-terms for molecular function (MF) were ascertained to all clusters (Figure 3C). Among these, 841 BP GO-terms and 259 MF GO-terms were uniquely ascertained to only one cluster (Figure 3C; Dataset S1F,H). The highest number of GO-terms was found for NB (246 unique and 124 non-unique BP+MF combined) and lowest for NE (26 unique and 12 non-unique BP+MF combined) (Figure 3C; Dataset S1D-H). Among the BP GO-terms that were ascertained to more than one cluster were terms related to general developmental and maintenance programs for zebrafish brain such as central nervous system development, neurogenesis, Notch signaling, cellular macromolecule biosynthesis, regulation of neurotransmitter levels and cell cycle regulation (Figure 3D; Dataset S1E). Unique BP GO-terms included various signaling pathways such as Wnt signaling (dRGC1), Insulin signaling (mRGC1), TGFb signaling (mRGC2), MAPK signaling (dRGC2), Estrogen signaling (vRGC2), cell cycle (NB), Retinoic acid signaling (OD), and JAK-STAT signaling (CPN) (Figure 3E; Dataset S1F). For MF GO-terms, we have found generic categories that are involved in neuronal activity, intracellular signal transduction and energy metabolism such as Calcium ion binding, Fatty-acyl CoA binding, oxidoreductase activity and heparin binding (Figure 3F, Dataset S1G). Unique MF GO-terms included various signaling pathways and receptor activity such as Wnt/Frizzled binding (dRGC1), epinephrine binding (mRGC1), collagen binding (vRGC1), retinoic acid binding (vRGC2), chromatin binding (NB), insulin binding (NE), and glutamate receptor binding (GIs+) (Figure 3G, Dataset S1H). Many of these signaling pathways were shown to be required for normal functioning and plasticity of the zebrafish brain (Bhattarai et al., 2016b; Diotel et al., 2013; Jiao et al., 2011; Shimizu et al., 2018; Sloin et al., 2018; Tallafuss et al., 2015; Than-Trong et al., 2018); however, their specific involvement for certain subtypes of NSPCs was not shown before. Therefore, our results provide a detailed resource on specific molecular regulatory networks, constitute a starting point for underlying the heterogeneity of the NSPC populations, and serves as a comprehensive repository for researchers interested in certain molecular programs.

Transcription factors (TF) are master regulators for cell fate determination and a plethora of cellular functions. Therefore, the TF expression codes for cell clusters could determine their lineage relationship, cellular physiology and the “stemness” of the NSPCs. We found that 112 TFs serve as marker gene for at least one cell cluster (Figure 3H, Dataset S1I). These TFs included known regulators of NSPC plasticity such as *sox2 (Lam et al., 2009)*, *id1 (Diotel et al., 2015)*, *neurod1 (Guo et al., 2014)*, *ascl1a (Karow et al., 2012)*, *hey1 (Than-Trong et al., 2018)*, *gsx2 (Pei et al., 2011)*, *zic3 (Sassa et al., 2007)* and neuronal markers *dlx5a (Wang et al., 2010)*, *eomesa (Ganz et al., 2011)*, *nkx2.1 (Ganz et al., 2011)*, *ctnnb1 (Wang et al., 2012)* (Figure 3I). When we determined the TF code similarity matrix (percentage of the TFs expressed in one cell cluster to the compared cell cluster) (Figure 3J), we found that the TF expression patterns in dRGC1, dRGC2 and mRGC1 share more than 60% similarity suggesting that these progenitor cells might represent a closer developmental origin and physiology. Additionally, 64% of TFs that are expressed in NB are also expressed in mRGC2, suggesting that NBs might be mainly derived from mRGC2 cells (Figure 3J). When the similarity index for TFs expressed in progenitor cells was compared to neuronal populations (Figure 3J), we found that the majority of the neuronal clusters share TF expression patterns with dRGC1 and dRGC2 with the exception of CSN that is more similar to mRGC1 in terms of TF code. These results suggest that the majority of neuronal subtypes in adult zebrafish telencephalon might derive from dRGCs and NBs might derive from mRGCs (Figure 3K). This prediction is supported by previous documentation of cortical neurons deriving from dorsal region of the vertebrate telencephalon (Cai et al., 2013; Radonjic et al., 2014) and a large portion of the proliferating cells in the zebrafish pallium are found in medial region (Adolf et al., 2006; Grandel et al., 2006; Kaslin et al., 2008; Kizil et al., 2012b; Than-Trong et al., 2018; Zupanc, 2008). Interestingly, we found that endothelial cells share 60% TF code similarity with dRGC populations (Figure 3J), suggesting that endothelial cells might be derived from dRGCs. A recent study where human brain organoids were transplanted into mouse brains, the nerovascularized organoid generated endothelial cells that are not of mouse but human origin (Mansour et al., 2018), suggesting that human neural progenitors might also contribute to production of endothelial cells and support our findings in zebrafish brain. Therefore, the TF code determination for individual cell clusters might give insight into novel aspects of the developmental origins and physiological relevance of NSPCs to individual neuronal and non-neuronal subtypes. Additionally, in disease conditions where specific neuronal subtypes are affected, molecular understanding of the heterogeneity of NSPC population would be instrumental to predict which progenitor cell types and molecular programs are employed by zebrafish brain to enable a successful regenerative response.

A prominent way cells interact with each other is through secreted molecules and their receptors in target cells. To determine such putative cell-cell communication through secreted molecules, we determined bioinformatically and based on the literature 646 ligands interacting with 658 receptors (Dataset S1J). Among the marker genes expressed in clusters (Dataset S1A), we found 101 ligands and 117 receptors were expressed in identified clusters (Dataset S1J). We generated an interaction map based on identified genes (Figure 3L, Dataset S1K) portraying a complex interaction map.

Many major signaling pathways such as Fgf, Wnt, Igf, Chemokine and Notch signaling are known to be involved in cell-cell communication in adult zebrafish brain (Alunni et al., 2013; Cacialli et al., 2016; Diotel et al., 2010; Jiao et al., 2011; Kaslin et al., 2009; Kizil et al., 2012a; Kizil et al., 2012c; Shimizu et al., 2018). To depict how cell clusters interact using these pathways, we plotted the intercommunication predictions for cell types expressing the receptor or ligand for these pathways (Figure 3M, Dataset S1L). We found that RGCs are highly regulated by Fgf signaling whereas RGCs generally affect other cell types using Wnt signaling. ODs are affected by Igf signaling and chemokines derived from mainly neurons regulate RGCs and NBs. Additionally, NBs are highly regulated by Notch signaling (Figure 3M). These results are consistent with published literature as Fgf signaling is important for NSPC proliferation (Ganz et al., 2010; Kaslin et al., 2009), Wnt signaling is active in adult zebrafish brain (Shimizu et al., 2018; Tallafuss et al., 2015), chemokine signaling is a prominent interaction mechanism for maintenance and regenerative ability of NSPCs (Belmadani et al., 2006; Diotel et al., 2010; Kizil et al., 2012a), and Notch signaling determines the neurogenic competency in neuronal progenitors (Alunni et al., 2013; Than-Trong et al., 2018). We also identified other potential interactions with less studied ligands and receptors such as *agrn*, *appa*, *ctgfa*, *gnai2*, *penkb*, *ptn*, *serpine*, *edil3a* and *hbegf* (Dataset S1L), which might provide novel candidates for regulation of NSPC plasticity. However, we still note the reservation that this map is a bioinformatically-designed map and does not take into account the spatial proximity of the cell types. Theoretically, distant cells may not have the capacity to interact through such ligand-receptor interaction, and the validation of our data will require experimental studies. Yet, our interaction studies provides an extensive high-resolution cell-cell communication map and provides a further level of elucidation for the heterogeneity of the NSPC populations and the influence of neuronal subtypes on NSPC plasticity.

### The effects of Aβ42 on NSPC plasticity and neuronal physiology in single cell resolution

Amyloid-beta42 (Aβ42) is one of the hallmarks of Alzheimer’s disease. Its direct effect on neuronal integrity and stem cell plasticity, and whether it is the causative agent for Alzheimer’s pathology are still controversial. We previously showed that Aβ42 treatment causes phenotypes reminiscent of human AD conditions, namely, cell death, synaptic degeneration, inflammation and cognitive deficits (Bhattarai et al., 2017a; Bhattarai et al., 2016b; Bhattarai et al., 2017b). However, in contrast to mammals, zebrafish NSPCs increased their proliferation and neurogenic output indicating that adult zebrafish brain responds to Aβ42 toxicity by enhancing NSPC plasticity and neuroregeneration (Bhattarai et al., 2016b; Bhattarai et al., 2017b). We also identified Interleukin-4 (IL4) as a crosstalk mechanism that activates the NSPCs and kick-starts the regeneration response (Bhattarai et al., 2016b). However, the cell types Aβ42 and IL4 affects and the molecular mechanisms affected in these cell types remained largely unknown. With our single cell analysis of cell types, their molecular programs, TF codes and cell-cell communication dynamics, we therefore extended our analyses to adult zebrafish brains treated with Aβ42 (Figure 4, Dataset S2A-K).

To determine the changes in gene expression in the cell types identified by unbiased clustering (Figure 4A), we performed differential gene expression analyses by comparing the Aβ42-treated brains with control brains (Figure 4B). Aβ42 led to differential expression of 475 genes on average per RGC cluster and NB, 427 genes per neuronal cluster, 267 genes in OD, 181 genes in endothelial cells, and 300 genes in immune cells (Figure 4B, Dataset S2A). GO-term analyses of differentially expressed genes showed that in total 1396 GO-terms for biological process (BP) and 669 GO-terms for molecular function (MF) were ascertained to all clusters (Figure 4C, Dataset S2B). Among these, 673 BP GO-terms and 445 MF GO-terms were uniquely ascertained to only one cluster (Figure 4C; Dataset S2B,C). The highest number of GO-terms was found for NB (102 unique and 94 non-unique BP+MF combined) and lowest for En (12 unique and 15 non-unique BP+MF combined) (Figure 4C; Dataset S2B-G). GO-term analyses revealed that biological pathways that are affected by Aβ42 in more than one cluster were related to aerobic respiration, synaptic transmission, neuronal development, metabolic processes, gene expression and response to stress (Figure 4C, Dataset S2B-D). When we analyzed the unique GO-terms ascertained to only one cell cluster (Dataset S2E), we found that Aβ42 affected a wide array of biological pathways related to (1) signaling mechanisms such as PDGF signaling in dRGC1, Wnt signaling in dRGC2, non-canonical Wnt signaling in mRGC2, Notch signaling in NB, Fgf signaling in CI1, Calcineurin signaling in GIs+, Nodal signaling in En, mTOR and p38 signaling in immune cells, (2) metabolic pathways such as isoprenoid synthesis in dRGC1, lipid metabolism in mRGC1, IL12 biosynthesis in CSN, phospholipid biosynthesis in GSN, polyprenol metabolism in GIs−, sphingolipid biosynthesis in OD, (3) stem cell plasticity pathways such as stem cell proliferation in dRGC2, radial glia migration in vRGC2, stem cell commitment in NE, cell cycle in NB, (4) neuronal development processes such as axon guidance in IN, cholinergic signaling in CPN, GABA synthesis in PPN, synaptic plasticity in GIs+, synapse maturation in MNP, neuromuscular junction formation in GSN, (5) behavioral pathways such as swimming behavior, memory, learning and cognition in CN, and (6) other processes such as autophagy in dRGC2, senescence in mRGC1, mitochondrial organization in MNP (Figure 4E). In terms of molecular functions (MF), antioxidant activity, oxidoreductase activity, RNA binding and transcriptional regulation activity were affected by Aβ42 (Figure 4F, Dataset S2F). For cluster-specific MF GO-terms, the most profound changes were observed in receptor activity (NGF and PDGF receptor binding in dRGC1, IL4 binding in mRGC2, IL1R activity in NB, CXCR binding in vRGC2, opioid receptor binding in GSN, glucocorticoid receptor activity in GIs−, L-DOPA binding in CN, histamine receptor activity in PPN, GABA receptor activity in CI1, TRH receptor binding in MNP, apelin receptor activity in En, Interferon and serotonin receptor activity in OD) (Figure 4G, Dataset S2G). These results indicate that Aβ42 treatment affects various cell types and diverse molecular programs. Majority of the molecular pathways we have found changing were not experimentally addressed in zebrafish brain; however, our single cell analysis results, which showed that IL4 signaling and its intracellular signal transduction route are affected by Aβ42 in mRGC2, NB, and CSN, are consistent with our previous findings that IL4 and interleukin signaling regulates NSPC plasticity in zebrafish and human (Bhattarai et al., 2016a; Papadimitriou et al., 2018). Thus, our results are likely to provide novel mechanistic insight into the effects of Aβ42 on NSPCs and neurons.

We identified 170 differentially expressed transcription factors (TFs) in all cell clusters after Aβ42 treatment (Figure 4H, I; Dataset S2H). Several TFs change their expression in RGC populations and in NB, for instance the TFs known to be associated with forebrain development and NSPC activity *fezf1* (upregulated in dRGCs and mRGC1), *her2* (upregulated in dRGCs and vRGCs), *ascl1a* (upregulated in dRGC2 and mRGC1), *msi1* (upregulated in dRGC2), *emx2* (upregulated in mRGC1), *sox3* (upregulated in vRGCs) and *zic3* (upregulated in dRGC1) (Figure 4I, Dataset S2H). Interestingly, a master regulator of neural stem cell plasticity *sox2* is upregulated in all RGC clusters as well as in NBs and in no other cell type after Aβ42 treatment (Figure 4I). This indicates an interesting finding that Aβ42 enhances NSPC plasticity in all glial cells and neuroblasts and provides a molecular explanation on our previous findings that NSPC proliferation increases after Aβ42 (Bhattarai et al., 2016a; Bhattarai et al., 2017a; Bhattarai et al., 2017b). Therefore, we propose that the regenerative ability of adult zebrafish brain after Aβ42 toxicity might rely on the ability of the NSPCs to upregulate *sox2* expression. In mammalian neural stem cells, Aβ42 treatment was shown to reduce *sox2* expression (Choi et al., 2014; Jung et al., 2012; Papadimitriou et al., 2018; Tincer et al., 2016), and therefore our model and single cell analyses can provide us with novel insights into how NSPCs can be converted to a more plastic state in Alzheimer’s disease conditions. Additionally, we identified many differentially expressed transcription factors in neurons, which can give insight into the effects of Aβ42 on individual neuronal subtypes (Dataset S2H).

To determine how Aβ42 affects the cell-cell communication through secreted molecules and their receptors, we generated an interaction map for cell clusters in Aβ42-treated adult zebrafish brain (Figure 4J, Dataset S2I-K). We determined the interactions that are differentially regulated after Aβ42. Here, the red lines represent the interactions that are not present in control brains but only after Aβ42 treatment and the green lines represent the interactions that are lost after Aβ42 treatment (Figure 4J). We found a complex change in the secreted molecule/ligand-based interaction. For instance the neuronal population GSN no longer interacts with the neurogenic populations (blue dots – RGCs, NE and NB) by Fgf and Bmp signaling, Aβ42 starts the regulation of NBs by Bmp signaling, several neuronal subtypes undergo a regulation by chemokine signaling emanating from neurogenic populations, vRGCs are no longer regulated by Notch, Appa signaling is potentiated in almost all cell populations but mRGC1, Ctgfa signaling is potentiated in ODs, endothelial cells activate Egf and Penkb-mediated interaction, Agrn signaling is lost in many cell types but activated in neuroepithelium (Figure 4J, Dataset S2I-K). These results suggest a plethora of changes in potential interactions between different cell types in adult zebrafish brain after Aβ42 treatment. This understanding is likely to provide important information on how neurogenic plasticity of stem cells are regulated by other stem cells or neurons and how Aβ42 pathology affect stem cells to elicit a neuro-regenerative response. We anticipate that the candidates that can be identified in our screens could be used towards efforts to better understand Alzheimer’s disease and potential stem cell-based regenerative therapies in humans.

### The effects of Interleukin-4 on adult zebrafish brain in single cell resolution

We previously showed that Interleukin-4 (IL4) is a response to Aβ42 toxicity in adult zebrafish brain and it increases the proliferative and neurogenic capacity of NSPCs in both zebrafish and humans (Bhattarai et al., 2016b; Bhattarai et al., 2017b; Kizil, 2018; Papadimitriou et al., 2018). However, the effects of IL4 on individual NSPC populations and neurons in zebrafish as well as the mechanisms by which it regulates NSPC plasticity remained unknown. To address the effects of IL4 in adult zebrafish brain, we performed a single cell sequencing on IL4-treated brains (Figure 5). After tSNE analyses (Figure 5A) and comparison to control cells (Figure 5B), we found that IL4-treated brains contain all clusters except for the Aβ42-specific CI2 cluster (Figure 5C). To determine the cell clusters that would be responsive to IL4, we determined the *il4r* (il4 receptor) expression in control cell clusters (Figure 5D) and found that all RGC clusters, NBs, NEs and immune cells express *il4r*, which is not expressed in any neuronal populations (Figure 5D). This suggests that IL4 signaling is mainly acting on RGCs, proliferating neuroblasts and immune cells. Therefore, for the rest of our analyses, we separated *il4r+* clusters from *il4r*-clusters.

To determine the changes in gene expression exerted by IL4, we performed differential gene expression analyses by comparing the IL4-treated brains with control brains (Figure 5B). IL4 altered the expression levels of 4211 different genes in *il4r+* (2955 genes) and *ilr4−* (1256 genes) clusters (Figure 5E, Dataset S3A).

Since Amyloid treatment leads to IL4 expression in adult zebrafish brain and both Aβ42 and IL4 changes expression of a number of genes, we hypothesized that if IL4 conveys the effects of Aβ42, the differentially expressed genes in Aβ42 treated brains and IL4 treated brains should correlate in *il4*-responsive cell clusters. Therefore, we plotted the differentially expressed genes (DEGs) that are in common between Aβ42 and IL4-treated samples (Figure 5F, Dataset S3I). Interestingly, on average 35.1% of the DEGs are common between Aβ42 and IL4 samples in *il4r+* clusters including the RGCs and NB, while in *il4r−* clusters the average overlap is only 12.3% (Figure 5F). The highest overlap is observed in NE with 54.4% (75 genes) and the highest number of genes overlapping is in dRGC1 (200 genes, 43.1%) (Figure 5F). These results clearly indicate that IL4 signaling is affecting RGCS and NBs, a significant portion of DEGs in RGC clusters and NBs after Aβ42 treatment are produced by IL4, and the effects of Aβ42 in *il4r−* clusters are conveyed through other mechanisms (e.g.: direct effect of Aβ42 or indirect effect through another intermediary program). Therefore, our findings propose IL4 as a prominent and important mechanism that controls NSPC plasticity directly by activating its downstream signaling cascade and IL4 is an important but not the only mediator of the effects of Aβ42 in RGCs.

GO-term analyses of differentially expressed genes showed that in total 1483 GO-terms for biological process (BP) and 722 GO-terms for molecular function (MF) were ascertained to all clusters (Figure 5G, Dataset S3B). Among these, 677 BP GO-terms and 481 MF GO-terms were uniquely ascertained to only one cluster (Figure 5G; Dataset S3B,C). The highest number of GO-terms was found for NB (123 unique and 138 non-unique BP+MF combined) (Figure 5G,H; Dataset S3B-G). GO-term analyses revealed that biological pathways that are affected by IL4 in more than one cluster were including those that are related to brain development, metabolic processes and gene expression (Figure 5H, Dataset S3B-D). When we analyzed the unique GO-terms ascertained to only one cell cluster (Figure 5I, Dataset S3D,F), we found that *il4r+* clusters are affected in (1) metabolic pathways (steroids in dRGC1, tetrapyrolle in dRGC2, choline in mRGC1, glutamine in dRGC2, cholesterol and acetyl-CoA in vRGC2, polyol in Im), (2) signaling pathways (MAPK, TGFb, ERK1, VEGF in dRGC1, GCSF, IL8 and BMP in dRGC2, NFkB in mRGC2, TOR in vRGC1, PKB in NB, Hippo in NE, (3) cell-cell and cell-matrix interaction (matrix adhesion and integrin binding in dRGC1, sheet morphogenesis in dRGC2, ciliary movement in mRGC1, epithelial migration in NE), and (4) regulatory mechanisms (apoptosis in dRGC2, histone methylation in mRGC1, miRNA activity in mRGC2, autophagy in vRGC1, mitochondrial transport in NE, chromosome organization and cell division in NB, antigen presentation and ECM remodeling in Im) (Figure 5I, Dataset S3D,F). In *il4r−* clusters, we also observed uniquely affected GO-term categories such as Activin, Nodal and Retinoic acid signaling in IN, Interferon signaling in CSN, Insulin signaling in CI1, Fgf signaling in MNP, Wnt signaling in GIs+, Dopamine, serotonin and thyroid hormone signaling in CPN, NGF signaling in En, transcytosis in IN, positive regulation of axon regeneration and oxygen transport in GSN, Amyloid precursor protein catabolism in GIs−, neuronal fasciculation in CN, ketone body catabolism in PPN (Figure 5I, Dataset S3D,F). We hypothesize that the effects of IL4 on *il4r−* clusters could be indirect either through IL4-dependent regulation of immune system and/or through the regulatory activity of NSCs on various neuronal subtypes.

When we analyzed the changes in transcription factors upon IL4 treatment (Dataset S3H), we found 109 TFs are differentially expressed in *il4r+* clusters (Figure 5J, Dataset S3H) and 59 in *il4r−* clusters (Figure 5K, Dataset S3H). *foxj1a*, *fosab*, *her6*, *rfx4*, *fosb*, *jun*, *sox2* and *mycb* are among the TFs that are upregulated in several RGC clusters and NBs. The highest number of TFs is differentially expressed in dRGC1, suggesting that IL4 might affect the dorsal RGCs most. *sox2*, one of the master regulators of stem cell plasticity and the TF upregulated in all RGC clusters and NB upon Aβ42, is upregulated in dRGC1, dRGC2, mRGC1 and NB and partially recapitulates the effects of Aβ42 (Figure 5J). In *il4r−* clusters, 59 TFs are differentially expressed and we consider those changes as indirect effects of IL4 as these cells do not have detectable levels of *il4r* transcript (Figure 5D,K).

When we analyzed the cell-cell interaction map based on the secreted molecules and their receptors (Figure 5L, Dataset S3J-L), we observed that compared to the controls, IL4 mainly downregulates the interaction through Bmp, chemokines, Ctgfa, Penkb, Agrn and Serpine (Figure 5L). In IL4-treated brains, some neurogenic populations (dRGC2, mRGC1, NE1) activate Notch signaling while others (dRGC1, mRGC2, vRGC2, NE2, and NB) are relieved of Notch regulation (Figure 5L, Dataset S3L). Interestingly, IL4 similar to Aβ42 activated the Appa signaling in almost all cell types except for the neuroepithelium (Figure 5L), suggesting that IL4 and Aβ42 have synergistic effects on the extent of Appa signaling. IL4 suppresses the Egf signaling in vRGC1, mRGC1 and dRGC1 populations, which we believe are quiescent progenitors (Figure 2, Figure 3) and activates the Egf signaling in vRGC2, mRGC2 and dRGC2 populations which are activated progenitors (Figure 5L). Given that EGF signaling defines the activated neural stem cell populations in mouse brains (Codega et al., 2014), our results further supports the pro-neurogenic and activating role of IL4 on neural stem cells.

Further investigation of our data and experimental validation of the candidates proposed in this manuscript could provide us with the molecular programs underlying the NSPC plasticity and regenerative response of adult zebrafish brain in Alzheimer’s disease conditions as well as the downstream regulation of IL4. Given that we have recently shown IL4 was sufficient to enhance the human NSPC plasticity and regenerative capacity in a 3D human Alzheimer’s model and circumvent the AD pathology (Papadimitriou et al., 2018), our data will undoubtedly provide more candidates that would be clinically relevant. Such an understanding would provide new therapeutic targets for clinical and pharmaceutical use in Alzheimer’s disease.

### Conclusion

Heterogeneity of neural stem cell activity and fate decisions mainly rely on activation or suppression of distinct molecular programs in respective cell types. Single cell sequencing uniquely allows investigation of individual cell types and their response to stimuli. In our analyses, we identified distinct cell types of adult zebrafish telencephalon, which is a widely-used experimental model for studying neuronal regeneration and neural stem cell plasticity. We also identified how Amyloid-beta42, the hallmark of Alzheimer’s disease, affected these cell types and which genes and pathways are altered upon Amyloid toxicity. Since zebrafish brain can enhance its stem cell activity and neurogenesis in Alzheimer’s disease conditions, understanding in more detail how Ab42 affects individual cells would certainly enhance our understanding how a vertebrate brain could counteract Alzheimer’s pathology and could propose novel targets for clinical use or drug development. Furthermore, we also investigated the activity of IL4, a key molecule that is required to activate the stem cell proliferation and neurogenesis in AD conditions in zebrafish. For both of these conditions, we identified numerous pathways and genes differentially regulated after the treatments.

Our results provide complex and multi-layered information. First, it provides a refinement of the cell types and progenitor states in the adult zebrafish telencephalon (Figures 1 and 2). This will allow investigation of certain regions or cell types in a higher resolution and can allow cell-specific investigation of biological phenomena. Our analyses of the pathways that are active in certain clusters in homeostatic states of the fish brain also provided a delineation of the pathways that were already known to be involved in stem cell activity and neurogenesis (e.g: Wnt, Fgf, Bmp) into certain cell types (Figure 3). Parallelism between our results and previous literature confirms the fidelity of our results (for instance Notch signaling affecting neuroblasts, Fgf signaling affecting the RGC populations, chemokine signaling emanating from immune cells and neurons affecting the RGC populations). However, in addition to known signaling molecules, our data provides extensive resource for identification of new molecules and cellular interactions in adult fish brain. Additionally, our analyses with Amyloid-beta42 (Figure 4) and IL4 (Figure 5) treatments provide unprecedented information on the stem cell plasticity and regenerative output of fish brain in Alzheimer’s conditions. Interestingly, Aβ42 treatment potentiates the sox2 expression in all RGC populations. This could be the key difference between humans and fish and might underlie the plasticity response and regenerative output in fish brains. Additionally, we identified Interleukin signaling affected by Aβ42 treatment verifying our previous findings. Interestingly, IL4 treatment – that affects RGC population, immune cells and neuroblasts as determined by the presence of IL4 receptor *il4r*, mimicked the differential expression of one third of the genes in RGC clusters when compared to Aβ42. This suggests that IL4 is a major component to regulate the neural stem cell plasticity in zebrafish brain after Aβ42 toxicity, and proposes other pathways involved. We are confident that with further investigation of our data, such molecular programs could be identified. We also believe that our datasets and investigation tools will provide a reference for studies on cell-cell interactions, stem cell plasticity, neurogenesis, regeneration and disease modeling in adult zebrafish brain.

## Materials and Methods

### Ethics statement

All animal experiments were performed under the permission of Landesdirektion Dresden with the following permit numbers: TVV-52/2015 with all relevant amendments.

### Cerebroventricular microinjection

6 months post fertilization (mpf) female Tg(her4.1:GFP) fish (Yeo et al., 2007) were injected either with PBS (control), Aβ42 (20 μM) or IL4 (1 μM), and were kept at 14 hr light/10 hr dark cycle in normal water system for 24 hours. Experimental fish were sacrificed using 0.2 % MESAB according to the animal experimentation permits.

### Cell Dissociation and sorting

The telencephalon of the fish were dissected in ice-cold PBS and directly dissociated with Neural Tissue Dissociation Kit (Miltenyi) at 28.5 °C as described previously (Bhattarai et al., 2016b). After dissociation, cells were filtered through 40 μM cell strainer into 10 ml 2% BSA in PBS, centrifuged at 300g for 10 min and cells resuspended in 4% BSA in PBS. Viability indicator (Propidium iodide) and GFP were used to sort viable GFP(+) or GFP(−) cells by FACS. The resulting single cell suspension was promptly loaded on the 10X Chromium system (Zheng et al., 2016). 10X libraries were prepared as per the manufacturer’s instructions. The raw sequencing data was processed by the cell ranger software provided by the 10X genomics with the default options. The reads were aligned to zebrafish reference transcriptome (ENSEMBL Zv10, release 91) and eGFP CDS (with arbitrary ensemble ID; ENSDARG99999999999). The resulting matrices were used as input for downstream data analysis by Seurat (Butler et al., 2018).

### Data Analysis by Seurat

All matrices were read by Read10X function; cell names were named and numbered as sample names and the column number, to trace back cells if required. We filtered out cells as following; cell with more than 15000 UMI or less than 1000 UMI, cells with less than 200 unique genes. The genes found less than 10 cells were filtered out. The remaining cells and genes were used for downstream analysis for all samples. (These are arbitrary thresholds! We could use some other thresholds, such as mitochondrial genes, which are marker of stress if higher than 6% in total UMI). The data normalized by using “LogNormalize” method, data scaled with “scale.factor = 1e4”. For each datasets variable genes found with FindVariableGenes with the default options. The top1000 most variable genes from each sample were merged, and the intersection of these with all variable genes in each sample were used for CCA analysis. The 4 Seurat object and the variable genes found above were used to generate a new Seurat object with RunMultiCCA function, using num.ccs = 50. The canonical correlation strength were calculated using num.dims = 1:20, the number dims determined after testing different num.dims upto 50, and 20 was found to be the best. The reudtion was done by using reduction.type = “pca”, dims.use = 1:20, then the data subset by using “var.ratio.pca” and “accept.low =”. The samples were aligned using dims.align = 1:20. The cell cluster were found using aligned CCA and 1:20 dims, with higher resolution 2.5, cluster were shown on 2D using t-SNE (RunTSNE function). Two cell clusters were found either with less 3 cells from at least one of samples, or no cells from any cells (cell number 6 and 20). (Most probably this is because of cell number depth, we have variable number of GFP(+) or GFP(−), this most probably cause the variation in cell types. Still, there is a possibility anew cell type or state might be induced/changed after treatments). When not possible, we did not calculate marker genes or differentially expressed genes for these cell clusters. Then we find the marker gens using FindConservedMarkers by comparing one cell cluster with the remaining all cells to find specific markers. (However, one marker can be highly expressed in a different cluster and can be seen in heatmap or dotplot). We used the genes from literature (Dataset S1B) and markers found by Seurat functions to define our cell types. For that, we generated FeaturePlot for all marker genes and checked the expression and specificity of the marker to define cell types.

### Identification of cell types

Feature plots were generated by Seurat software (Figure S3) and cell types were determined by the expression of marker genes *adma*, *aldocb*, *aplnra*, *aplnrb*, *apoea*, *apoeb*, *ascl1a*, *aurkb*, *bdnf*, *btg1*, *cahz*, *calb2a*, *calm1b*, *calm2a*, *cbln1*, *ccna2*, *ccr9a*, *cd59*, *cd63*, *cd74a*, *cd74b*, *cd82a*, *cd8b*, *cd99*, *cd99l2*, *cdk1*, *cdk2*, *cdon*, *chata*, *chrna1*, *chrnb1*, *chrnb2*, *clstn2*, *clu*, *ctgfa*, *cux1b*, *cx43*, *cx43.4*, *cxcl12a*, *cxcl18b*, *cxcr4b*, *cxcr5*, *cysltr1*, *cysltr2*, *cysltr3*, *ddc*, *dla*, *dld*, *dlx1a*, *dlx5a*, *ebf3a*, *edf1*, *eef1g*, *egfra*, *elavl3*, *elavl4*, *elov1b*, *emx2*, *eno1a*, *eno1b*, *eno2*, *eomesa*, *etv5a*, *fabp7a*, *fasn*, *fezf1*, *fli1a*, *fli1b*, *foxb1a*, *foxg1a*, *foxj1a*, *foxp3a*, *foxp3b*, *foxp4*, *fxyd6l*, *gabra5*, *gabrb3*, *gabrr2b*, *gad1b*, *gad2*, *gap43*, *gfap*, *GFP*, *glula*, *glulb*, *gsx2*, *hbaa1*, *hbaa2*, *hbba2*, *hepacama*, *her4.1*, *her4.2*, *her4.3*, *her4.4*, *hopx*, *htr1aa*, *htr1ab*, *htra1b*, *icn*, *id1*, *id2a*, *il10*, *il12bb*, *il4*, *il4r*, *il4r.1*, *isl1*, *kdrl*, *krt18*, *krt8*, *laptm5*, *lck*, *lcp1*, *lhx2a*, *lhx2b*, *lhx5*, *lhx8a*, *lmo3*, *lmo4b*, *lpar5a*, *map1aa*, *map1ab*, *marcksl1b*, *mbpa*, *mcm5*, *mdka*, *mki67*, *mpeg1.1*, *mpeg1.2*, *mpeg1.3*, *mpx*, *msi1*, *msi2b*, *nav3*, *nell2b*, *neurod1*, *neurod6a*, *neurog1*, *nkx2.1*, *notch1a*, *notch1b*, *notch3*, *npas4b*, *nrgna*, *nrxm1a*, *olig1*, *olig2*, *pax6a*, *pclaf*, *pcna*, *pcp4a*, *pcp4b*, *pecam1*, *penkb*, *pvalb7*, *pvalb7a*, *s100a10b*, *s100b*, *scrt1b*, *six3a*, *six3b*, *slc1a3b*, *slc1a6*, *slc2a3a*, *slc32a1*, *slc43a3b*, *slc6a1b*, *slit3*, *snap25*, *sox10*, *sox1a*, *sox2*, *sox3*, *sox6*, *sparcl1*, *stmn1a*, *stmn1b*, *sv2a*, *sypb*, *syt1a*, *tagln2*, *tbr1a*, *tbr1b*, *tnfa*, *tnfb*, *tph2*, *tubb2*, *vegfaa*, *vim*, *wasb*, *zic4* that define specific cell types (Figure 2, Dataset S1B).

### GO-term analyses

We used the all marker genes with False Detection Rate < 0.1 for Gene Ontology analysis and KEGG pathway analysis using GOStats (1.7.4) (Falcon and Gentleman, 2007) and GSEABase (1.40.1), p-value < 0.05 as threshold as described previously (Papadimitriou et al., 2018). To determine the differentially expressed genes (DEGs) after Aβ42 and IL4 treatment, we used FindMarkers function using cell cluster that have at least 3 cells from all samples. Then, we used the p-value <0.05 for significantly expressed genes. These genes were used for GO and pathway analysis using the scripts that are available on https://kizillab.org/resources.

### Transcription factors

We determined the transcription factors by using the GO-term 0003700 (https://www.ebi.ac.uk/QuickGO/term/GO:0003700). We determined the zebrafish orthologues of these TFs using SCENIC (Aibar et al., 2017). The TFs expressed in individual cell types are given in Datasets S1I, S2H, and S3H.

### Construction of interaction maps

For the ligand-receptor interaction, we downloaded all Ligand and receptors from a previous publication (Ramilowski et al., 2015) (http://fantom.gsc.riken.jp/5/suppl/Ramilowski_et_al_2015/vis/#/hive), and orthologs for zebrafish were found using BiomaRt of ENSEMBL. The ligand-receptor for zebrafish were used for downstream analysis as following; for cell-cell interaction, (i) all ligands found in 20% of a cell types were chosen, then their receptors were identified in all cell types. If a ligand-receptor pathway was found, then we draw a direction from the cell with the ligand to the cells with the receptor. We used igrap in R to draw interaction map as previously described (Skelly et al., 2018). In order to show the lost and/or new interaction after treatment, we calculated all the interaction for PBS and the treatment (Aβ42 or IL4) and then generated an interaction maps from all interaction of PBS and the treatment. Then, we colored the edges as green (lost after treatment), red (induced after treatment) and black that are not affected by the treatment.

## Author contributions

M.I.C. and C.K. conceived and designed the experiments. P.B. performed cerebroventricular microinjections. Y.Z. provided the Amyloid peptide, M.I.C. performed RNA isolation, M.I.C performed bioinformatics analyses, C.K. supervised the study, prepared the figures, and wrote the manuscript.

## Acknowledgements

This work was supported by DZNE and Helmholtz Association (VH-NG-1021, C.K.), DFG (KI1524/6, KI1524/10, and KI1524/11, C.K.), CRTD, TU Dresden (FZ-111, 043_261518). We would like to thank Drs. Andreas Petzold, Susanne Reinhardt and Andreas Dahl from CRTD NGS facility for library preparation.

## Supplementary Figure Legends

**Figure S1: Histograms for cell sorting.** (A) Control. (B) Aβ42-treated. (C) IL4-treated.

**Figure S2: Heat-map for cell type markers.** A high-resolution heat map for marker genes identified in individual cell clusters by Seurat software.

**Figure S3: Feature plot for marker genes.** Feature plot showing the expression of identified marker genes in individual cells. Violet points indicate expression.

## Supplementary Dataset Legends

**Dataset S1: Analyses on control cells.** (A) List of marker genes for cell clusters. (B) List of genes used for identification of cell types. (C) List of GO-terms. (D) GO-term chart. (E) Balloon chart for all GO-terms (biological process - BP). (F) Balloon chart for unique BP GO-terms. (G) Balloon chart for all GO-terms (molecular function - MF). (H) Balloon chart for unique MF GO-terms. (I) List of transcription factors expressed in individual cell types. (J) List of potential ligand-receptor interactions in zebrafish. (K) List of ligands and receptors expressed in control brains in individual cell types. (L) Interaction map for Bmp, Chemokine, Ephrin, Fgf, Igf, Notch, Wnt, Agrn, Appa, Ctgfa, Tgfb, Gnai1, Hbegfa, Penkb/Ptn, and Serpine.

**Dataset S2: Analyses on Aβ42-treated cells.** (A) List of differentially expressed genes (DEGs) in cell clusters after Aβ42 treatment. (B) List of GO-terms for DEGs. (C) GO-term chart. (D) Balloon chart for all GO-terms for DEGs (biological process - BP). (E) Balloon chart for unique BP GO-terms. (F) Balloon chart for all GO-terms for DEGs (molecular function - MF). (G) Balloon chart for unique MF GO-terms. (H) List of transcription factors expressed in individual cell types. (I) List of ligands expressed in control brains in individual cell types. (J) List of receptors expressed in control brains in individual cell types. (K) Interaction map for Bmp, Chemokine, Ephrin, Fgf, Igf, Notch, Wnt, Agrn, Appa, Ctgfa, Tgfb, Gnai1, Hbegfa, Penkb/Ptn, and Serpine.

**Dataset S3: Analyses on IL4-treated cells.** (A) List of differentially expressed genes (DEGs) in cell clusters after IL treatment. (B) List of GO-terms for DEGs. (C) Balloon chart for all GO-terms for DEGs (biological process - BP). (D) Balloon chart for unique BP GO-terms. (E) Balloon chart for all GO-terms for DEGs (molecular function - MF). (F) Balloon chart for unique MF GO-terms. (G) GO-term figure chart. (H) List of differentially expressed transcription factors in individual cell types after IL4 treatment. (I) List of TFs expressed in cell clusters after IL4 treatment. (J) List of ligands expressed in control brains in individual cell types. (K) List of receptors expressed in control brains in individual cell types. (L) Interaction map for Bmp, Chemokine, Ephrin, Fgf, Igf, Notch, Wnt, Agrn, Appa, Ctgfa, Tgfb, Gnai1, Hbegfa, Penkb/Ptn, and Serpine.

