## Supplementary material for "Single cell analyses of the effects of Amyloid-beta42 and Interleukin-4 on neural stem/progenitor cell plasticity in adult zebrafish brain": SD_3C_PBS_and_IL4_BP_Up_part1.pdf

**Balloon Plot for balloon.dfr.tmp, CellID by balloon.dfr.tmp, GO_TERM.**
**Area is proportional to balloon.dfr.tmp$GeneCounts.**

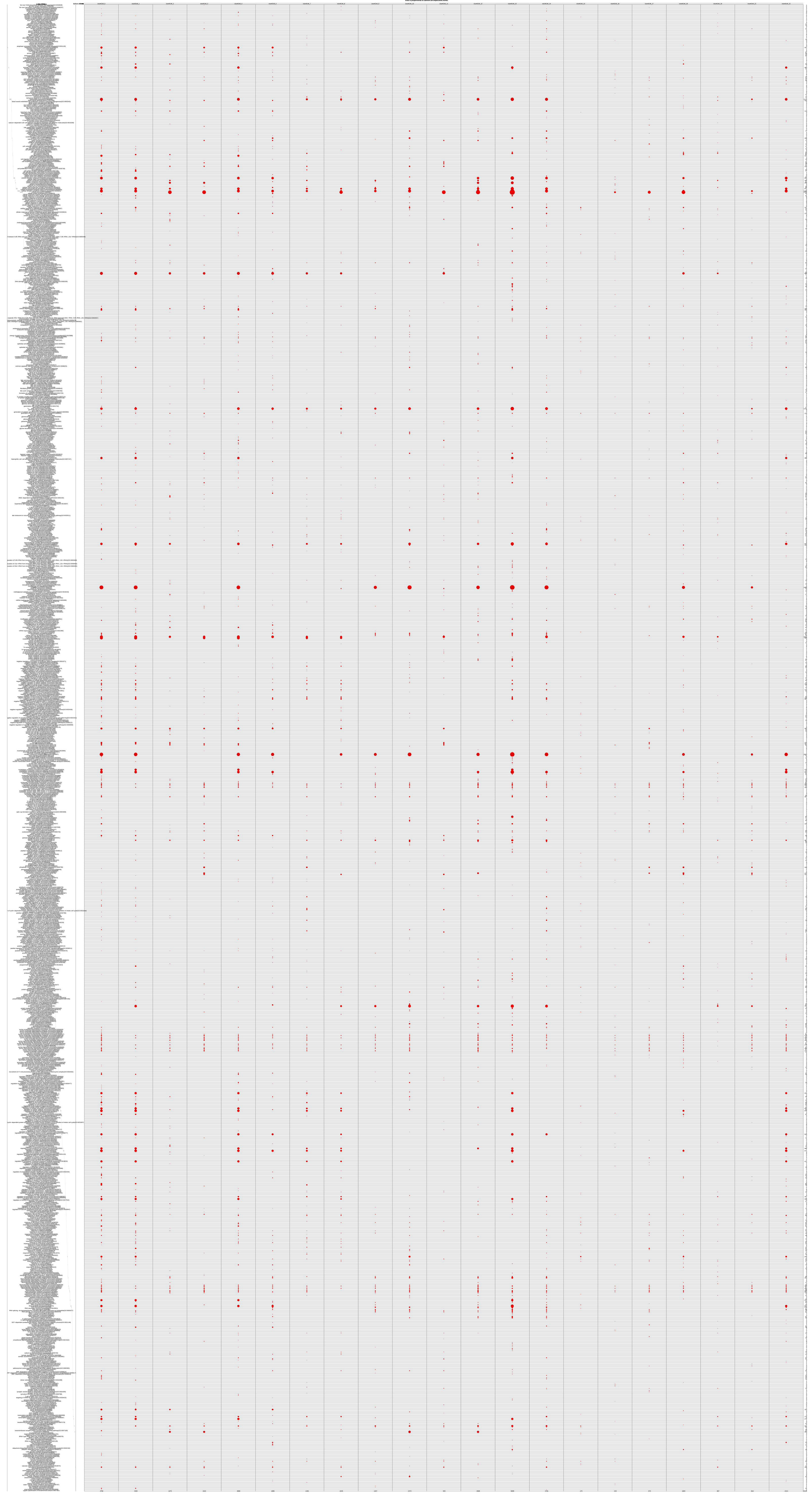
