## Supplementary figures and images for "Single cell analyses of the effects of Amyloid-beta42 and Interleukin-4 on neural stem/progenitor cell plasticity in adult zebrafish brain"

### SD_1F_Control_BP_unique.pdf

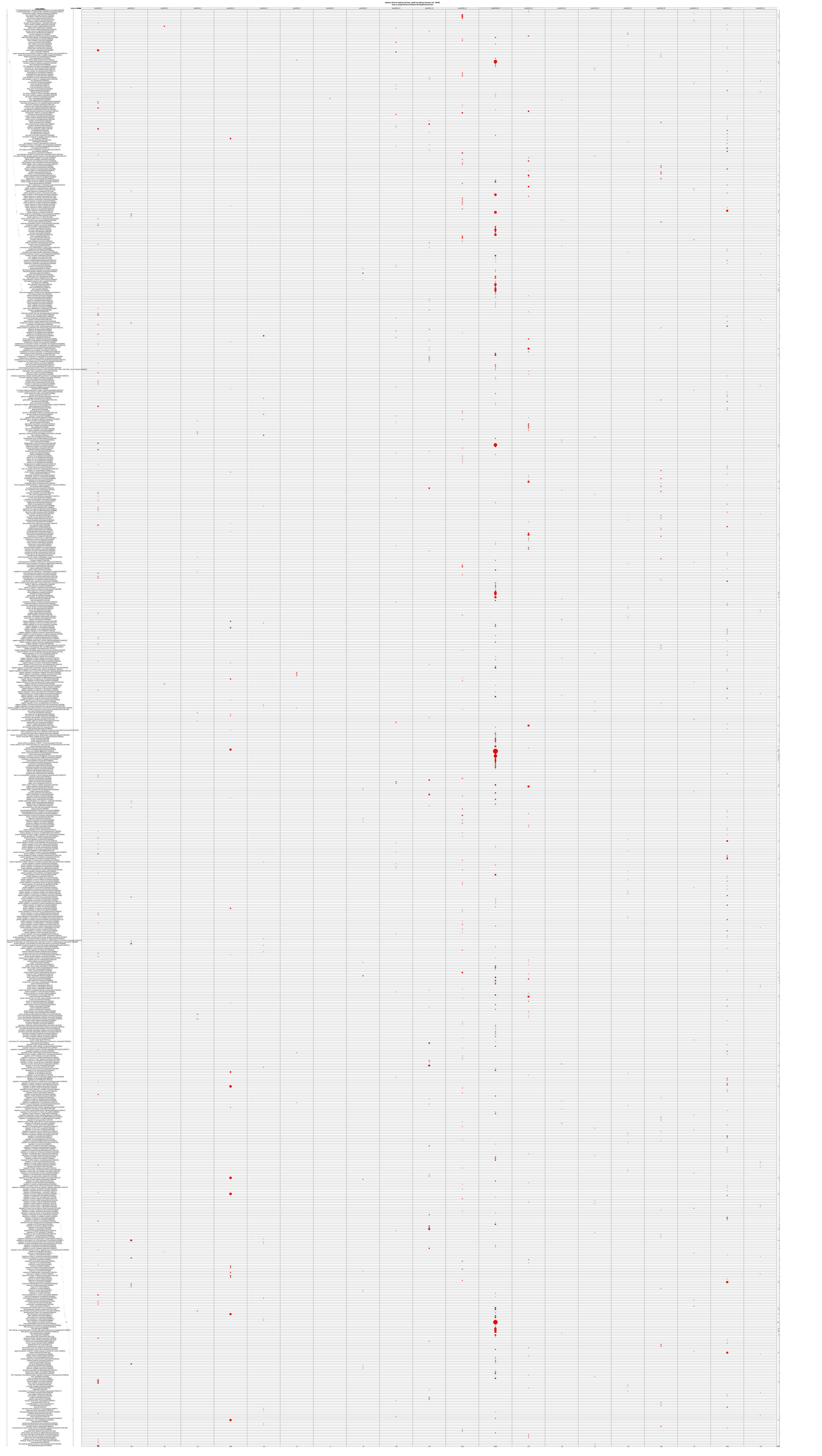

### SD_1G_Control_MF.pdf

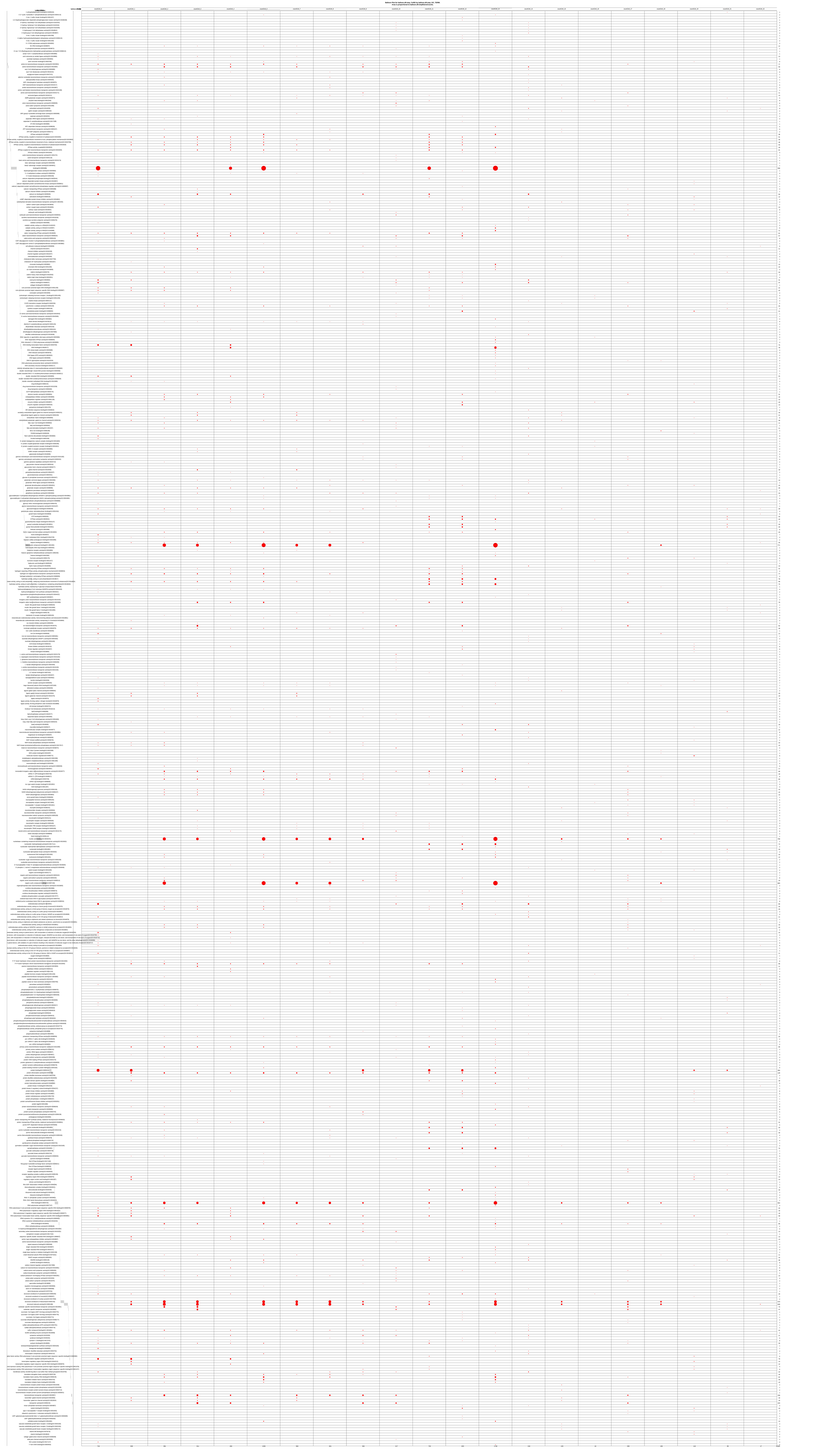

### SD_1L_PBS_ALL_interactions.pdf

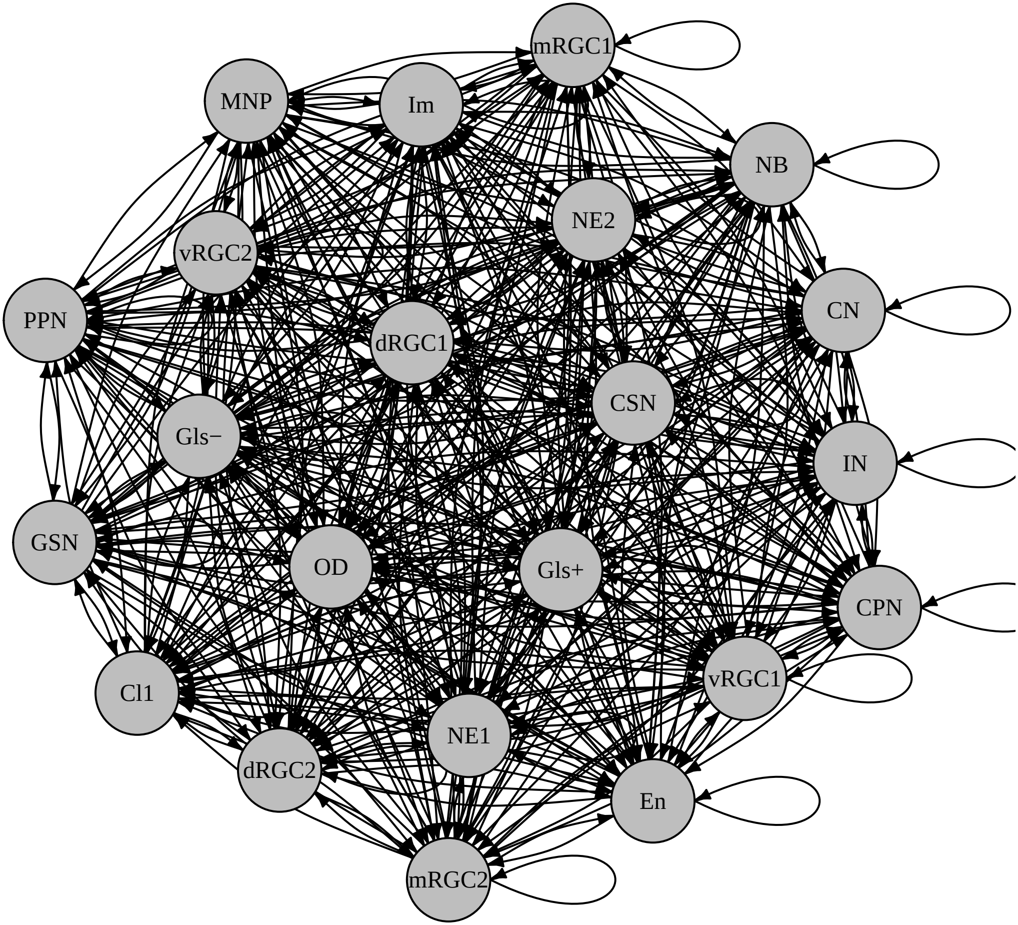

### SD_1L_PBS_uniqintrxs_bmp.pdf

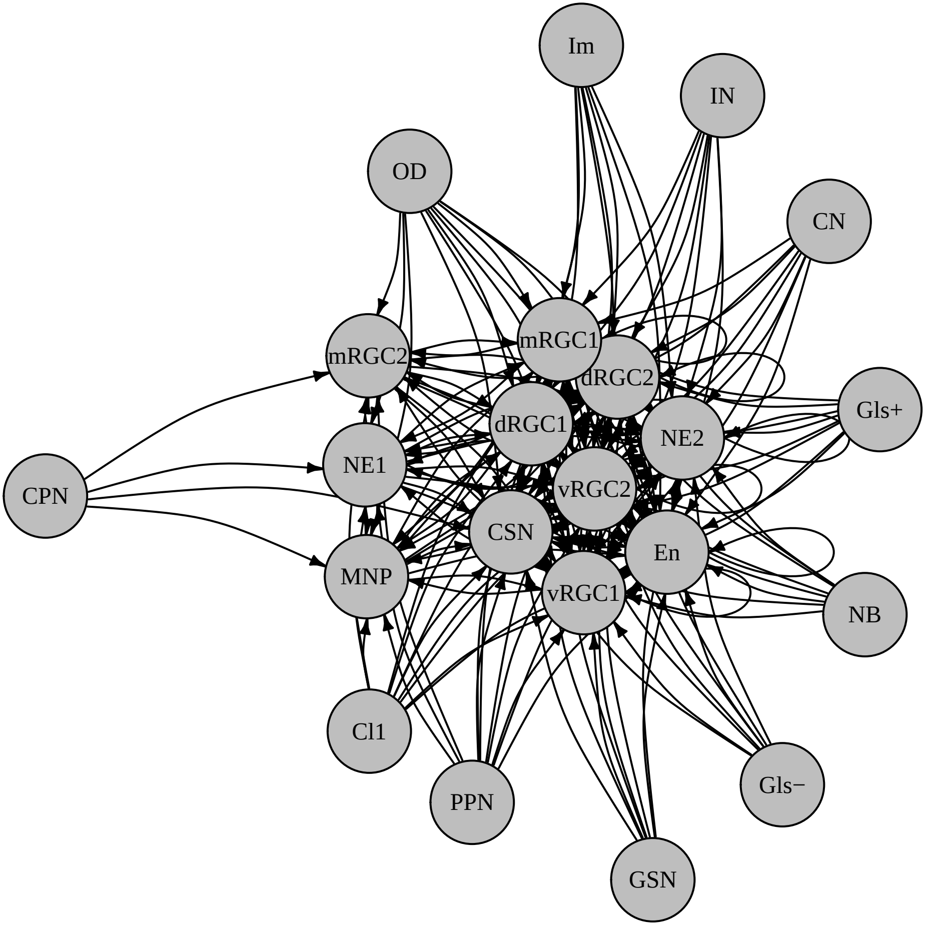

### SD_1L_PBS_uniqintrxs_chemokine.pdf

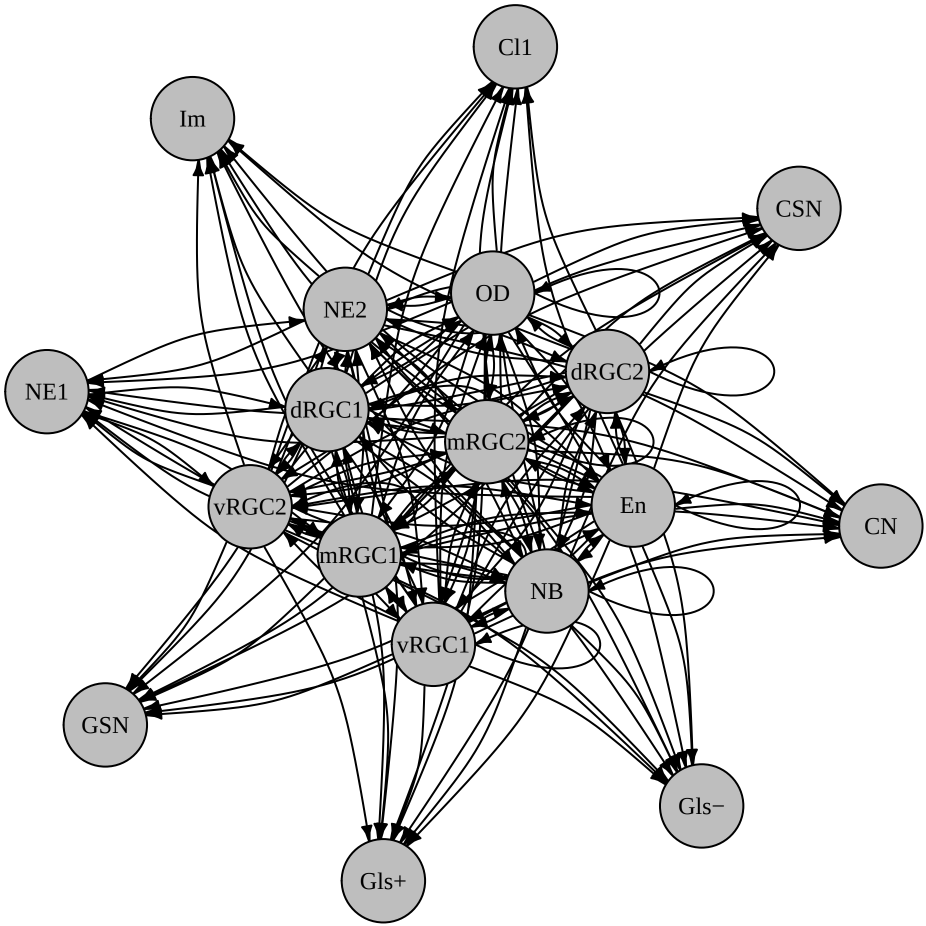

### SD_1L_PBS_uniqintrxs_epha.pdf

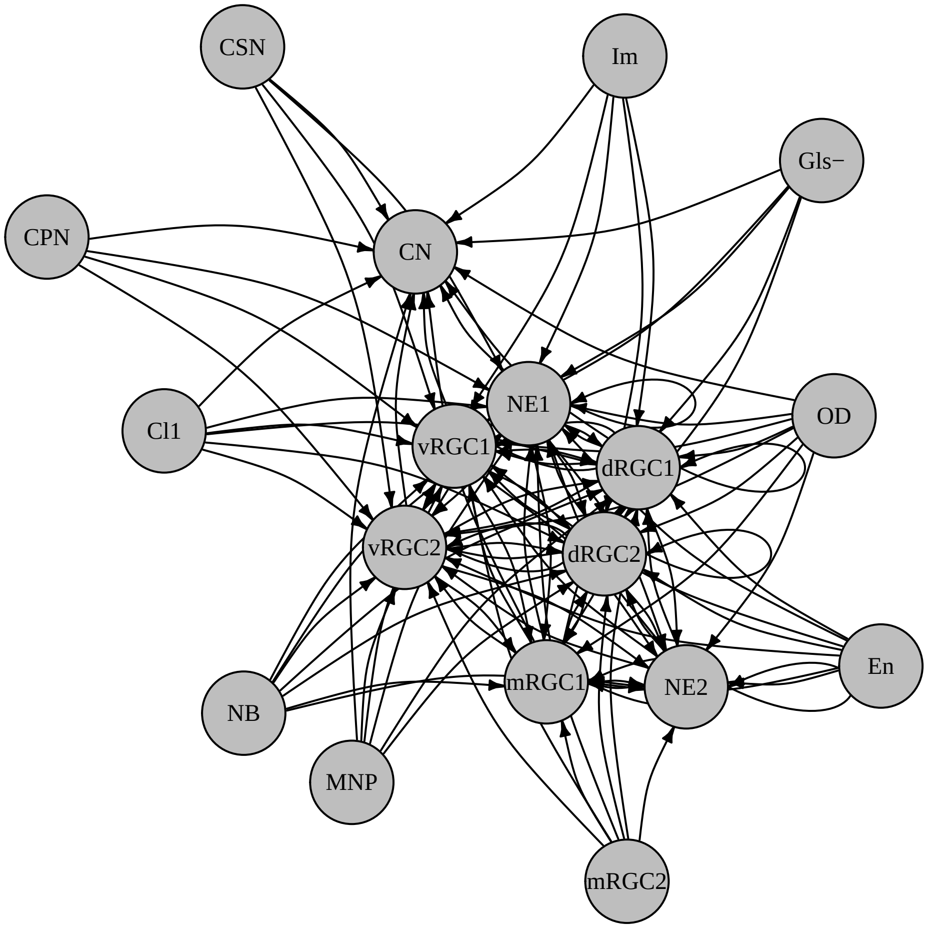

### SD_1L_PBS_uniqintrxs_fgf.pdf

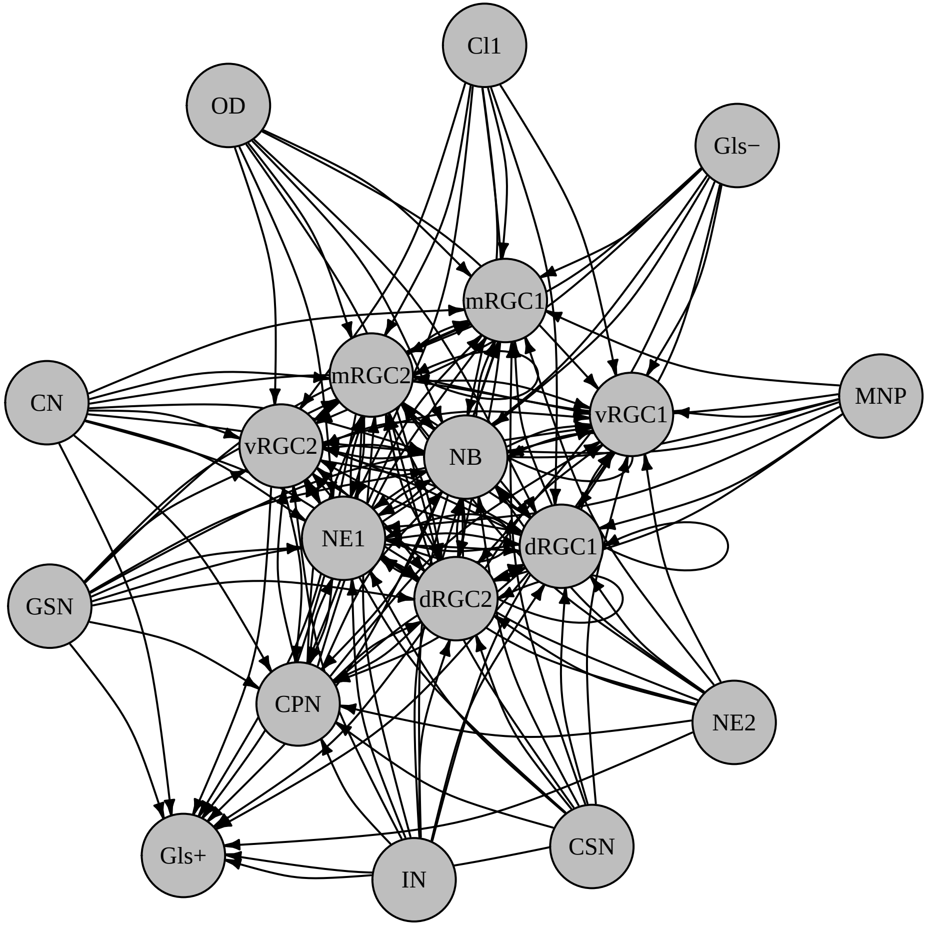

### SD_1L_PBS_uniqintrxs_igf.pdf

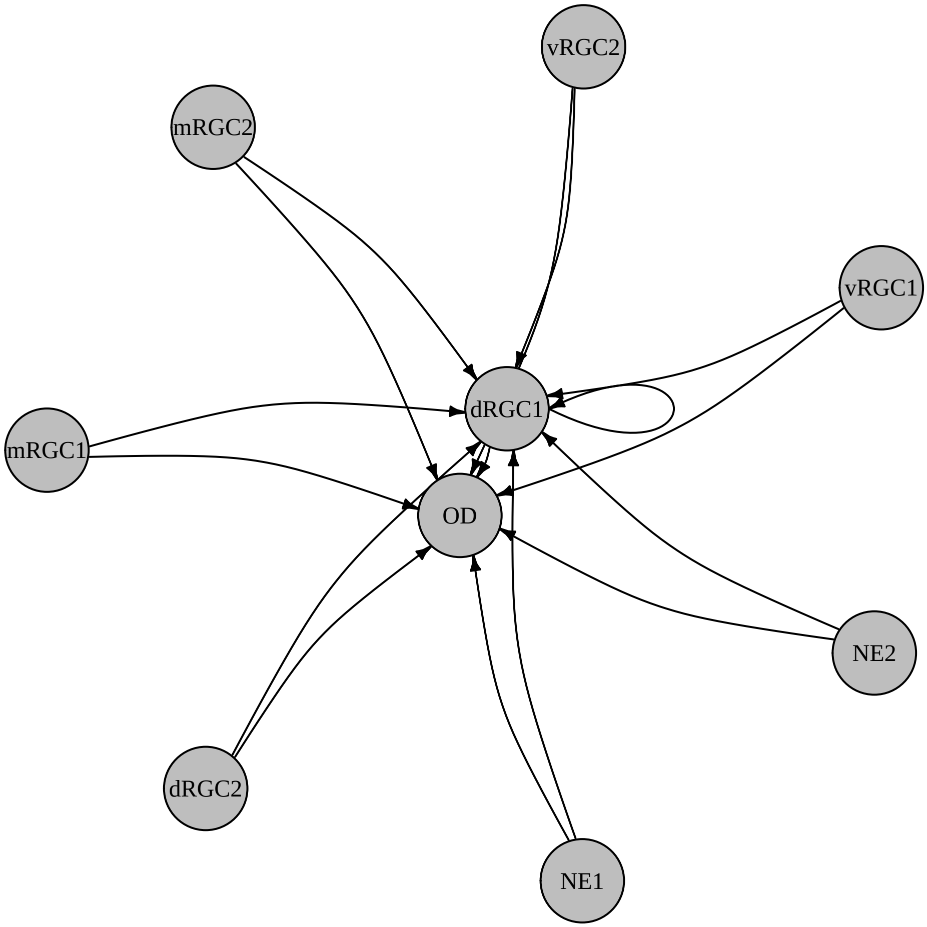

### SD_1L_PBS_uniqintrxs_notch.pdf

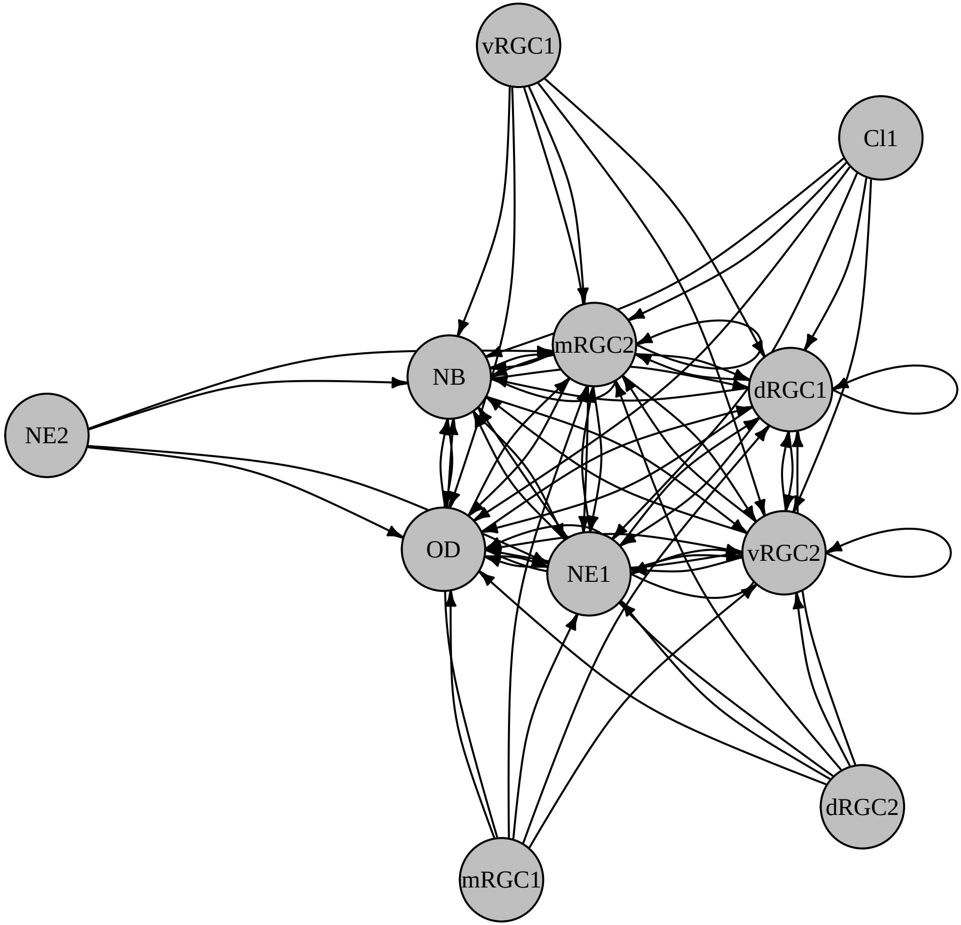

### SD_1L_PBS_uniqintrxs_others_agrn.pdf

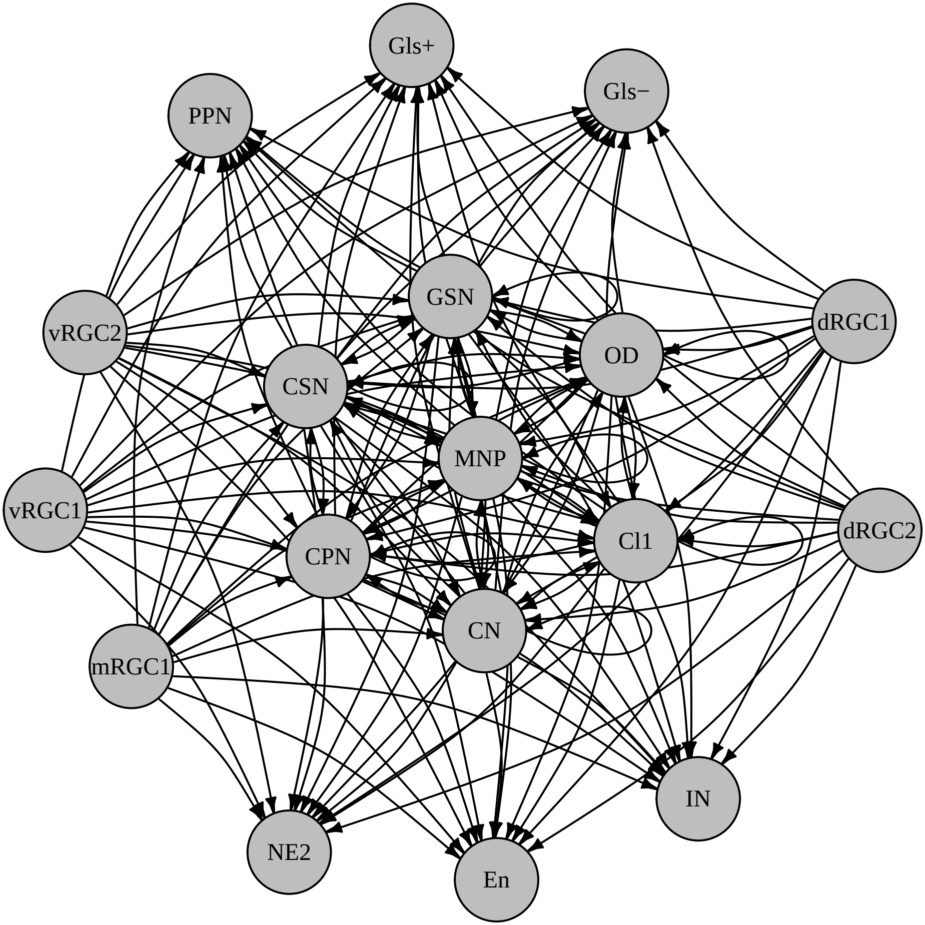

### SD_1L_PBS_uniqintrxs_others_appa.pdf

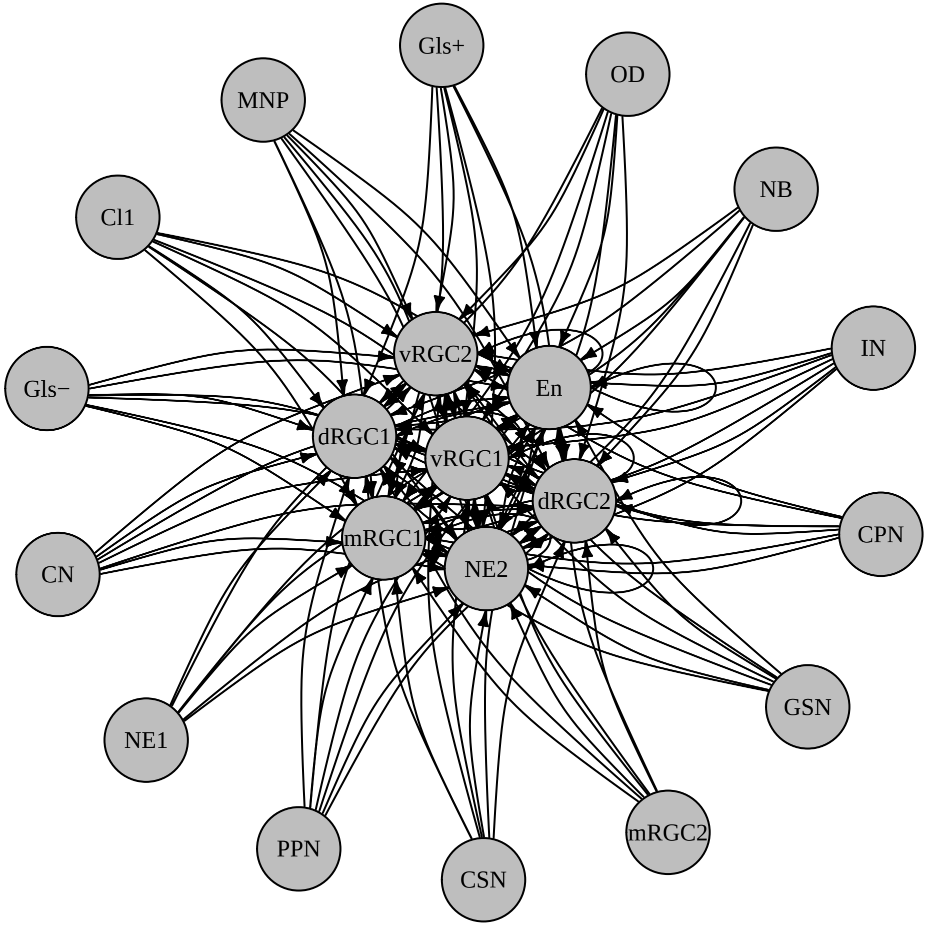

### SD_1L_PBS_uniqintrxs_others_ctgfa.pdf

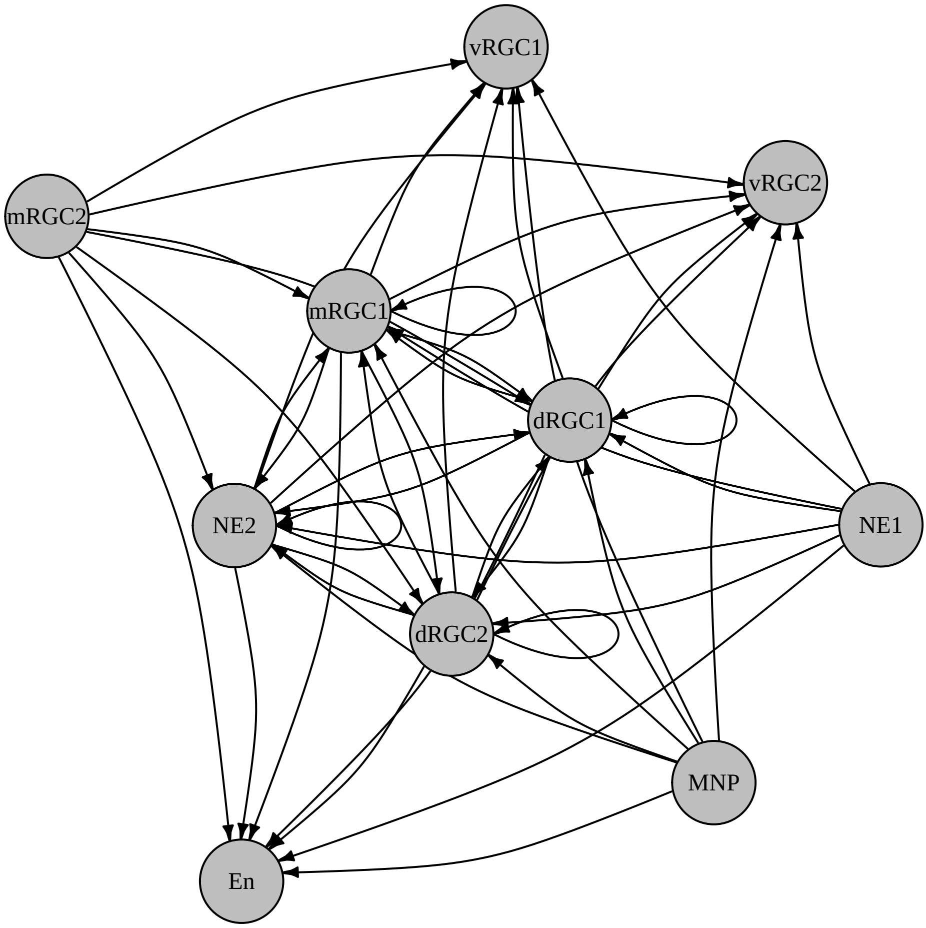

### SD_1L_PBS_uniqintrxs_others_edil.pdf

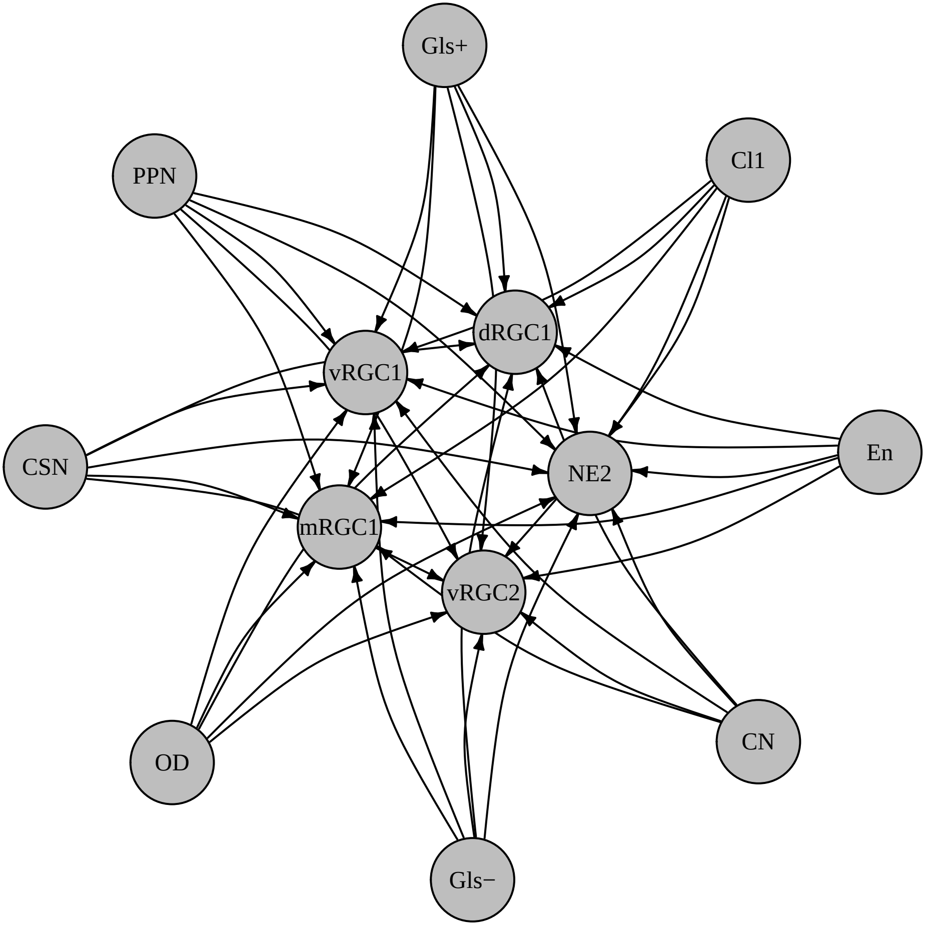

### SD_1L_PBS_uniqintrxs_others_gnai.pdf

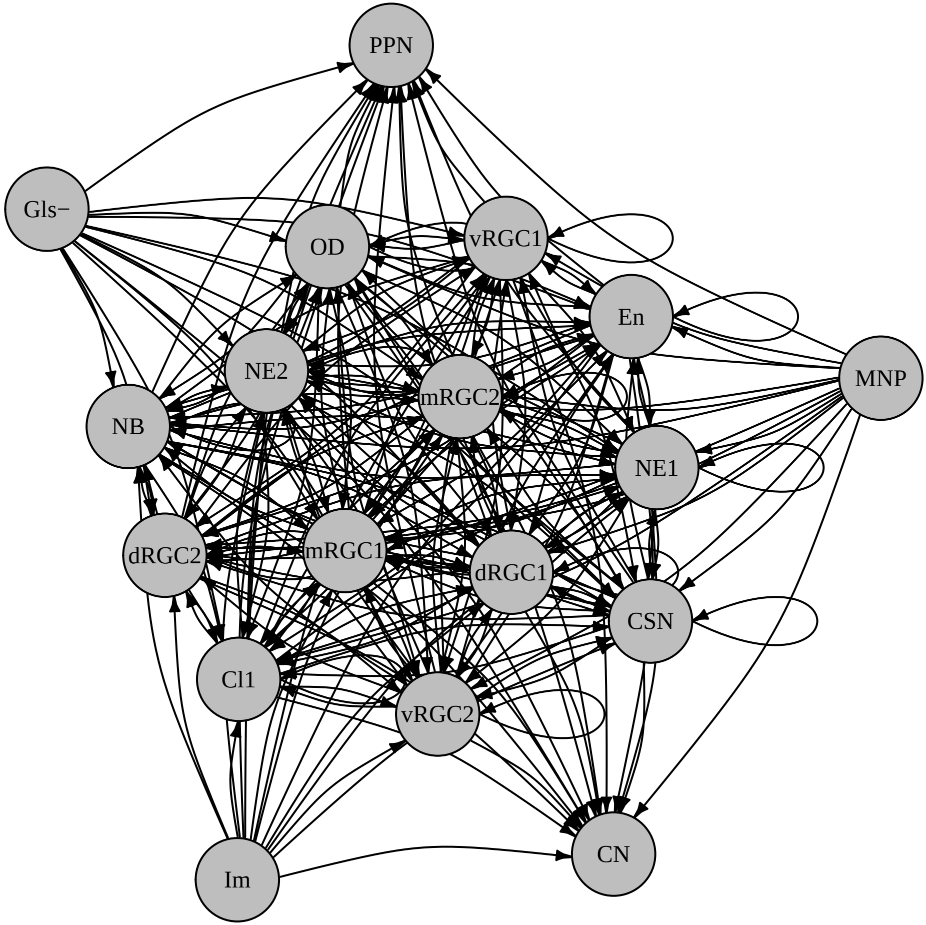

### SD_1L_PBS_uniqintrxs_others_hbegfa.pdf

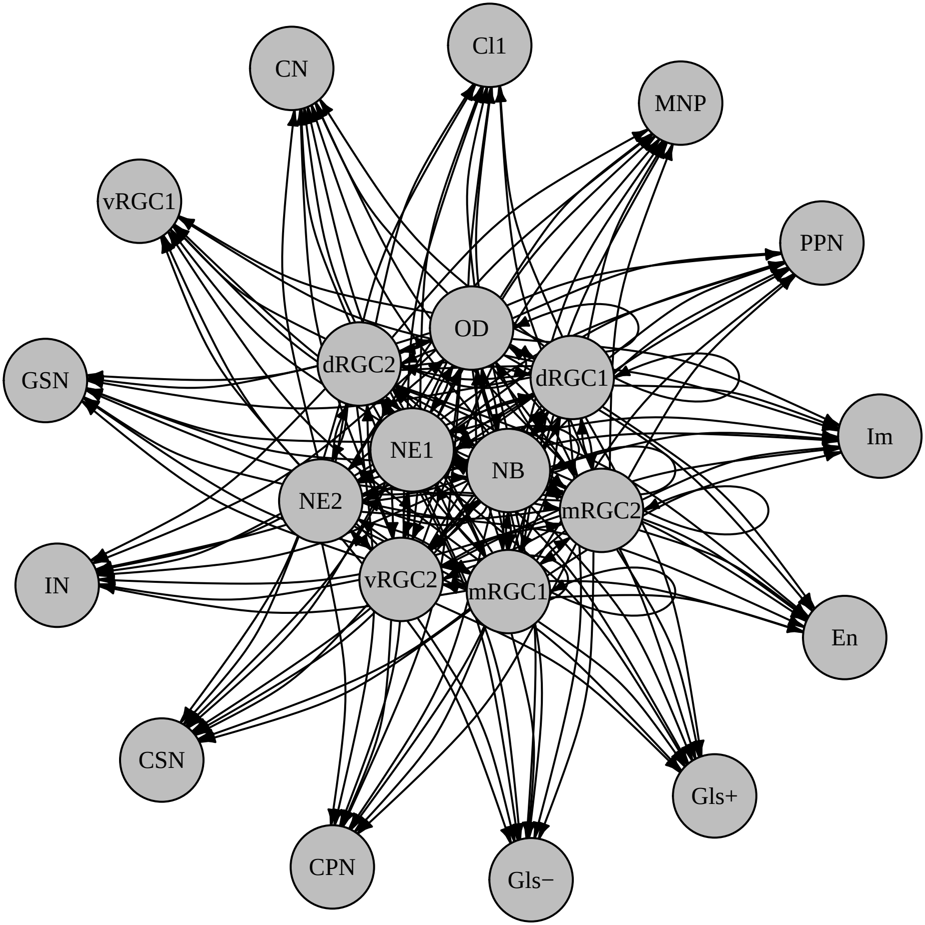

### SD_1L_PBS_uniqintrxs_others_penk_ptn.pdf

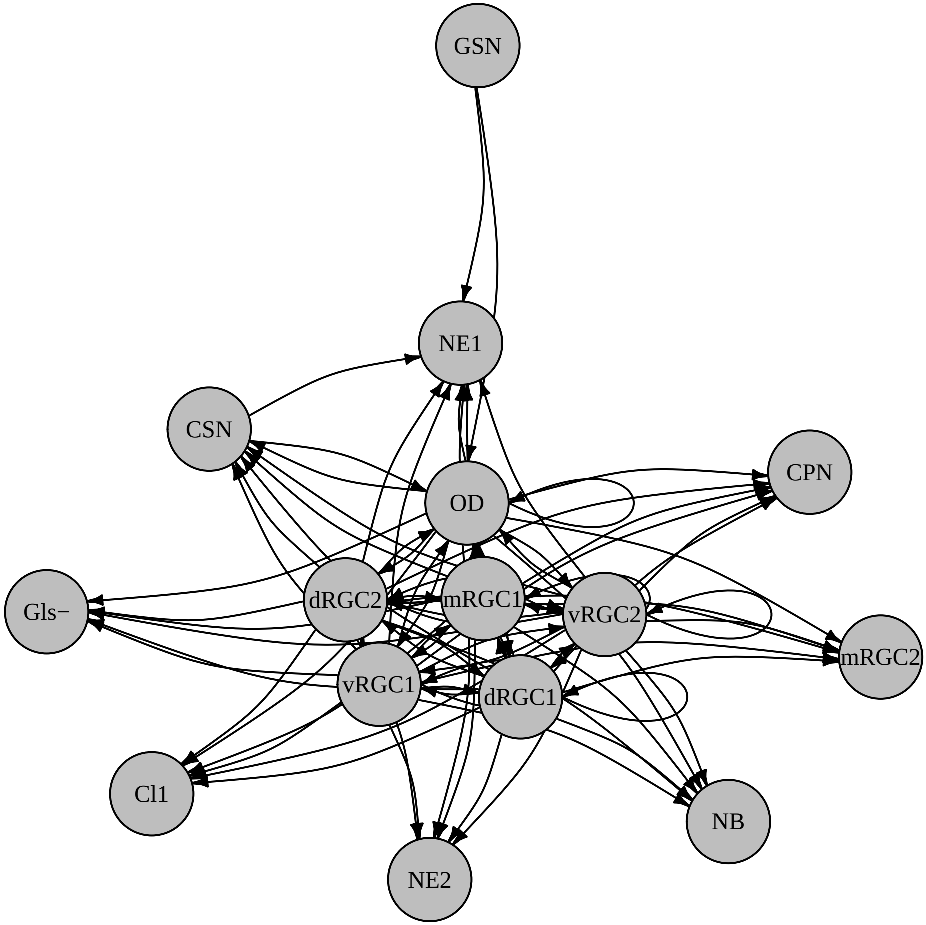

### SD_1L_PBS_uniqintrxs_others_serpine.pdf

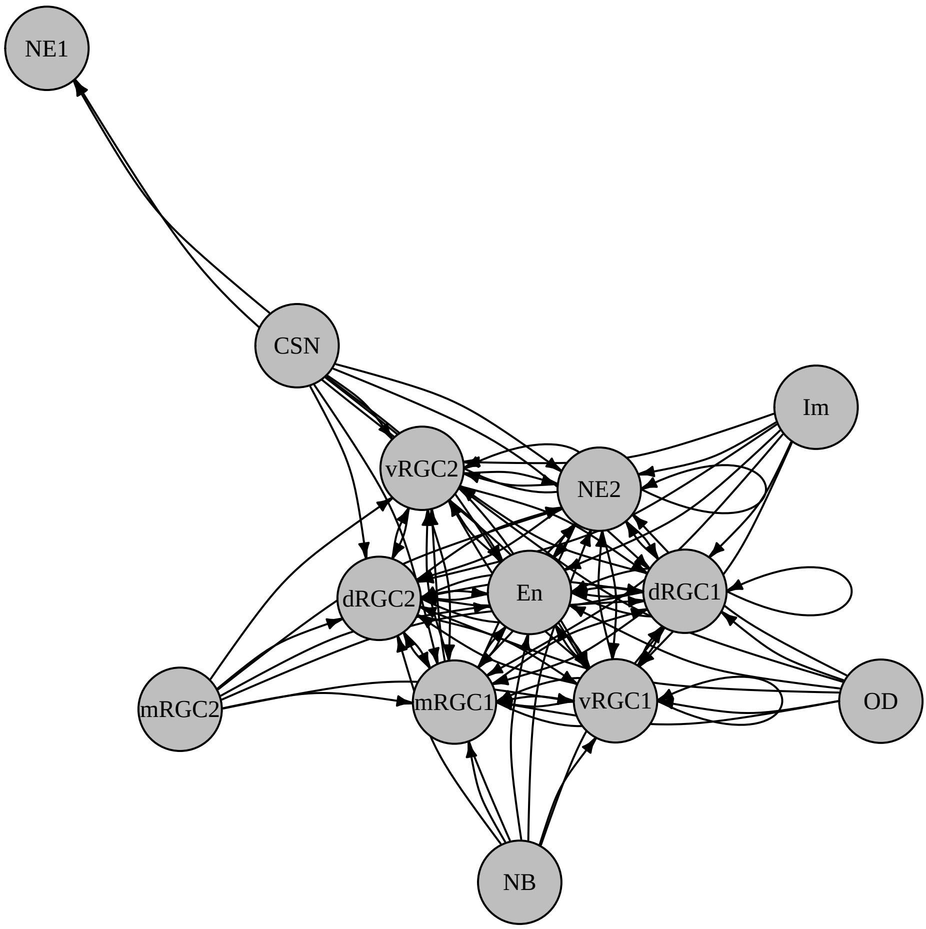

### SD_1L_PBS_uniqintrxs_wnt.pdf

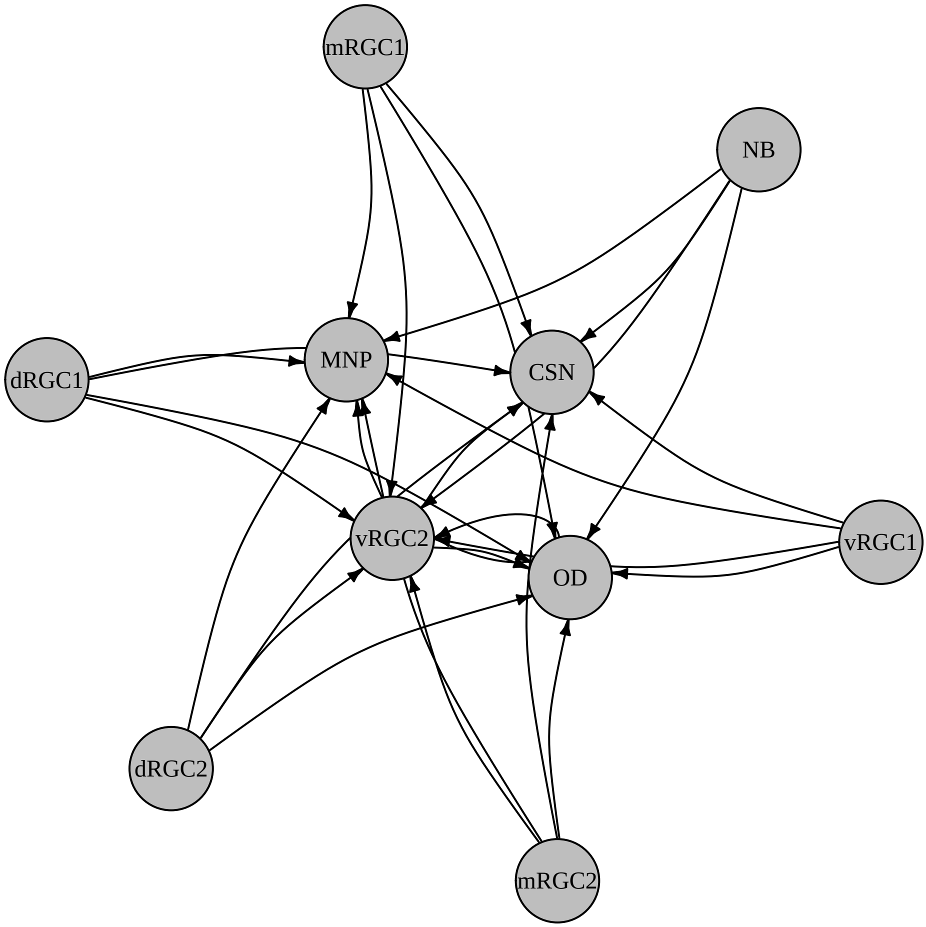

### SD_2F_PBS_and_AB42_MF_Up_part1.pdf

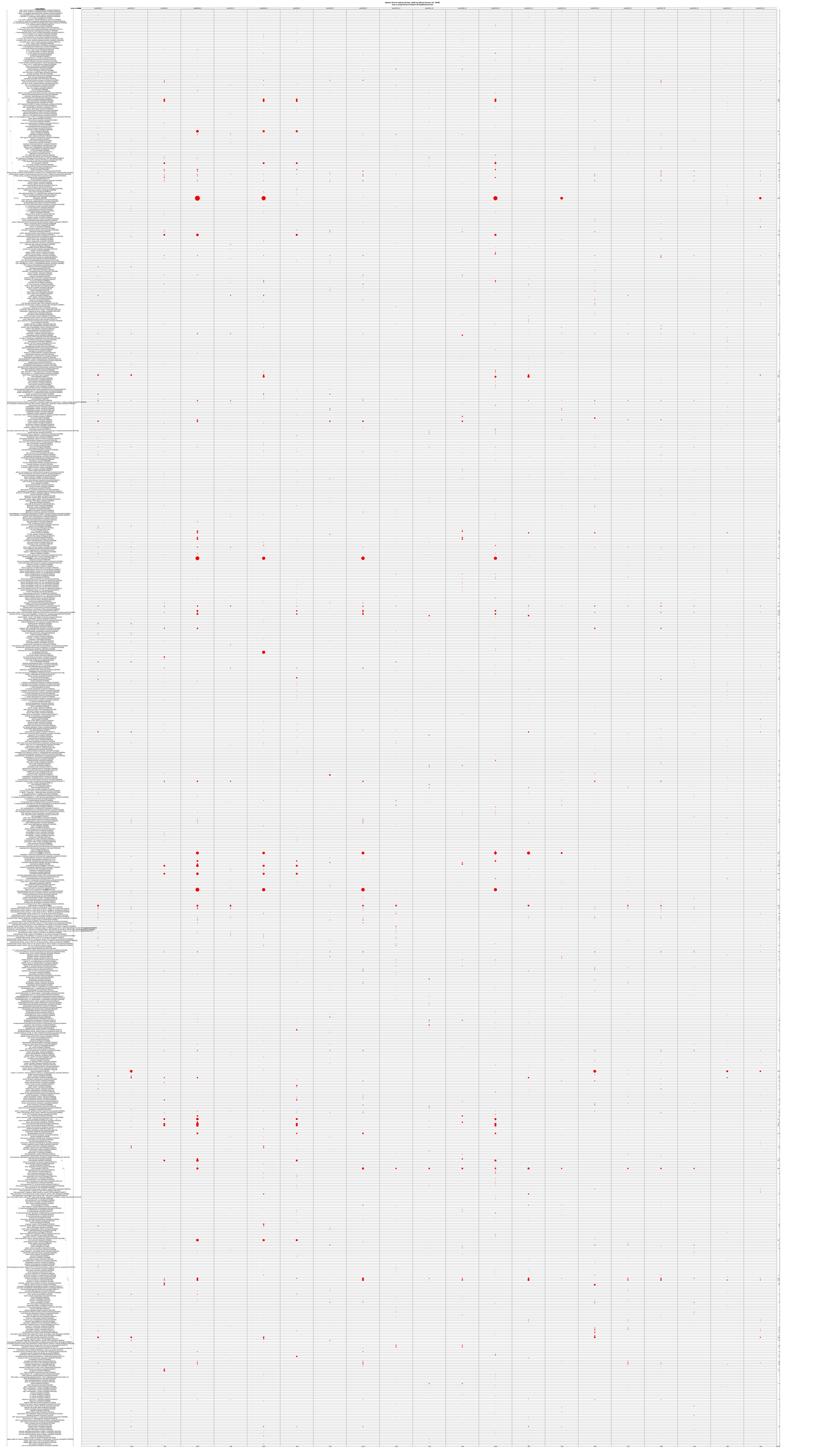

### SD_2G_PBS_and_AB42_MF_Up_uniq_part1.pdf

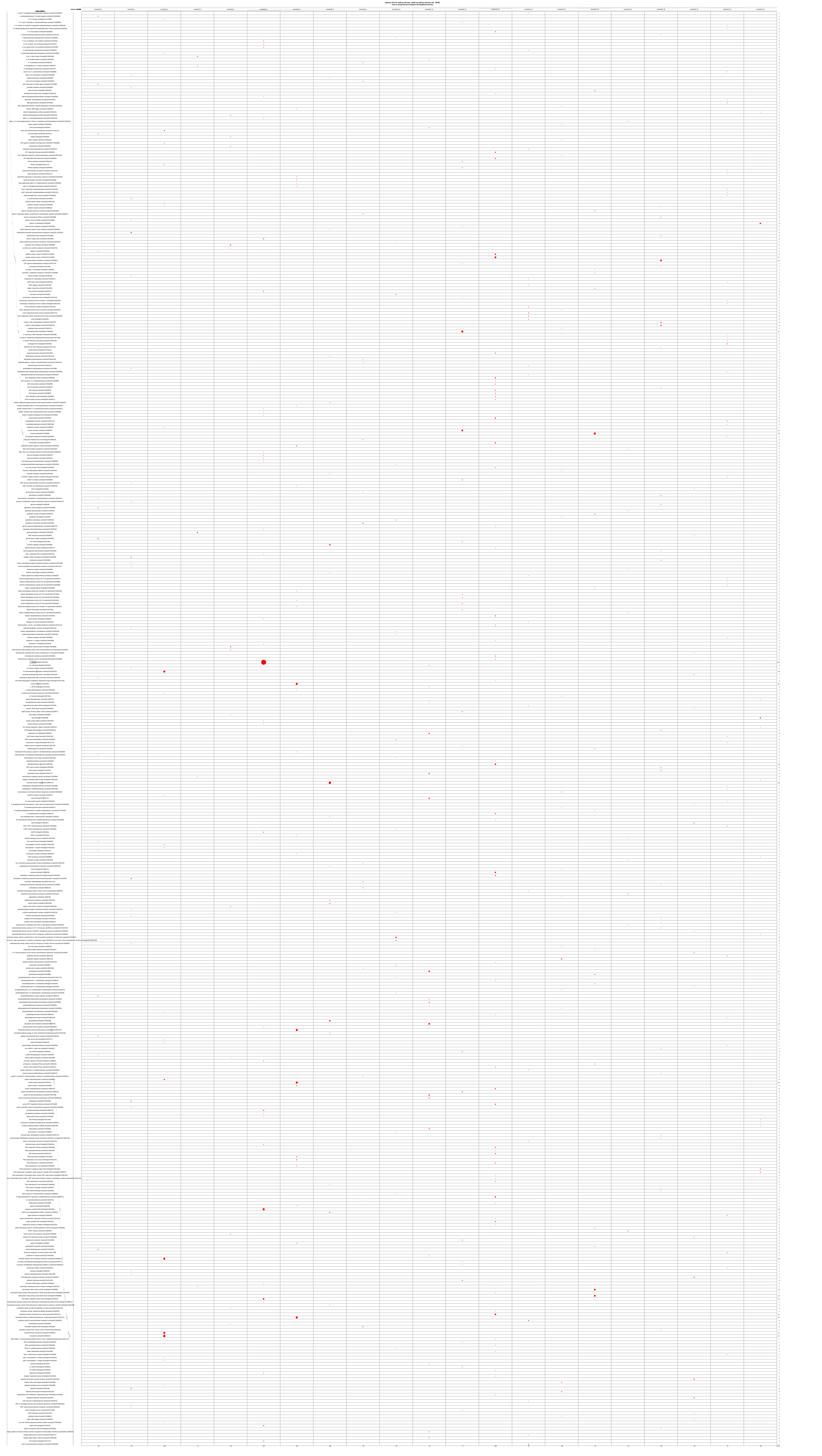

### SD_2K_AB42_vs_PBS_bmp.pdf

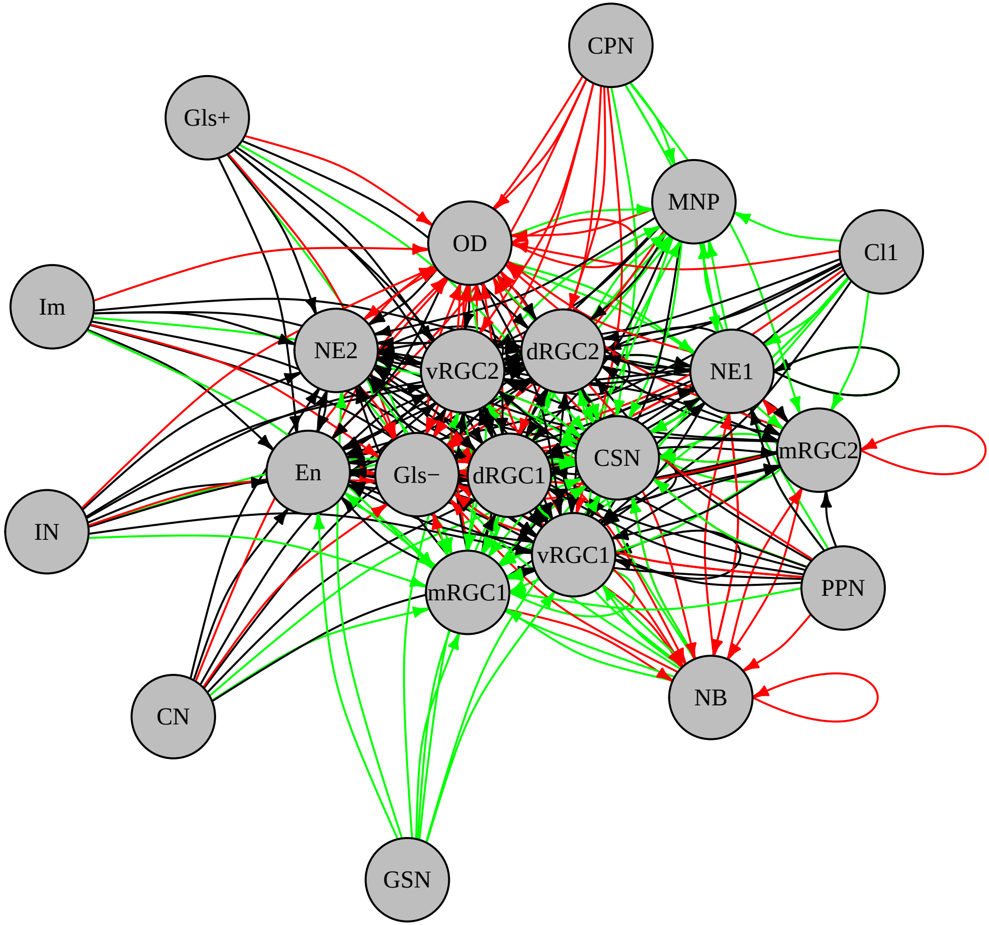

### SD_2K_AB42_vs_PBS_chemokine.pdf

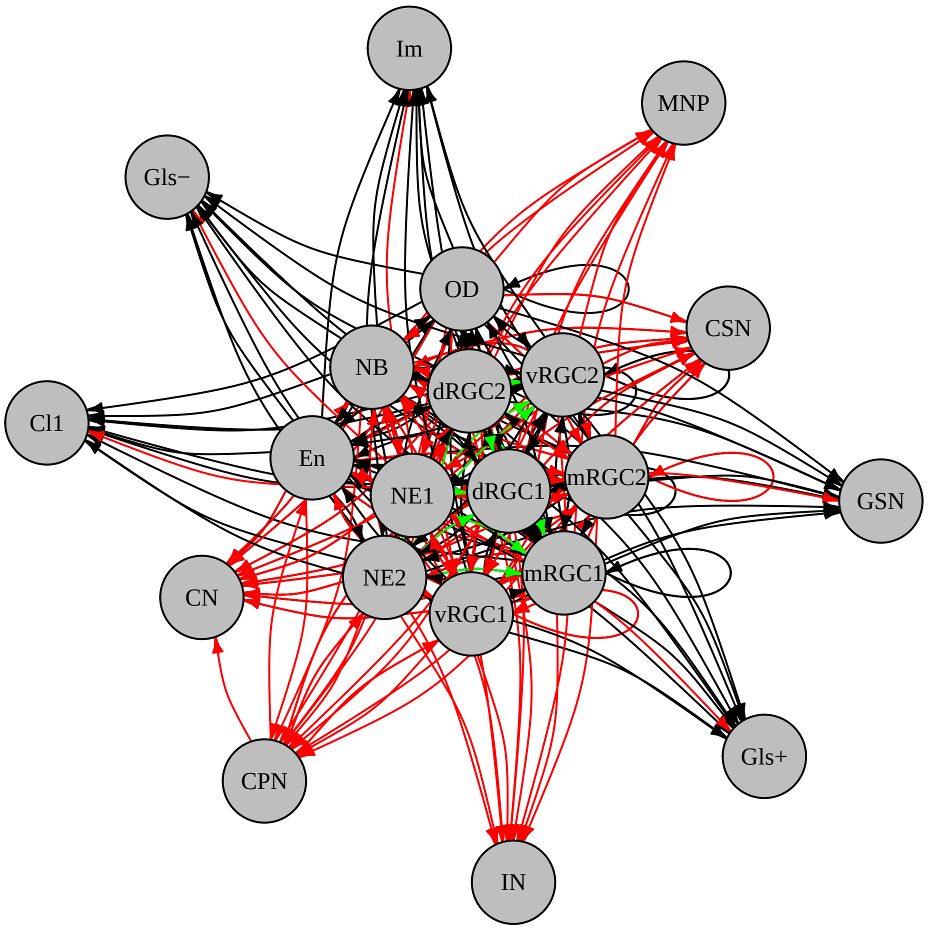

### SD_2K_AB42_vs_PBS_epha.pdf

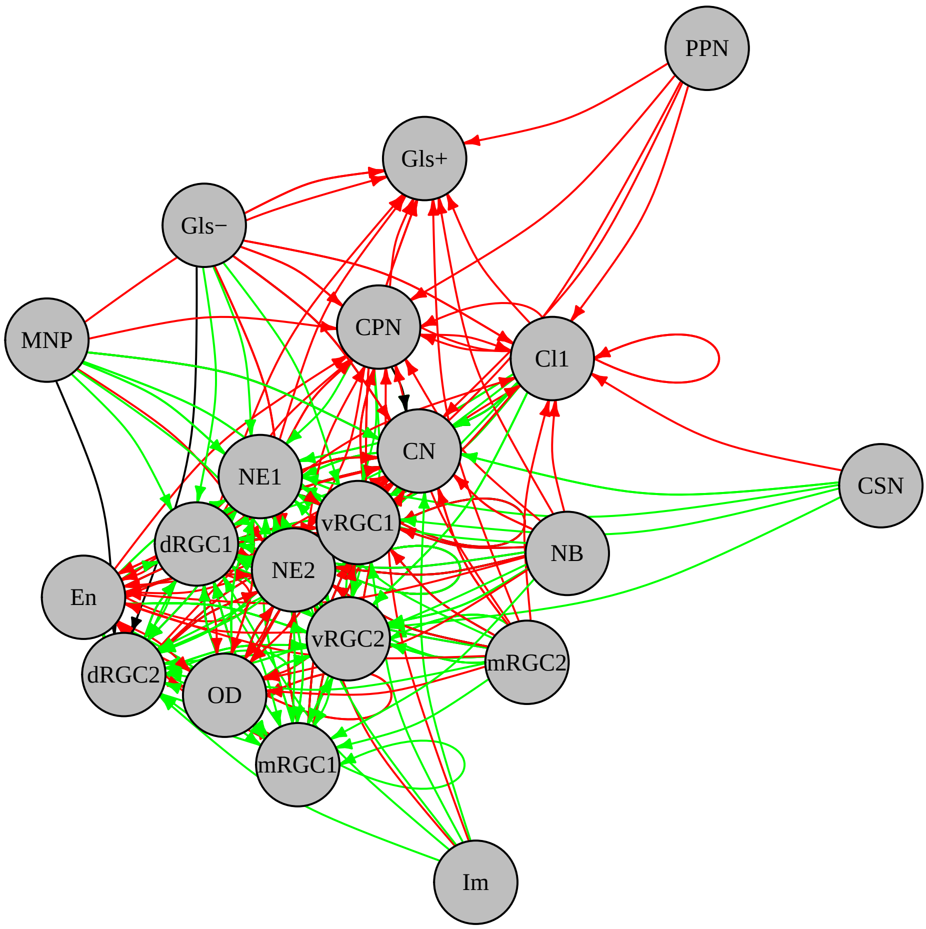

### SD_2K_AB42_vs_PBS_fgf.pdf

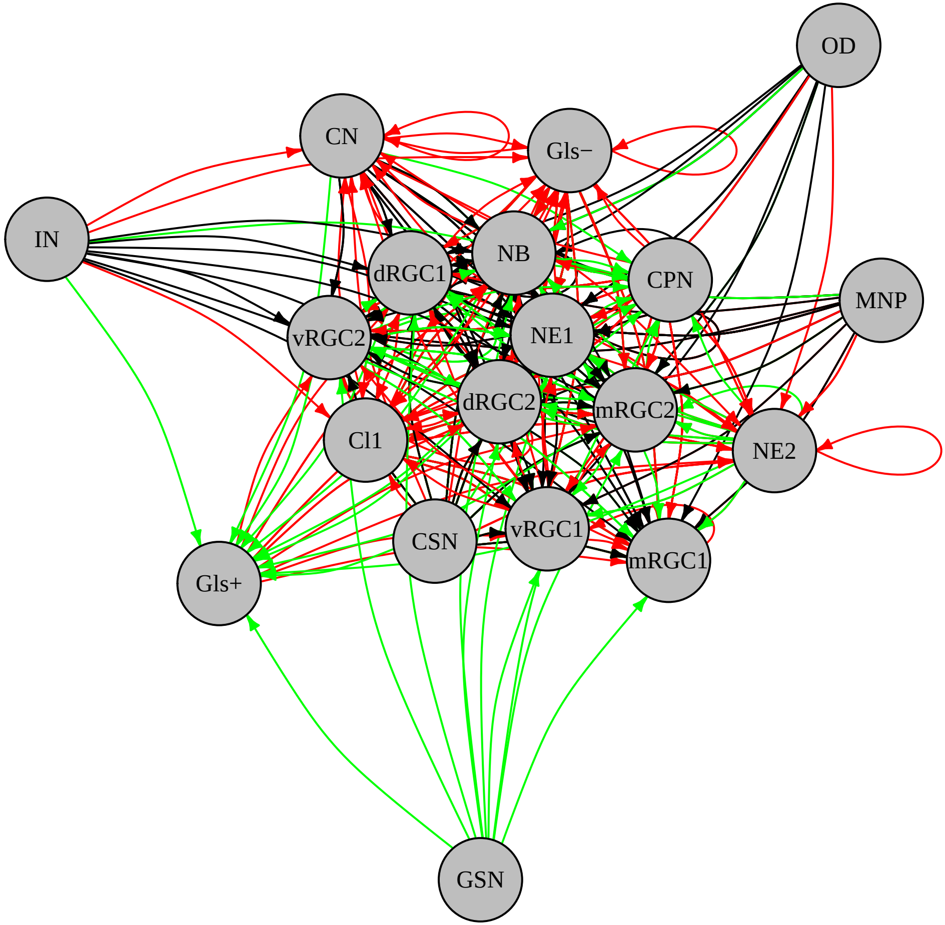

### SD_2K_AB42_vs_PBS_igf.pdf

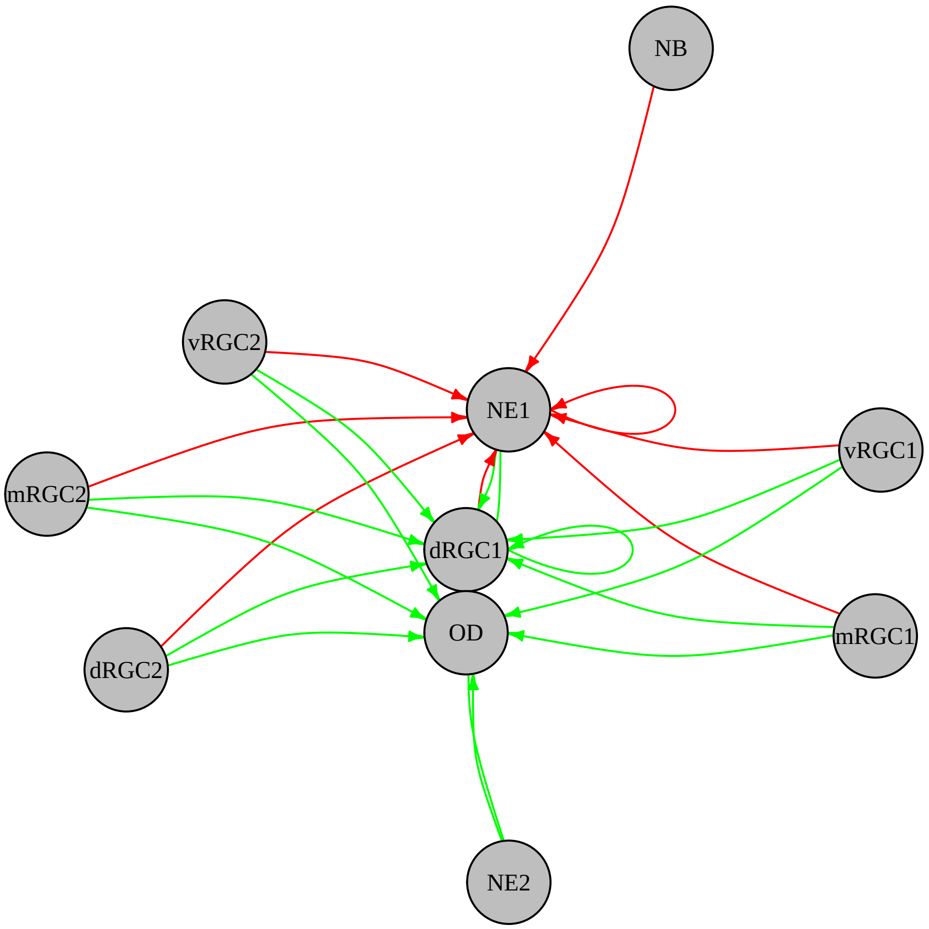

### SD_2K_AB42_vs_PBS_notch.pdf

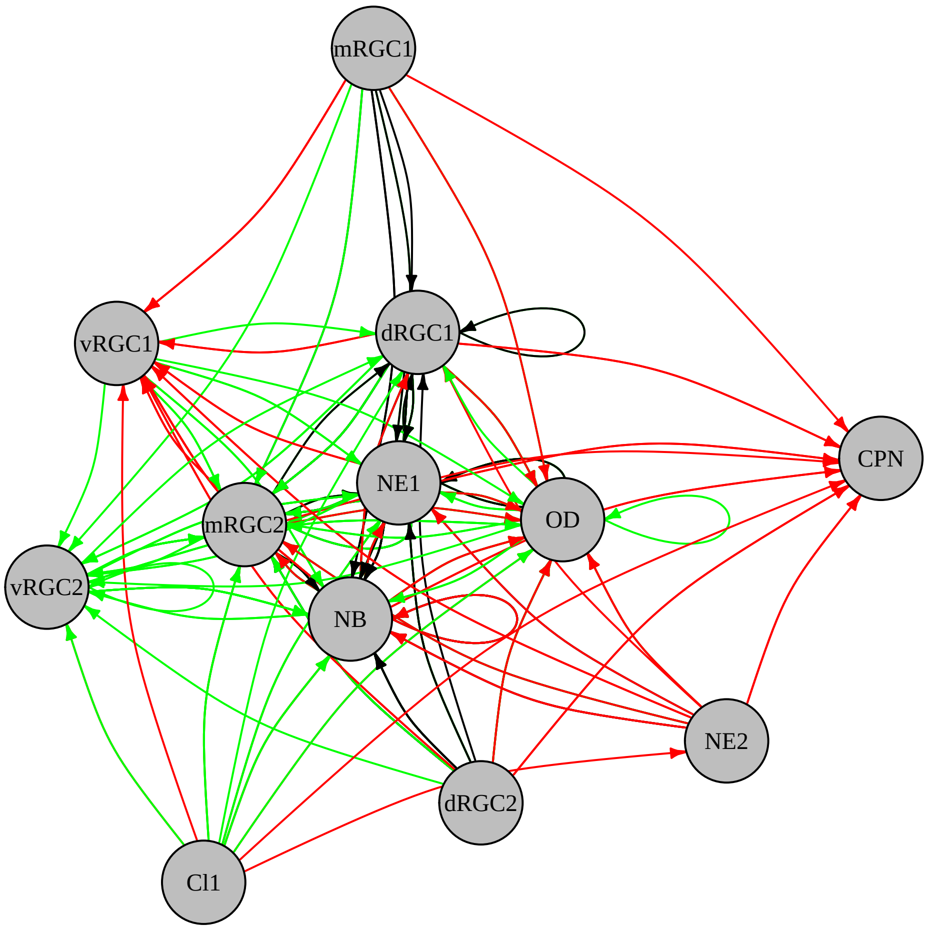

### SD_2K_AB42_vs_PBS_others_agrn.pdf

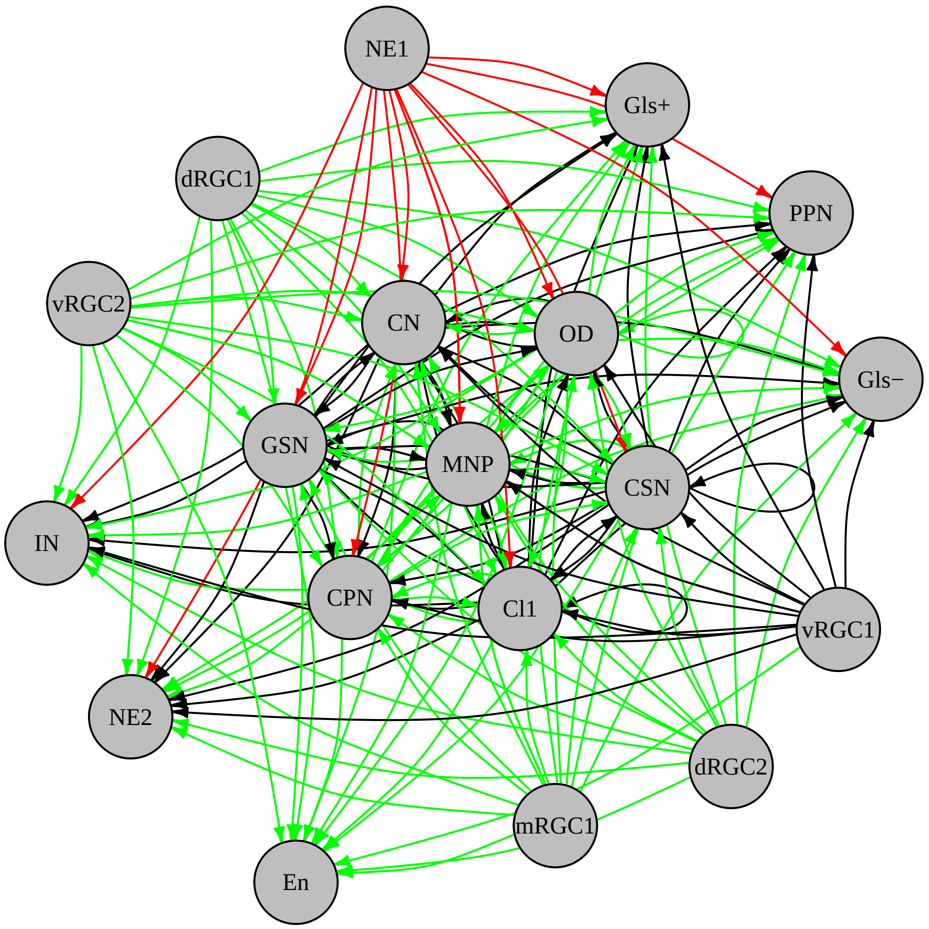

### SD_2K_AB42_vs_PBS_others_appa.pdf

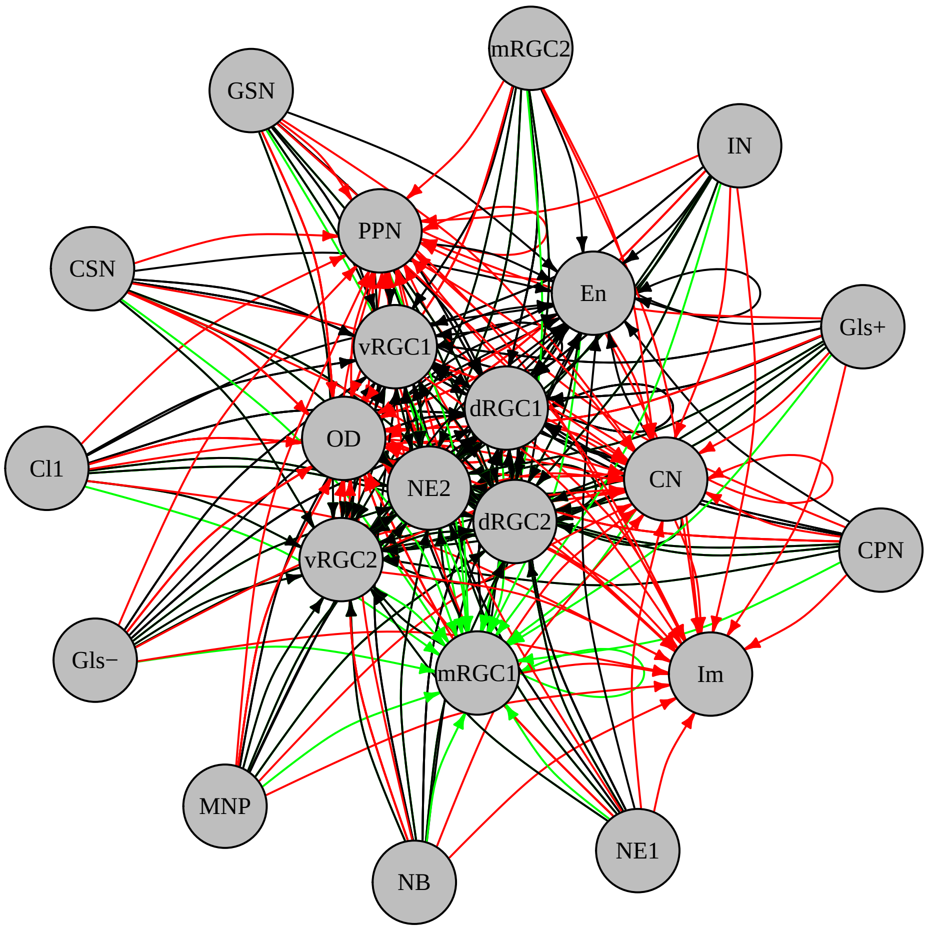

### SD_2K_AB42_vs_PBS_others_ctgfa.pdf

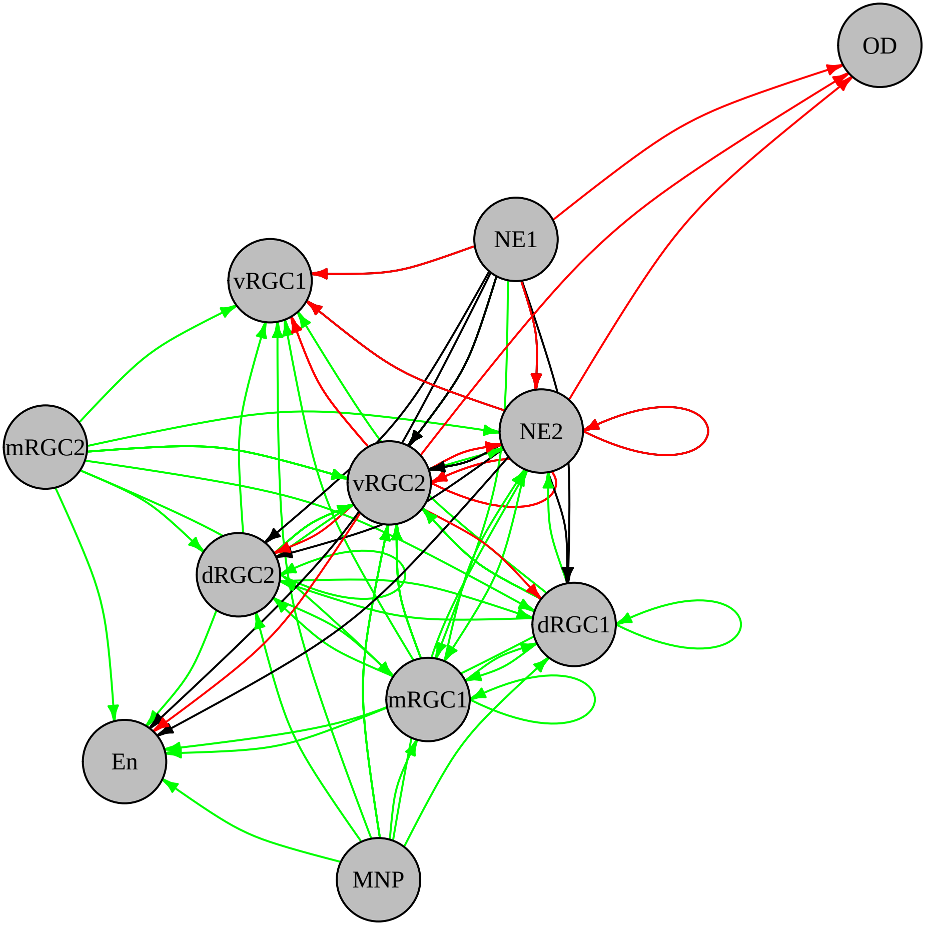

### SD_2K_AB42_vs_PBS_others_edil.pdf

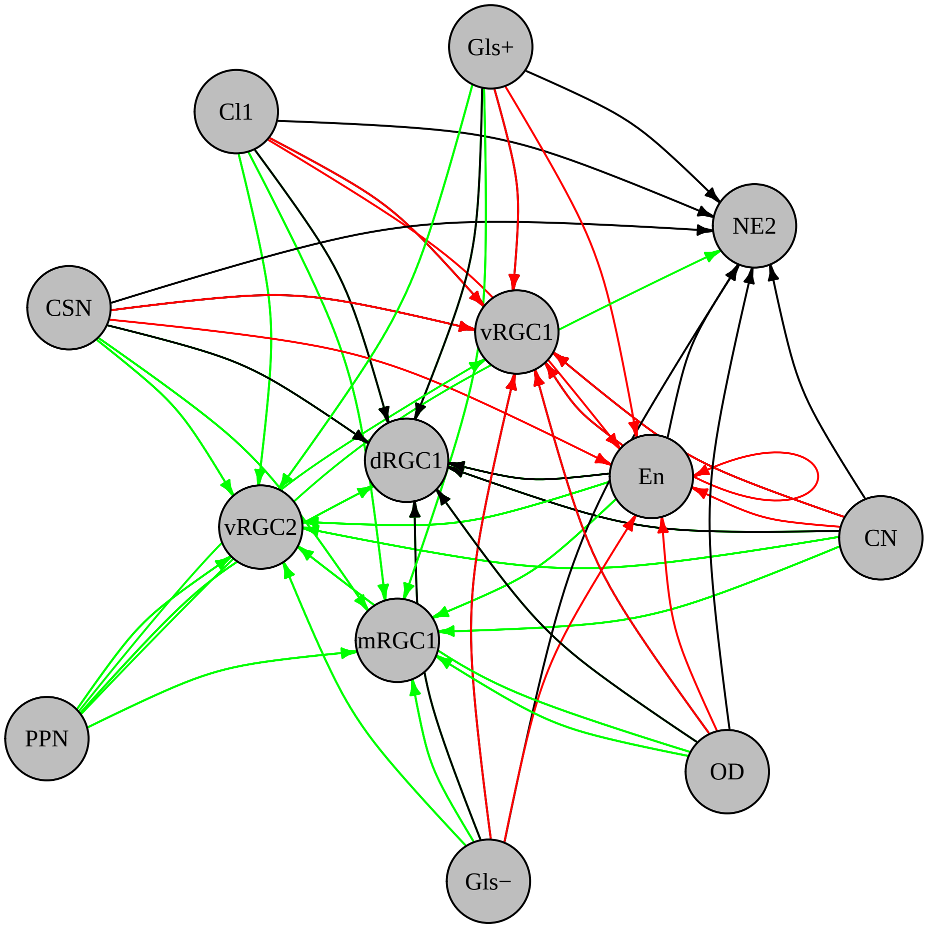

### SD_2K_AB42_vs_PBS_others_gnai.pdf

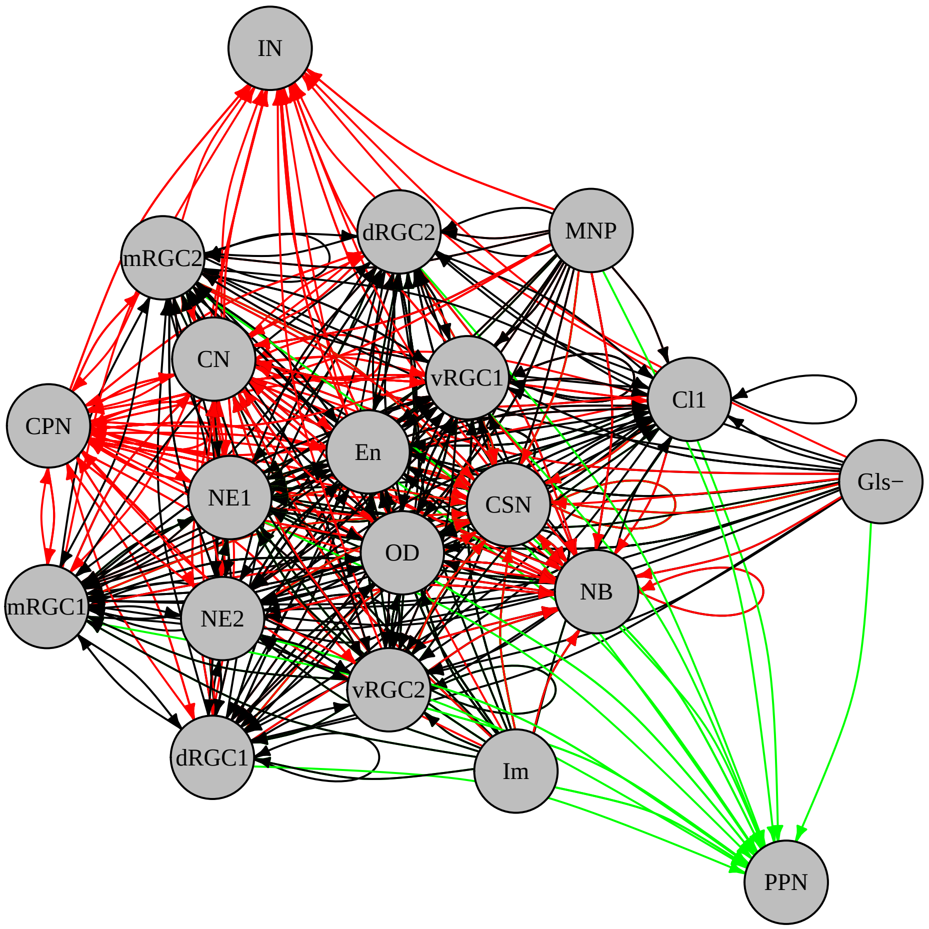

### SD_2K_AB42_vs_PBS_others_hbegfa.pdf

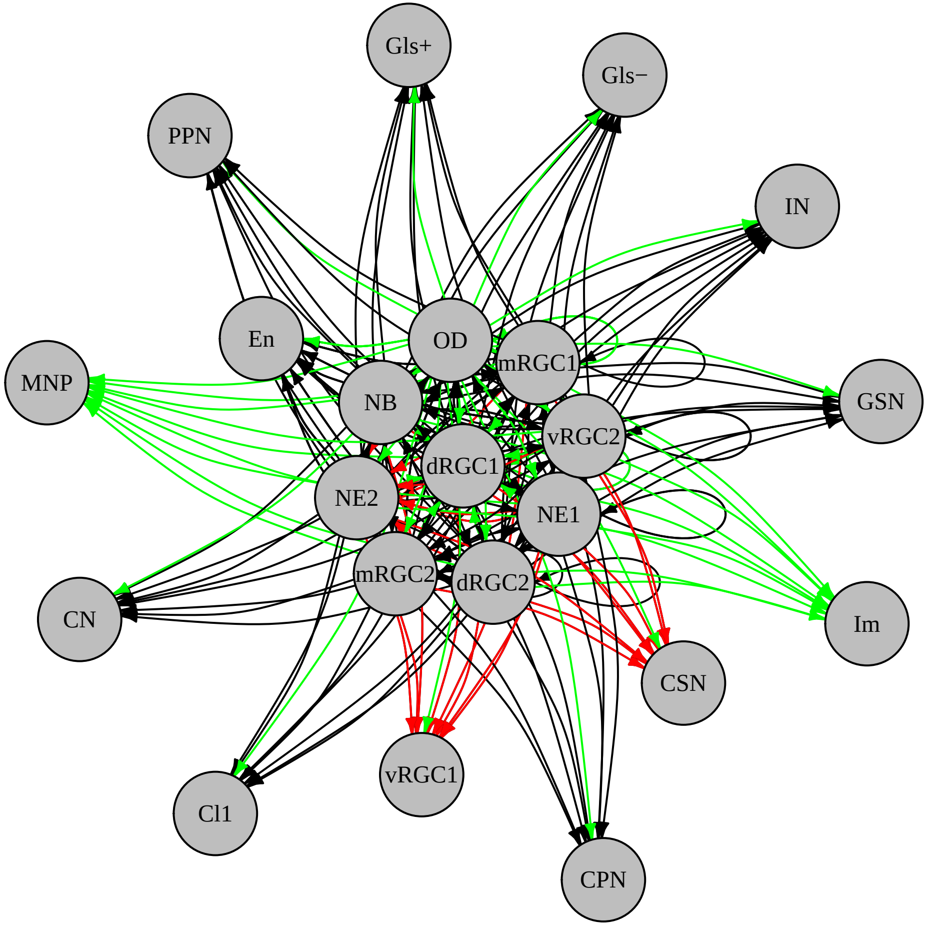

### SD_2K_AB42_vs_PBS_others_penk_ptn.pdf

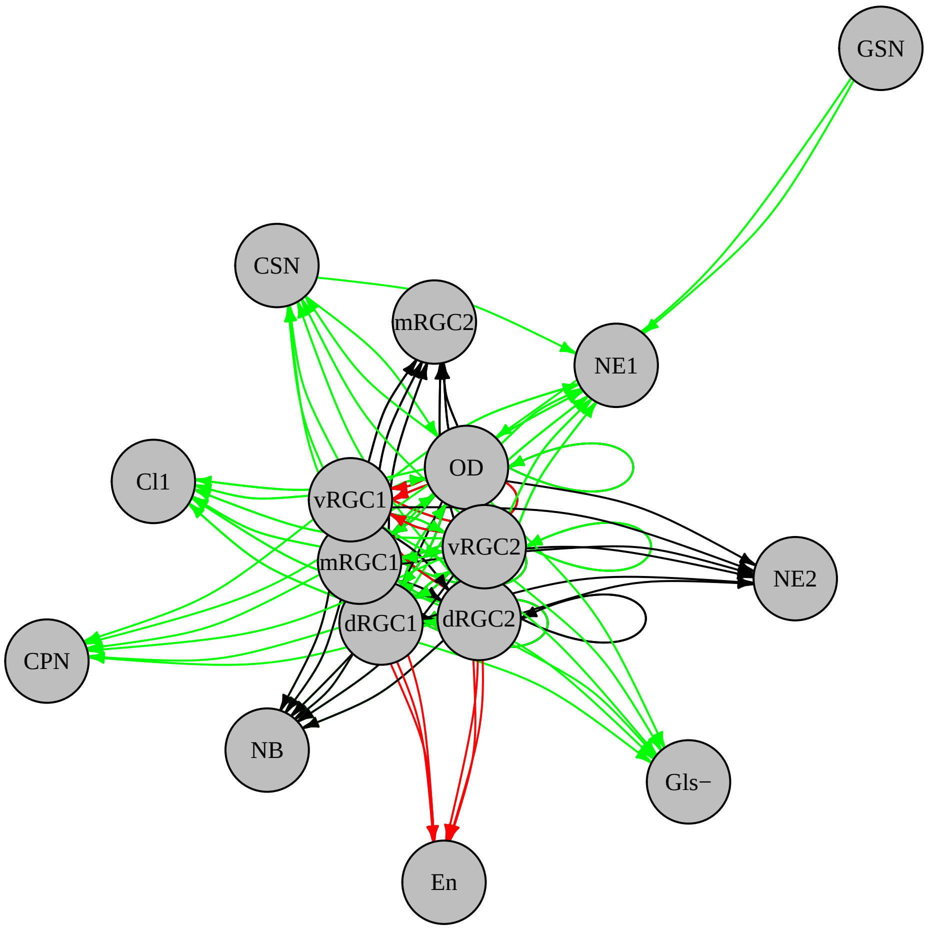

### SD_2K_AB42_vs_PBS_others_serpine.pdf

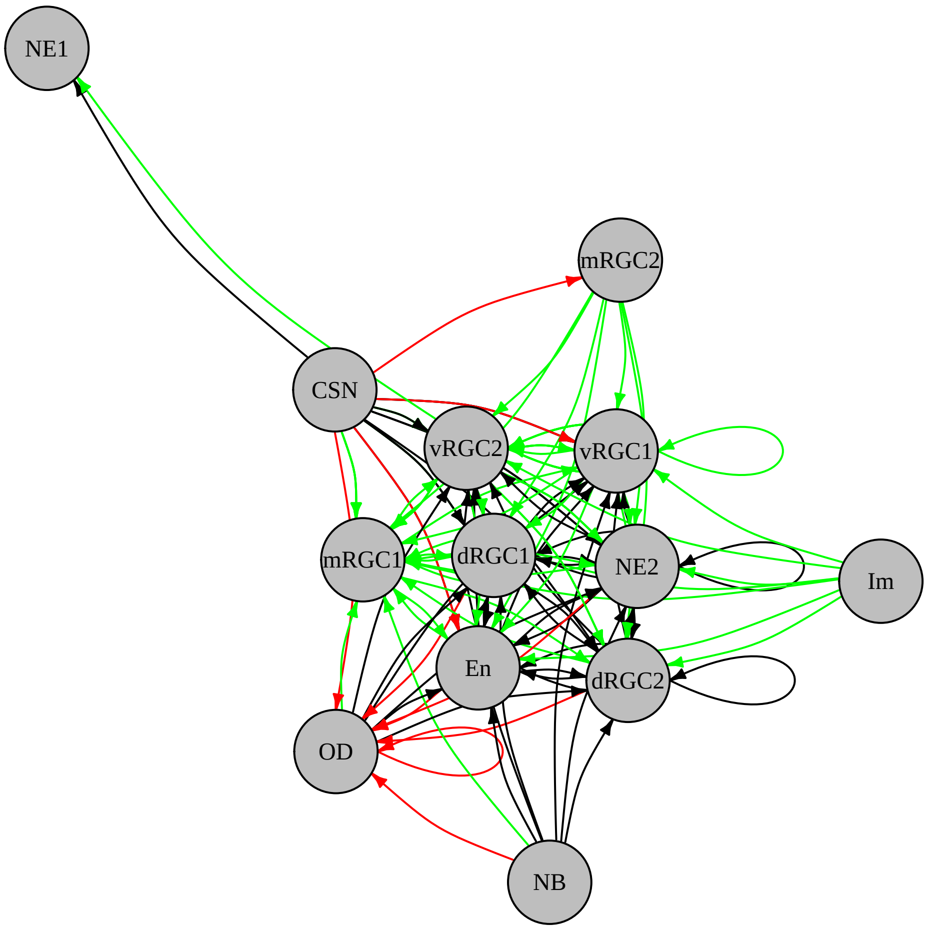

### SD_2K_AB42_vs_PBS_wnt.pdf

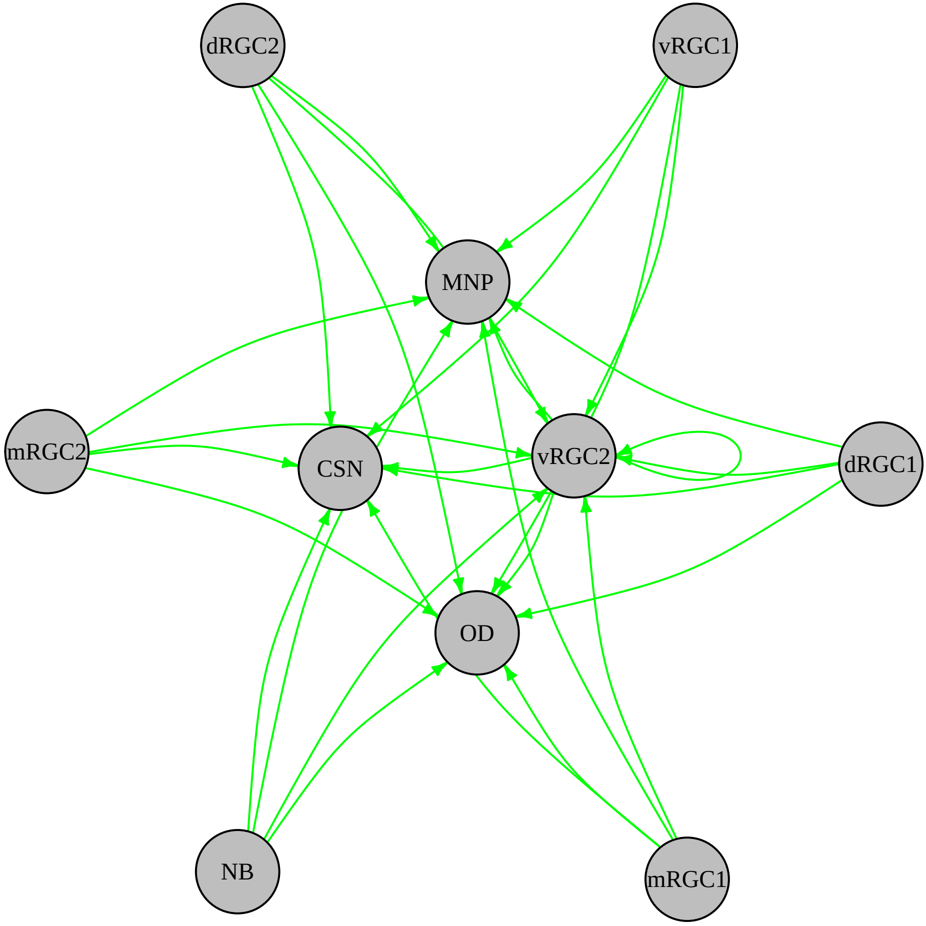

### SD_3C_PBS_and_IL4_BP_Up_part2.pdf

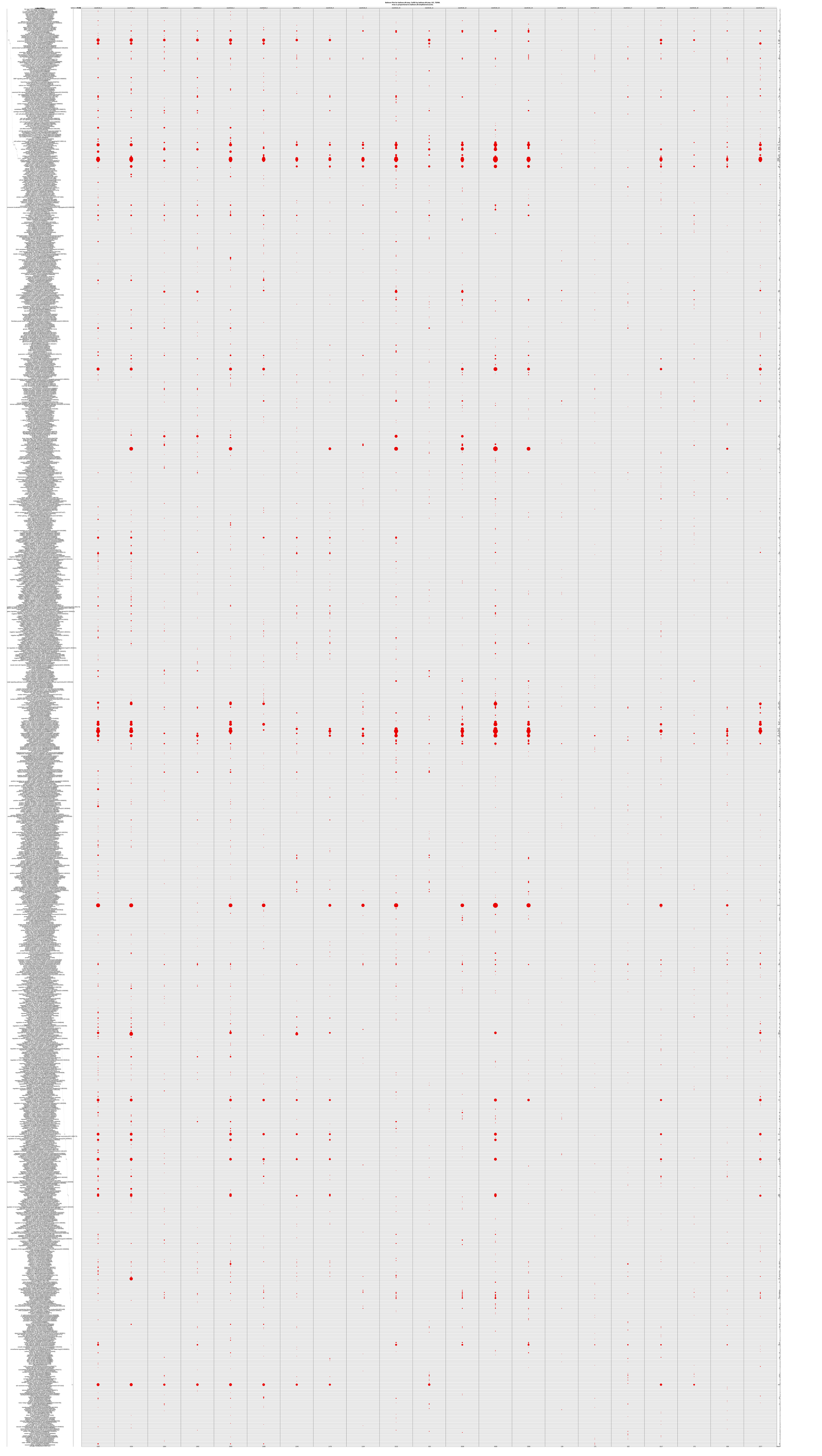

### SD_3D_PBS_and_IL4_BP_Up_uniq_part1.pdf

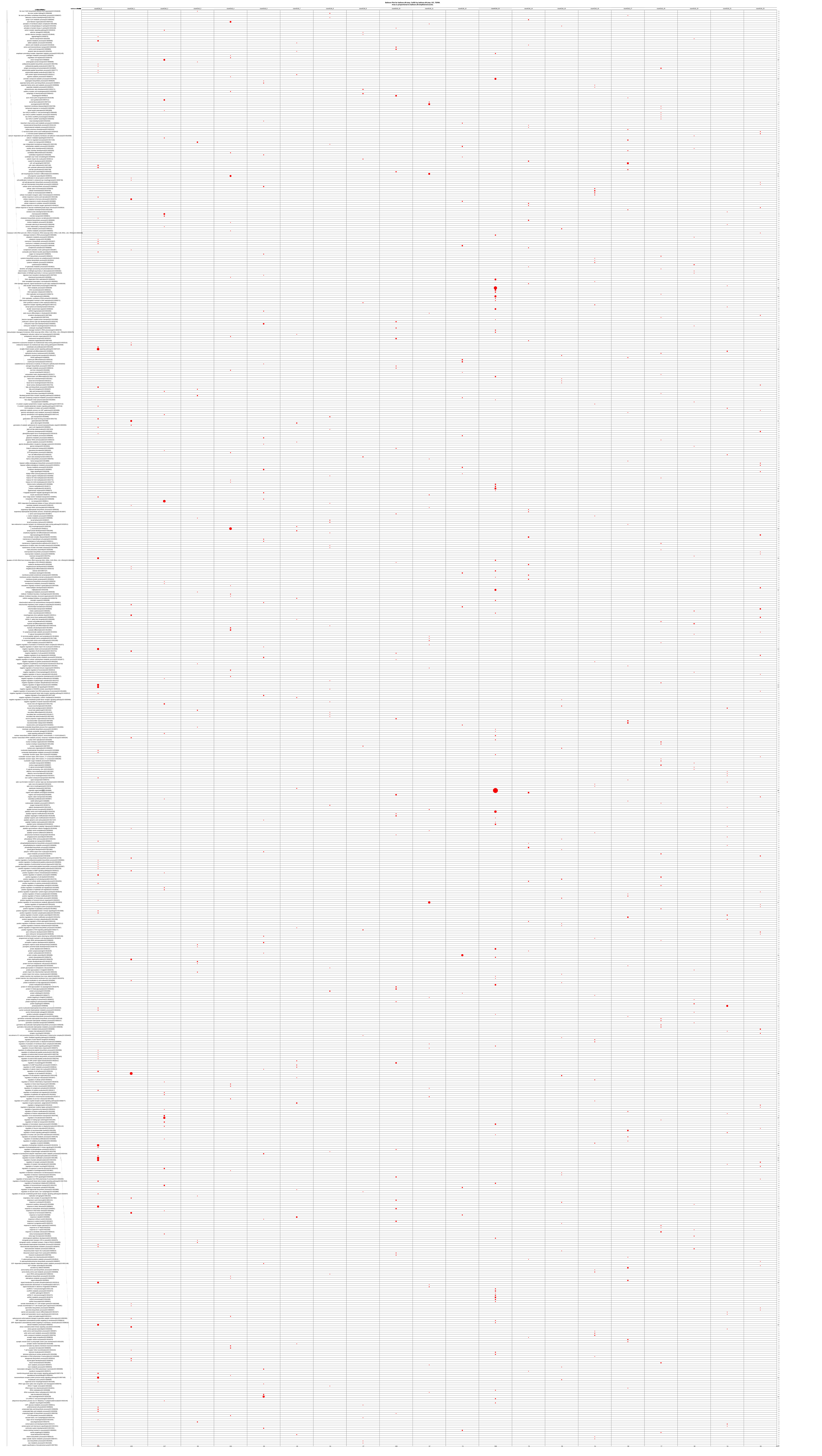

### SD_3F_PBS_and_IL4_MF_Up_uniq_part1.pdf

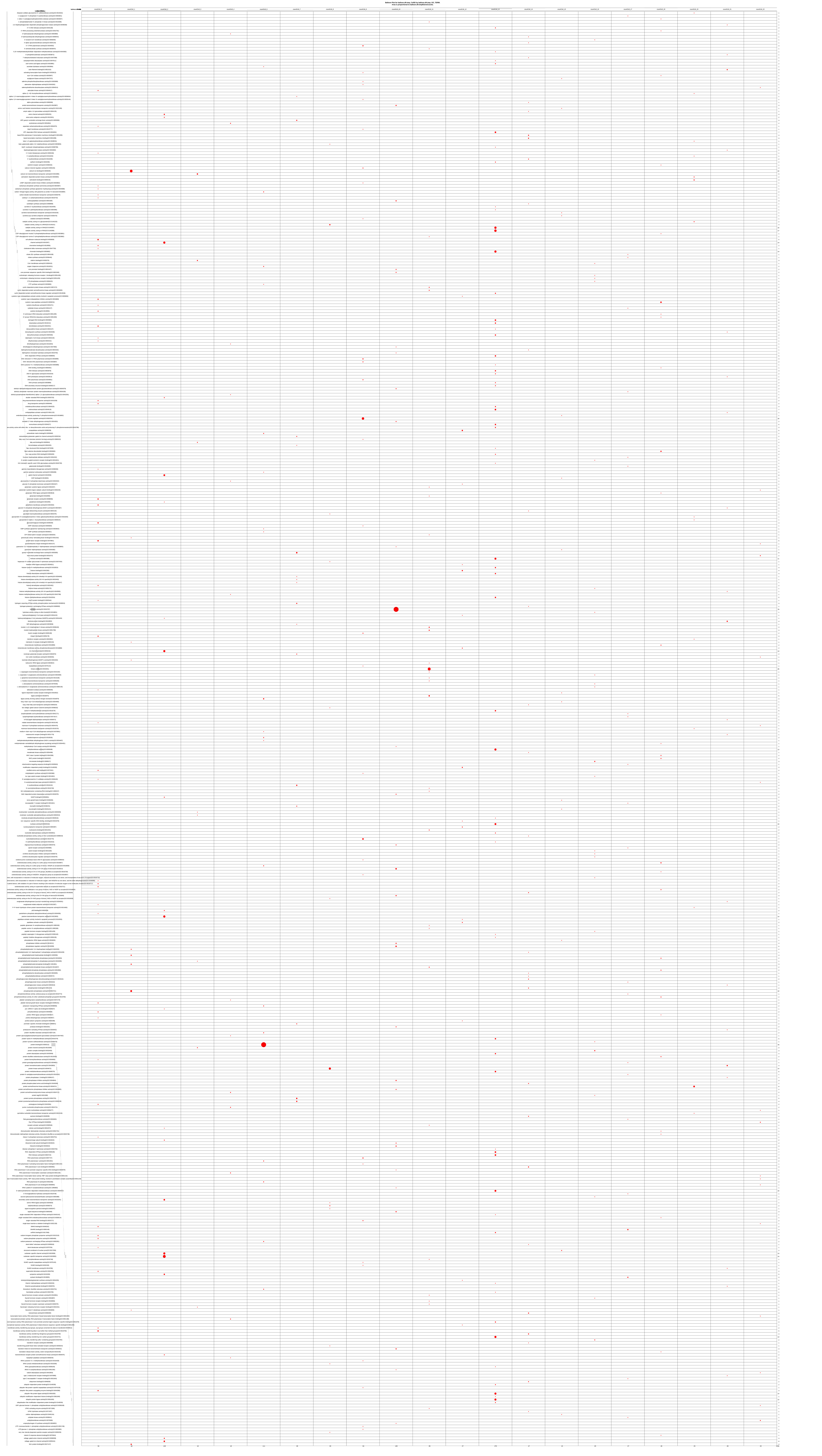

### SD_3L_IL4_vs_PBS_bmp.pdf

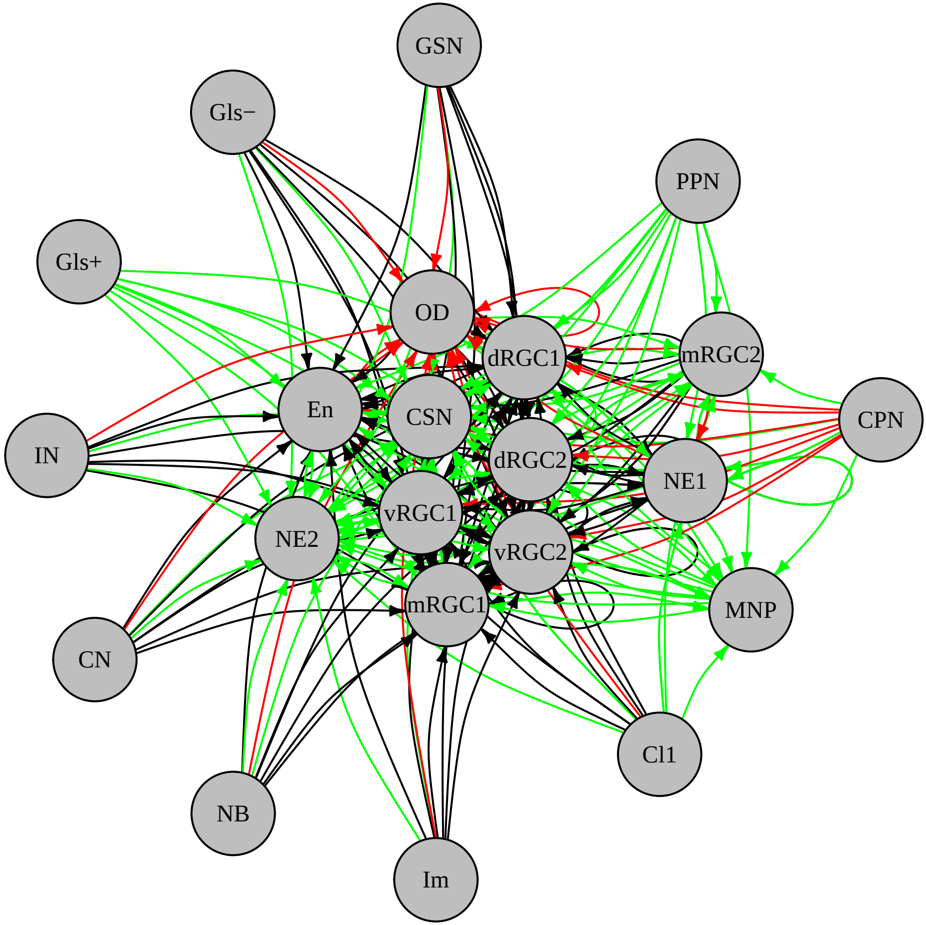

### SD_3L_IL4_vs_PBS_chemokine.pdf

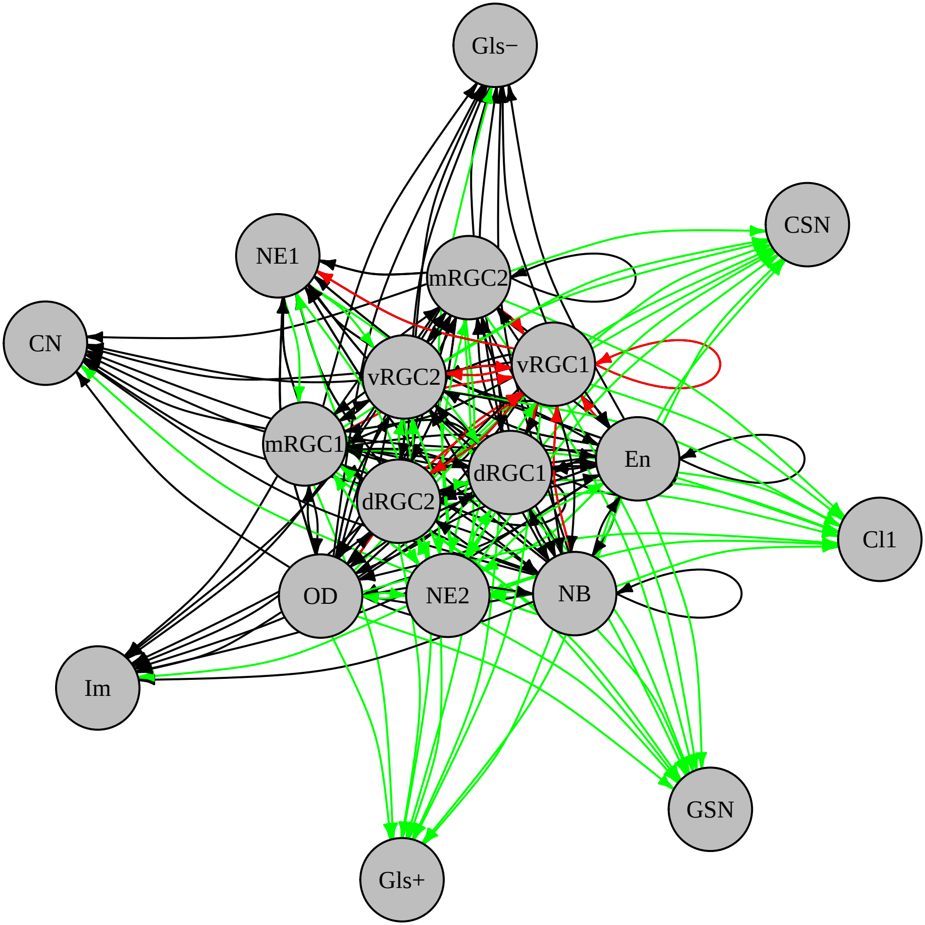

### SD_3L_IL4_vs_PBS_epha.pdf

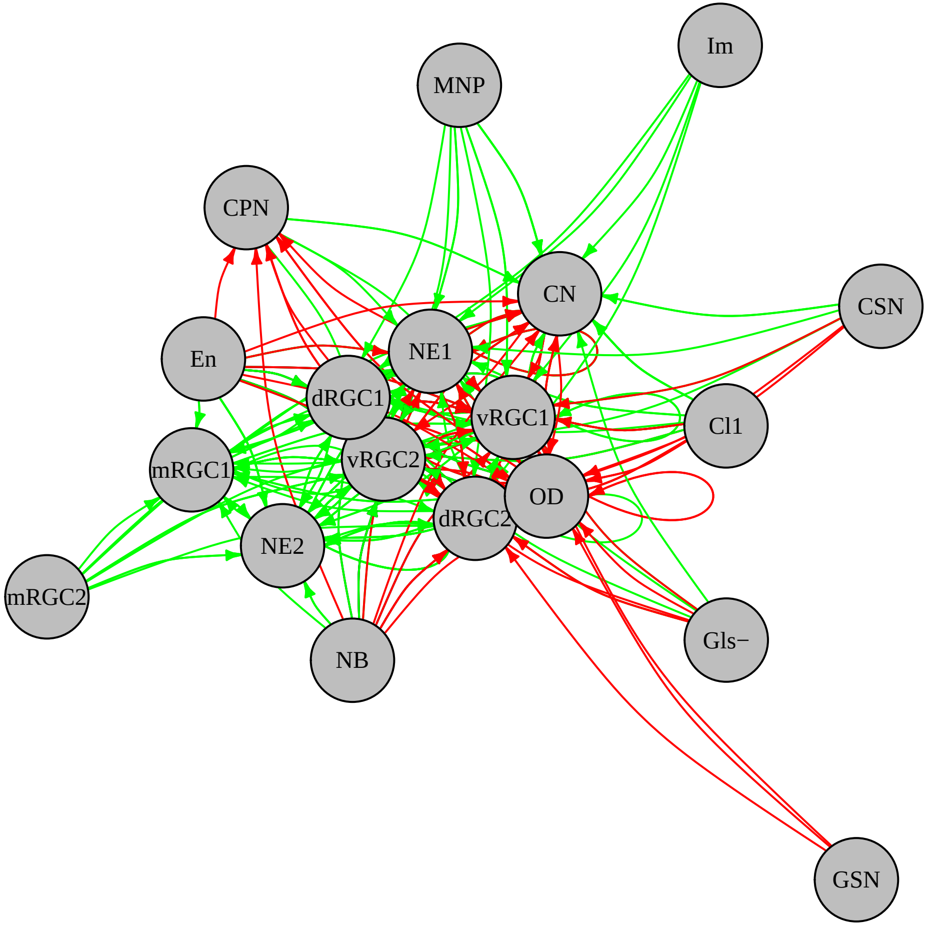

### SD_3L_IL4_vs_PBS_fgf.pdf

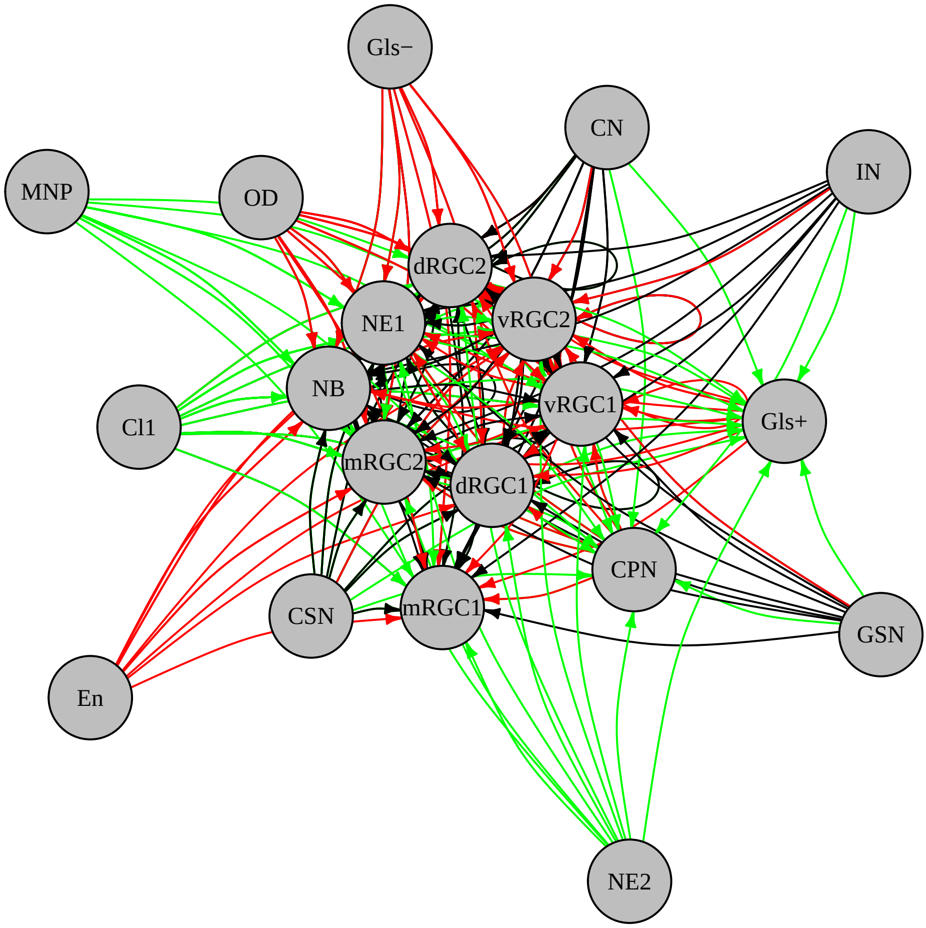

### SD_3L_IL4_vs_PBS_igf.pdf

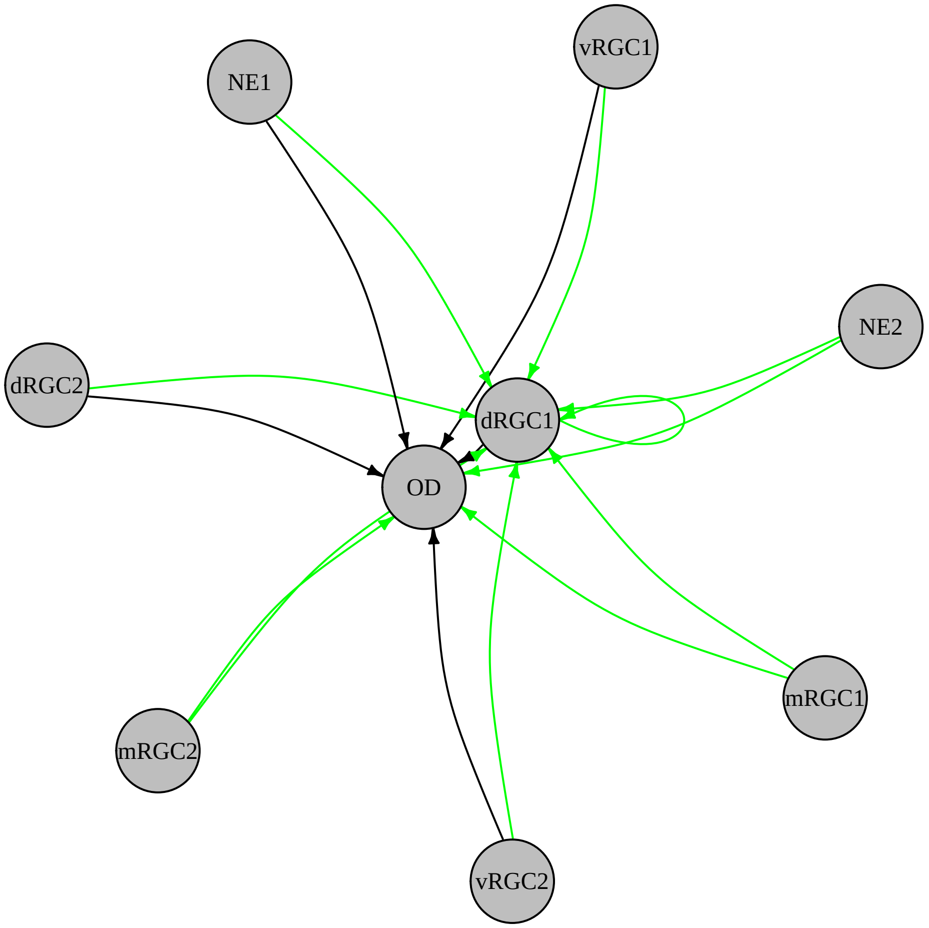

### SD_3L_IL4_vs_PBS_notch.pdf

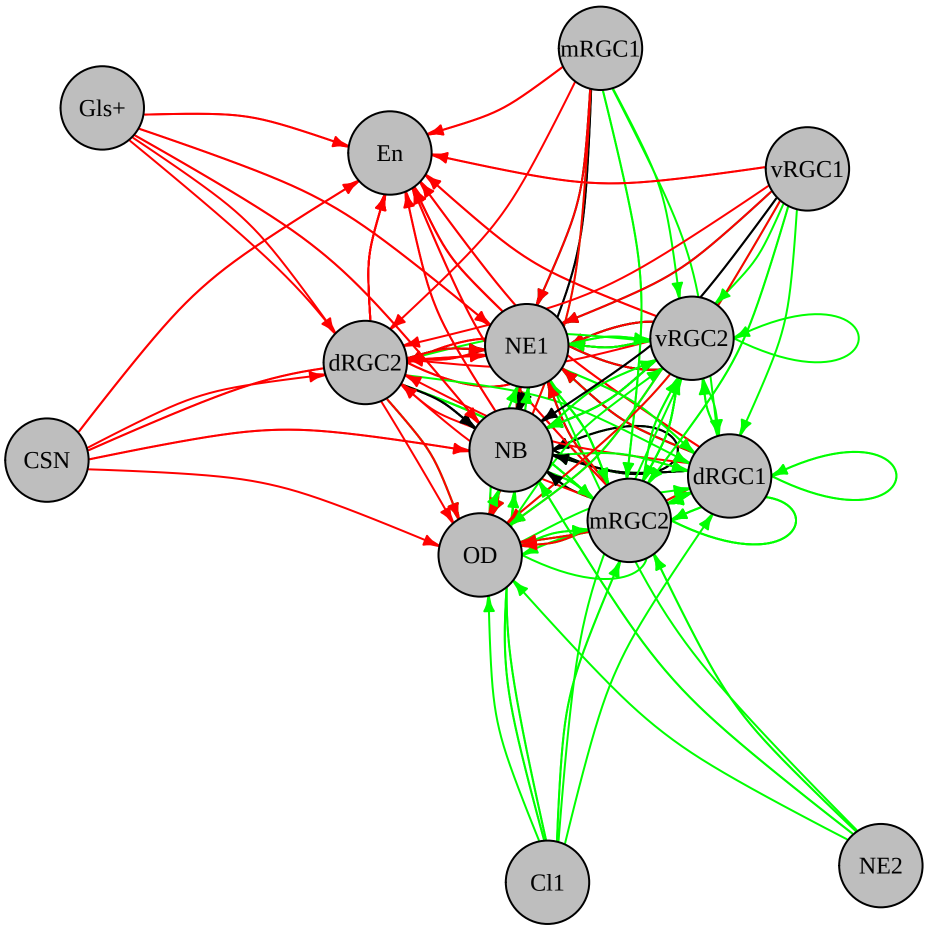

### SD_3L_IL4_vs_PBS_others_agrn.pdf

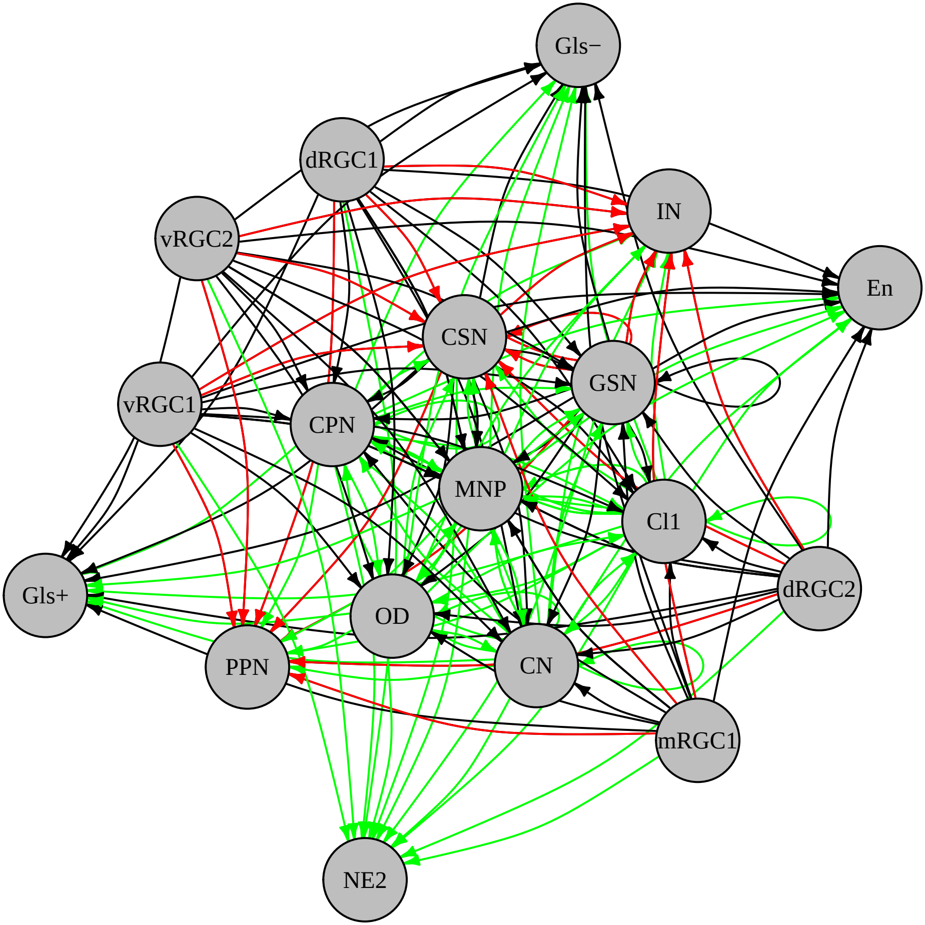

### SD_3L_IL4_vs_PBS_others_appa.pdf

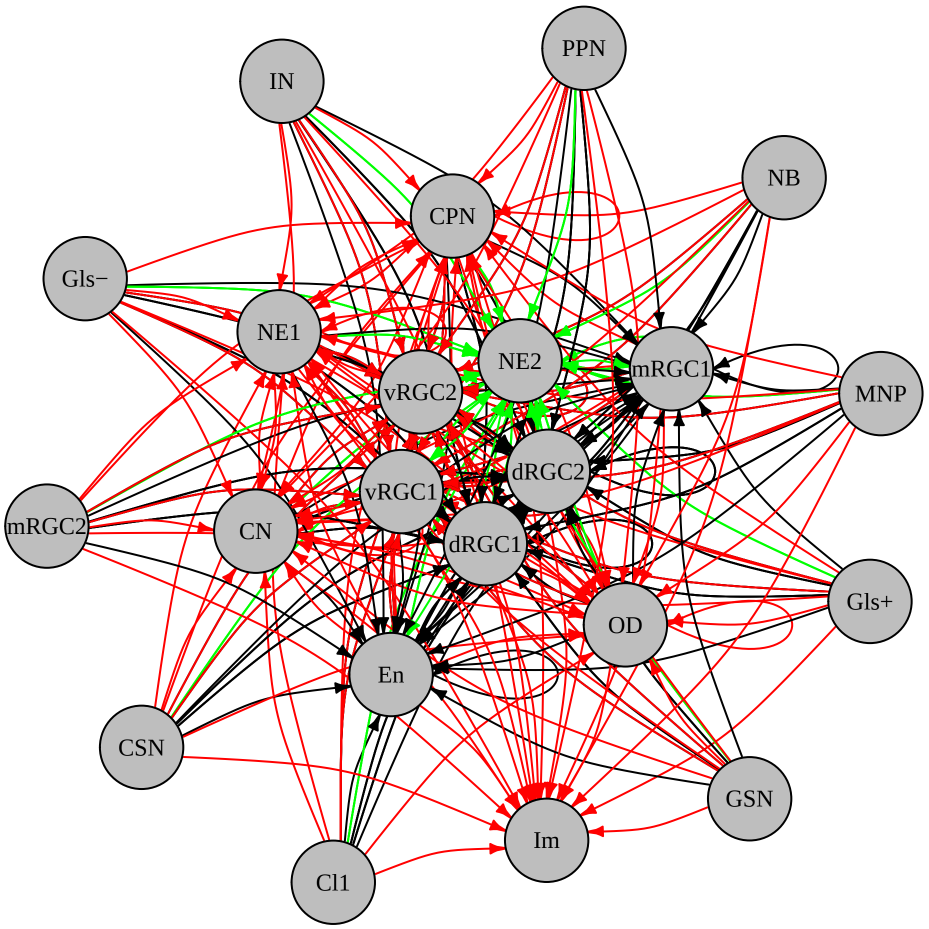

### SD_3L_IL4_vs_PBS_others_ctgfa.pdf

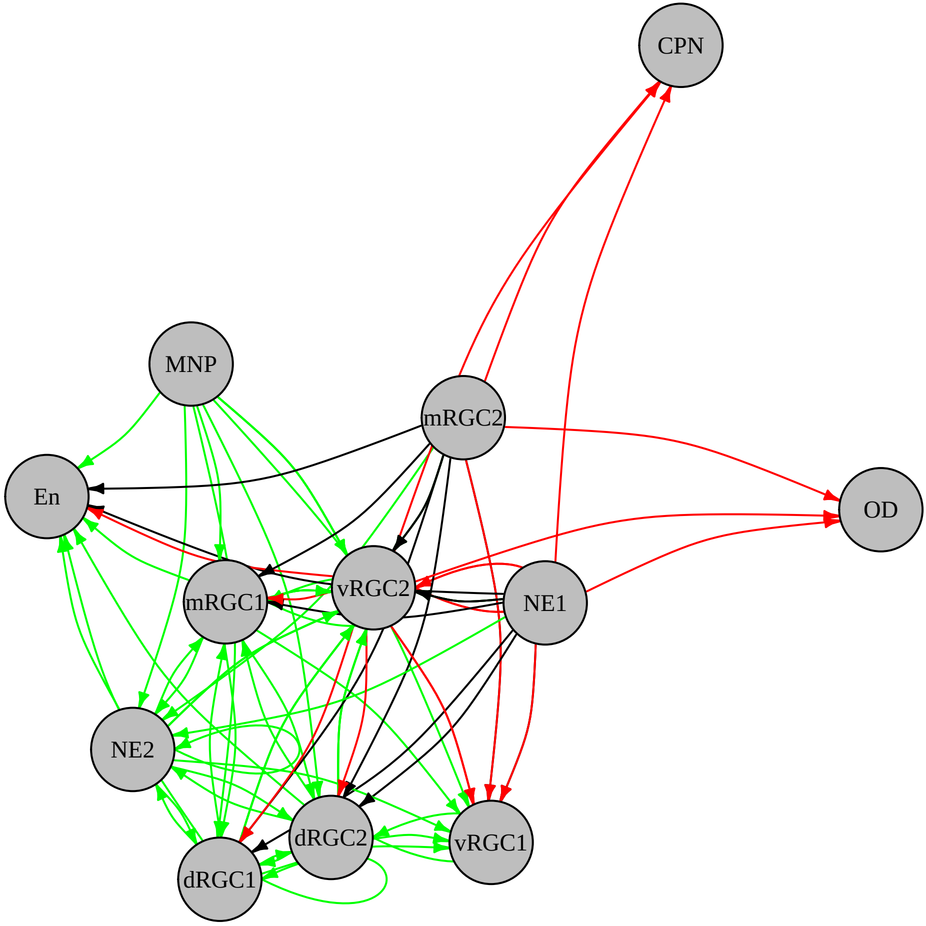

### SD_3L_IL4_vs_PBS_others_edil.pdf

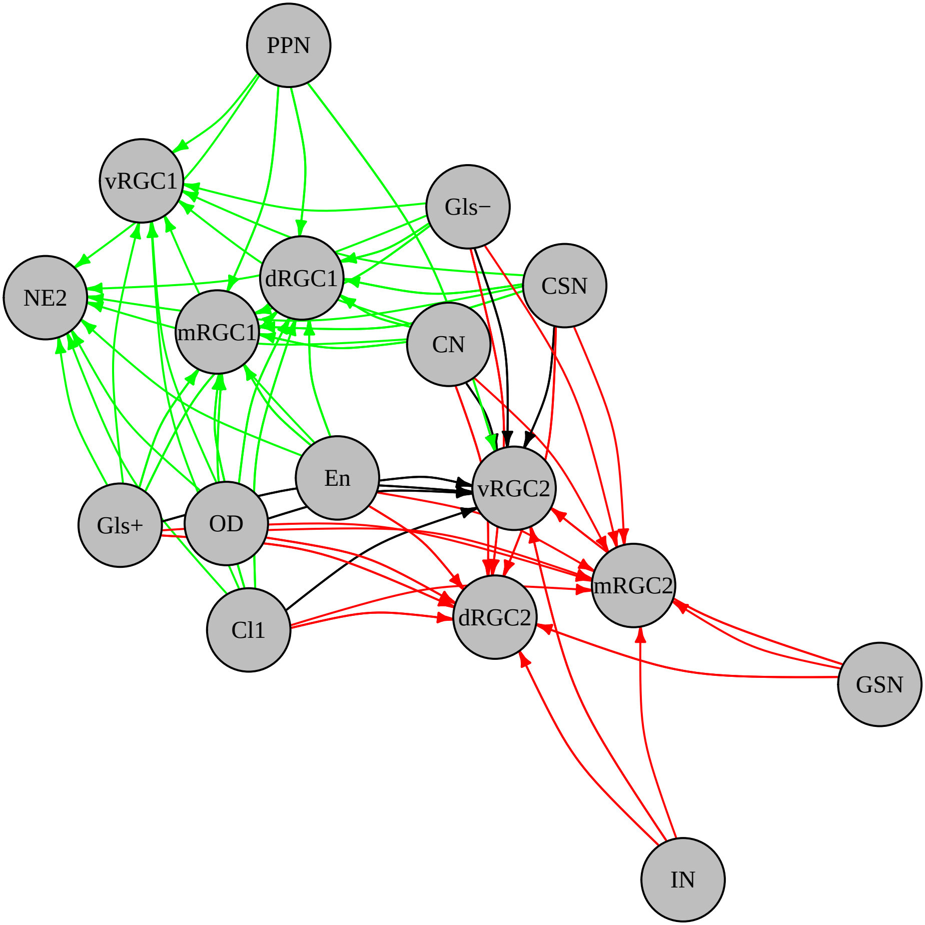

### SD_3L_IL4_vs_PBS_others_gnai.pdf

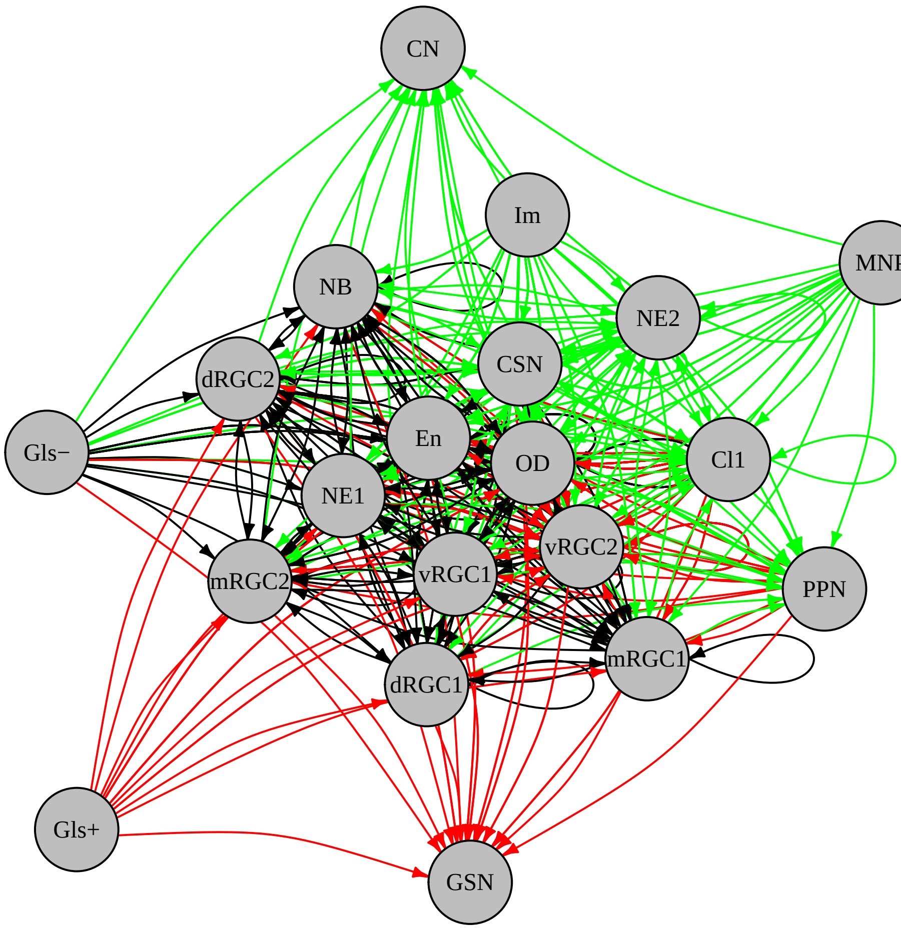

### SD_3L_IL4_vs_PBS_others_hbegfa.pdf

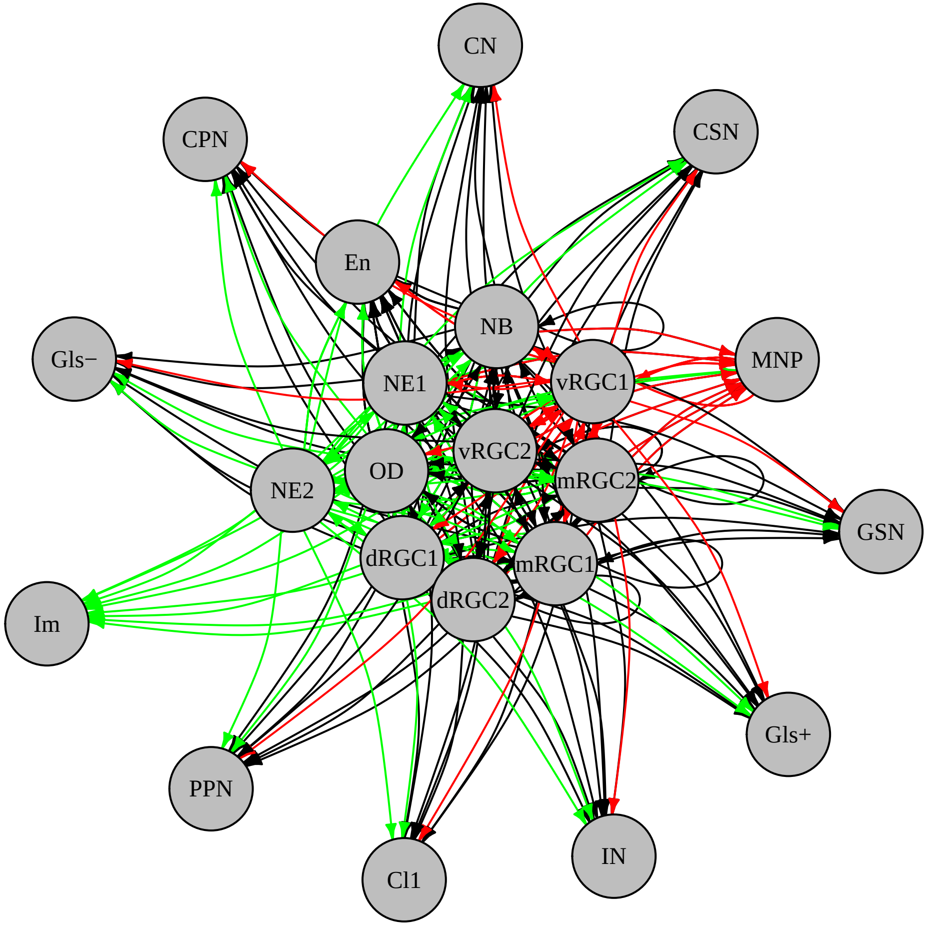

### SD_3L_IL4_vs_PBS_others_penk_ptn.pdf

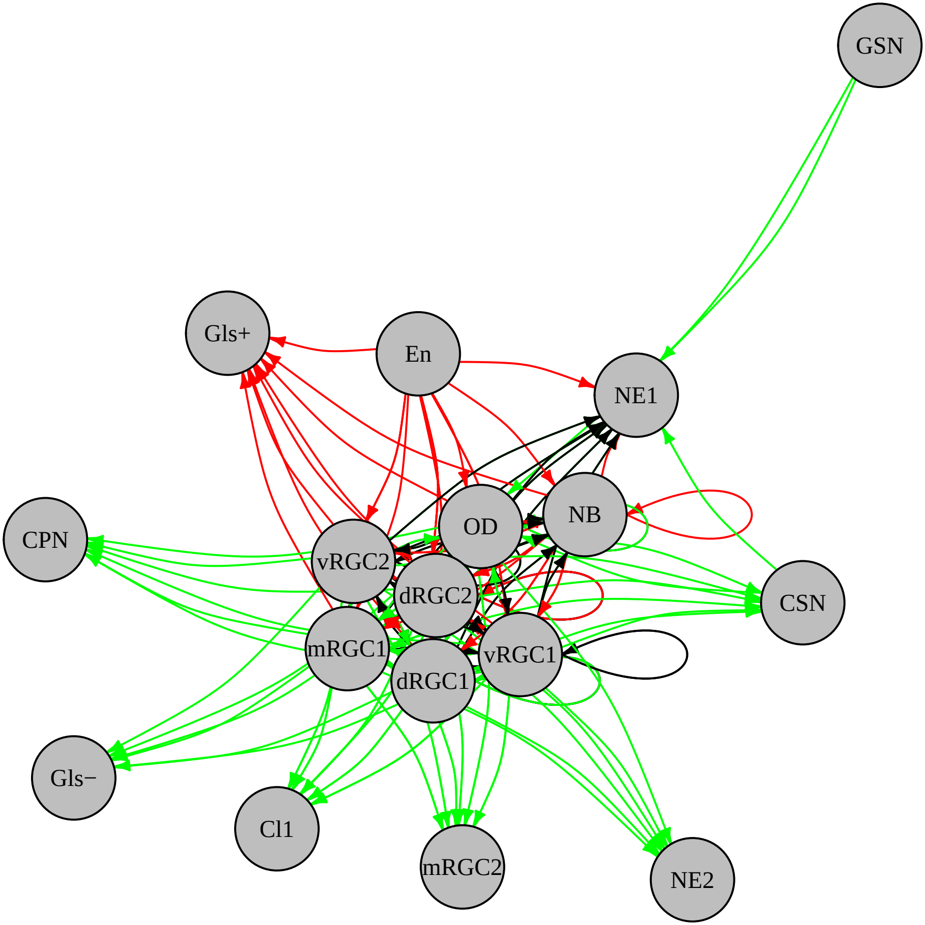

### SD_3L_IL4_vs_PBS_others_serpine.pdf

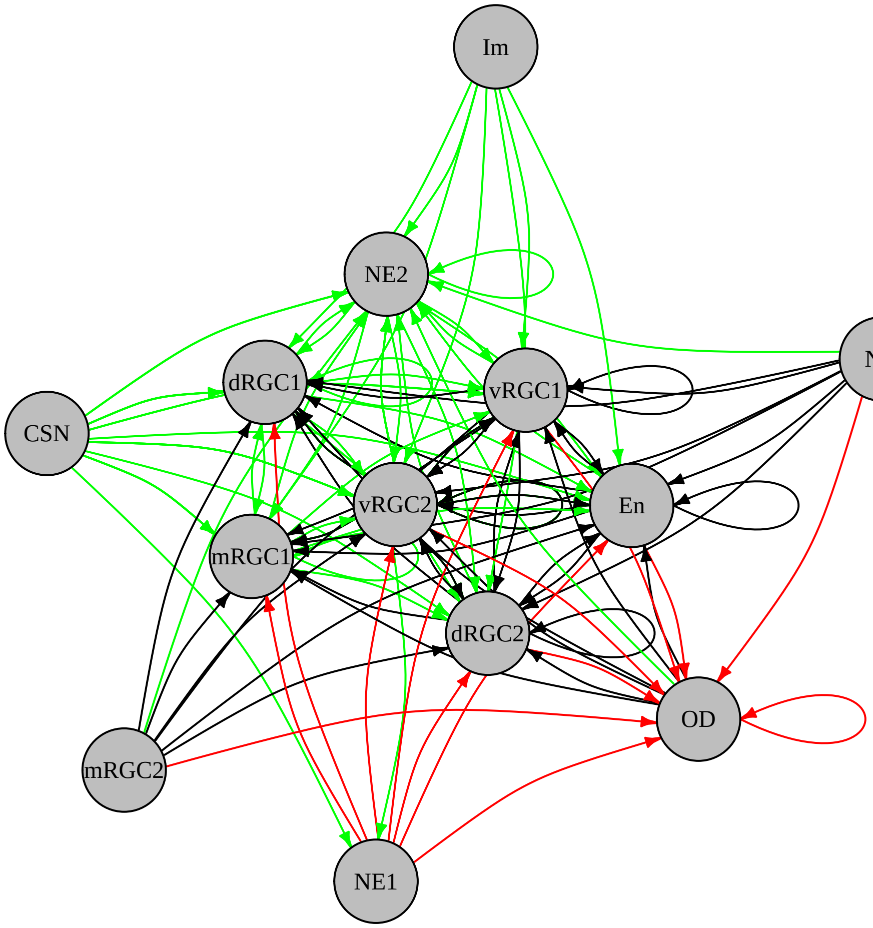

### SD_3L_IL4_vs_PBS_wnt.pdf

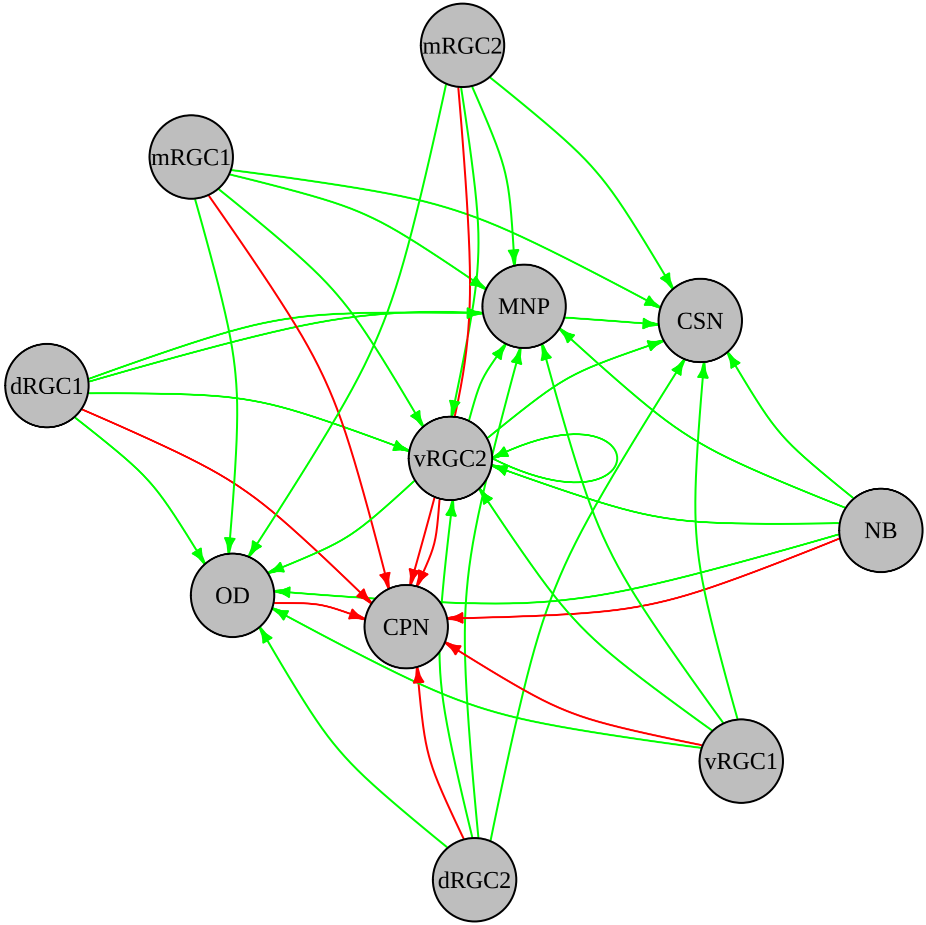

### Supplementary Materials

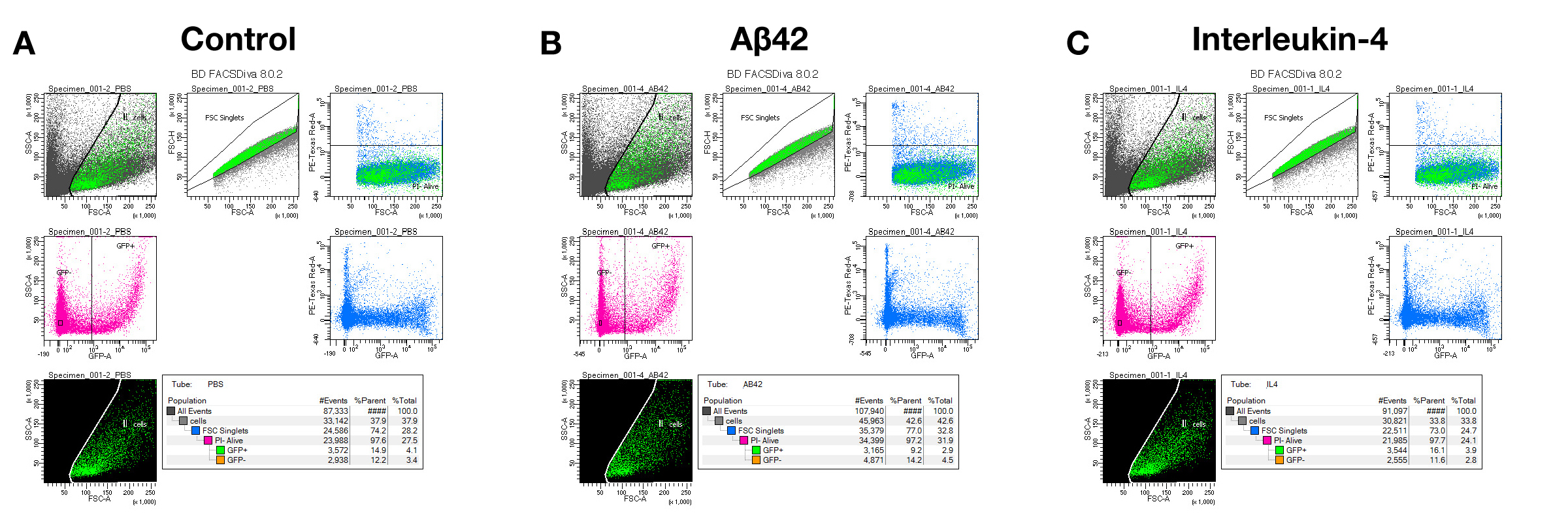

### Supplementary Materials

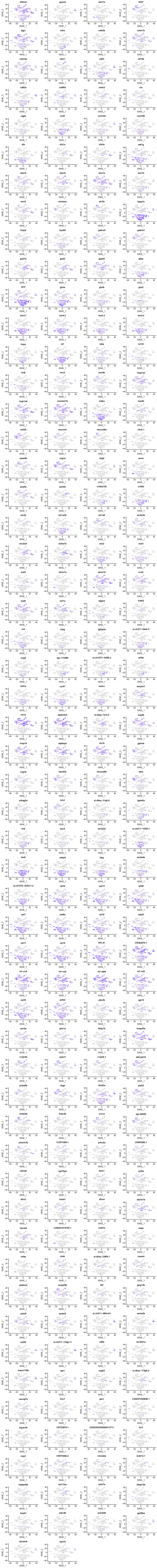

### Supplementary Materials

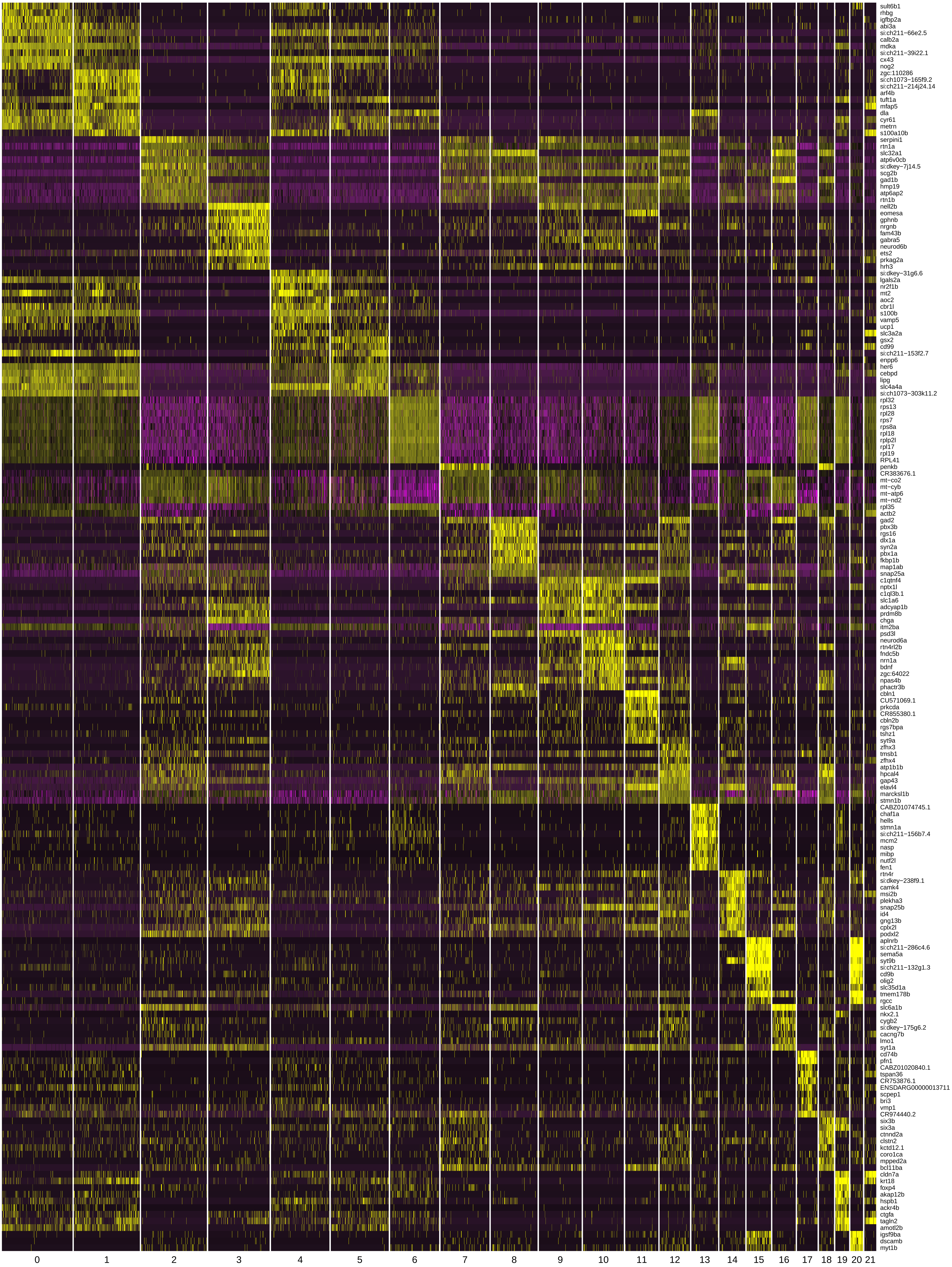
